# Molecular Mechanism of Action of Mitochondrial Therapeutic SS-31 (Elamipretide): Membrane Interactions and Effects on Surface Electrostatics

**DOI:** 10.1101/735001

**Authors:** Wayne Mitchell, Emily A. Ng, Jeffrey D. Tamucci, Kevin Boyd, Murugappan Sathappa, Adrian Coscia, Meixia Pan, Xianlin Han, Nicholas A. Eddy, Eric R. May, Hazel H. Szeto, Nathan N. Alder

## Abstract

Mitochondrial dysfunction includes heritable diseases, acquired pathologies, and age-related declines in health. Szeto-Schiller (SS) peptides comprise a class of amphipathic tetrapeptides that have demonstrated efficacy in treating a wide array of mitochondrial disorders, and are believed to target mitochondrial membranes due to their enrichment in the anionic phospholipid cardiolipin (CL). However, little is known regarding how SS peptides interact with or alter the physical properties of lipid bilayers. In this study, we have analyzed the interactions of the lead compound SS-31 (Elamipretide) with model and mitochondrial membranes using biophysical and computational approaches. Our results show that this polybasic peptide partitions into the membrane interfacial region with affinity and binding density that are directly related to surface charge. SS-31 binding does not destabilize lamellar bilayers even at the highest binding concentrations; however, it does cause saturable alterations in lipid packing. Most notably, SS-31 modulates the surface electrostatic properties of model and mitochondrial membranes, which could play a significant role in the mitoprotective properties of this compound. As a proof of concept, we show that SS-31 alters ion distribution at the membrane interface with implications for maintaining mitochondrial membranes subject to divalent cation (calcium) stress. Taken together, these results support a mechanism of action in which SS peptides interact with lipid bilayers and alter the biophysical (primarily electrostatic) properties of mitochondrial membranes as their primary mechanism of action. Understanding this molecular mechanism is key to the development of future compound variants with enhanced efficacy.

**Significance:** Szeto-Schiller (SS) peptides are among the most promising therapeutic compounds for mitochondrial dysfunction. However, the molecular target(s) and the mechanism of action of SS peptides are poorly understood. In this study, we evaluate the interaction of the lead compound SS-31 (Elamipretide) with mitochondrial and synthetic model membranes using a host of biophysical techniques. Our results show that SS-31 membrane interaction is driven largely by the negative surface charge of mitochondrial membranes and that SS-31 alters lipid bilayer properties, most notably electrostatics at the membrane interface. This work supports a mechanism in which SS peptides act on a key physical property of mitochondrial membranes rather than with a specific protein complex, consistent with the exceptionally broad therapeutic efficacy of these compounds.

## Introduction

Mitochondria are eukaryotic organelles that orchestrate the majority of cellular energy metabolism as well as a range of other processes including lipid biosynthesis, ion homeostasis, and programmed cell death (apoptosis). Mitochondrial dysfunction encompasses a wide array of both genetically encoded and acquired pathologies. Several mitochondrial diseases originate from heritable mutations in genes encoding mitochondrial proteins, either in nuclear or mitochondrial DNA (nDNA and mtDNA, respectively) (1). Moreover, reduced mitochondrial function accompanies the general decline in cellular bioenergetic capacity that occurs with aging as well as with many complex pathologies, including frailty, cardiomyopathy, cancer, and neurodegeneration (2). Features of mitochondrial dysfunction include gross alterations of the inner membrane cristae morphology, destabilization of respiratory complexes of the oxidative phosphorylation (OXPHOS) system, and overproduction of reactive oxygen and nitrogen species (ROS and RNS). Yet despite recent progress aimed at finding pharmacological approaches for mitochondrial disorders (3), there are presently no FDA-approved therapeutic compounds to treat them.

Szeto-Schiller (SS) peptides are among the most promising compounds currently under investigation for the treatment of mitochondrial dysfunction (4, 5). These synthetic tetrapeptides have a characteristic motif of alternating cationic and aromatic side chains (**Fig. 1**). Early in their development, SS peptides were shown to be cell-permeable in a wide range of cell types and to specifically target mitochondria. Despite their strong positive charge density (formal charge of +3), exogenously added SS peptides traverse the plasma membrane in an energy-independent and non-saturable manner, and accumulate strongly (1000 to 5000-fold) at the mitochondrial inner membrane (6, 7). The intrinsic therapeutic activity of the lead compound, SS-31 (**Fig. 1A**), was first demonstrated in cell culture studies, wherein the peptides were shown to protect cells against induced oxidative stress by curtailing oxidative cell death, reducing intracellular ROS, maintaining membrane potential (ΔΨ_m_), and preventing lipid peroxidation, all in a dose-dependent manner (6, 8). SS peptides have since been shown by many independent groups using cell culture and animal disease models to have significant efficacy in restoring mitochondrial function with a wide range of pathologies, including cardiomyopathy and heart failure, skeletal muscle injury/atrophy, ischemia and ischemia-reperfusion injury, kidney injury and disease, neurodegenerative diseases, cancer, and the heritable disease Friedreich’s ataxia (summarized in ref. 5). These studies have all underscored the safety and favorable pharmacokinetic profile of SS peptides. Currently, Stealth BioTherapeutics is conducting early to advanced-phase clinical trials with SS-31 (proprietary name Elamipretide) for a range of primary mitochondrial and aging-related diseases.

**Fig. 1.**
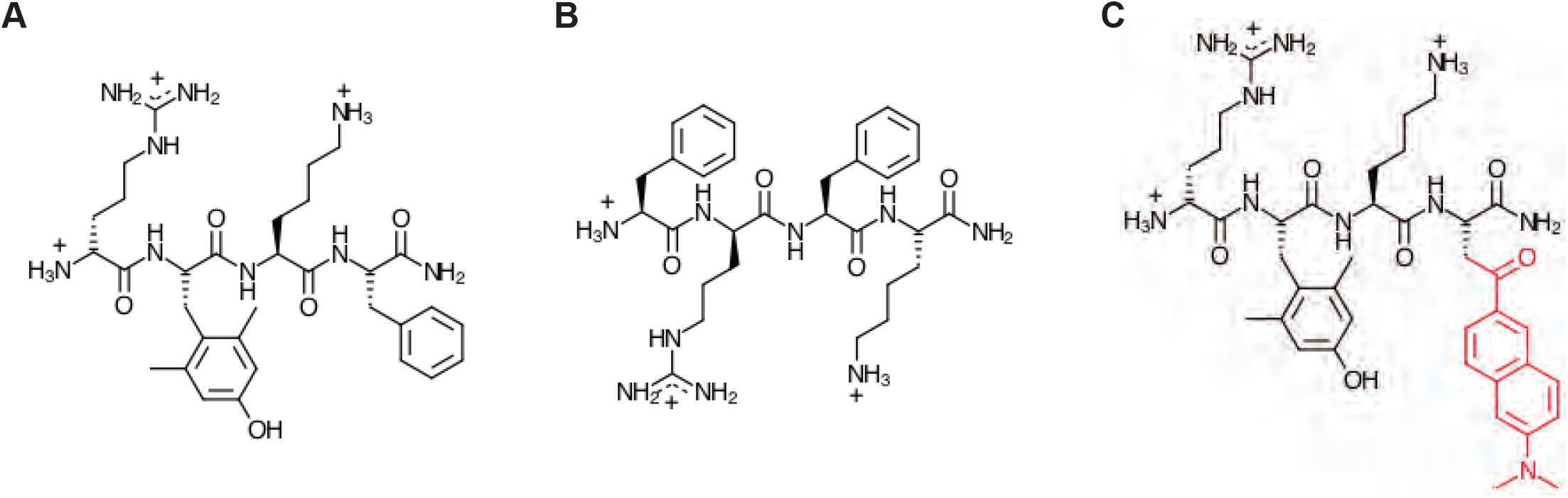
SS peptide chemical structures. (*A*) SS-31 (D-Arg-2’6’-dimethylTyr-Lys-Phe-NH_2_). (*B*) SS-20 (Phe-D-Arg-Phe-Lys-NH_2_). (*C*) [ald]SS-31, SS-31 with an aladan moiety (red) in place of Phe.

Despite abundant evidence for the broad therapeutic potential of SS peptides, the molecular mechanism of action (MOA) of these compounds is poorly understood. It was originally proposed that SS peptides act as mitochondria-targeted antioxidants. Peptide variants such as SS-31, which contain a free radical-scavenging tyrosine (or dimethyltyrosine) moiety, could in principle serve to reduce ROS burden (9). Yet antioxidant chemistry is not likely to be the primary mechanism of SS peptides, given: (i) that such scavenging would not occur catalytically, and (ii) that other peptide variants such as SS-20 (**Fig. 1B**), which do not contain scavenging side chains, have also proven effective in preclinical studies.

Perhaps the most critical insight into the MOA of SS peptides is that they are believed to target the mitochondrial inner membrane (MIM) by virtue of its enrichment in the anionic phospholipid cardiolipin (CL) (10–13). CL has an unusual dimeric structure with a two-phosphate headgroup and four acyl chains (**Fig. S1A**) and comprises roughly 10-20 mol% of the total lipids within the MIM. CL plays a central role in many mitochondrial processes, including apoptosis, mitophagy, signaling, fission/fusion dynamics, and energy metabolism (14). Within the protein-rich MIM, CL mediates many interactions with peripheral and integral membrane proteins (15, 16), underpinning its role in the assembly of OXPHOS complexes into supercomplexes (17). Moreover, with its unique physicochemical properties and structural polymorphism, CL has a complex phase behavior in lipid bilayers (18, 19). Thus, local enrichment of CL can promote negative curvature, which likely helps to stabilize cristae architecture (20, 21). Nascent CL is synthesized *de novo* within the MIM and subsequently undergoes a remodeling process to render mature species with a highly unsaturated complement of acyl chains that is species- and tissue-specific (22) (**Fig. S1B**). Alterations in CL distribution and biogenesis are associated with a number of diseases (23), including Barth syndrome (24), which is caused by defects in the transacylase tafazzin with concurrent aberrant remodeling of CL and buildup of the its lysolipid form, monolysocardiolipin (MLCL). Hence, being critical for mitochondrial structure and function, the anionic phospholipids of mitochondrial membranes are promising targets for therapeutic agents.

A primary mode of interaction between SS peptides and CL-rich lipid bilayers would be a truly unparalleled type of MOA. This is because the vast majority of drug compounds are designed to target and act upon specific proteins, not the lipid bilayer (with rare exceptions such as general anesthetics, whose mechanisms may involve alteration of bilayer properties (25)). In this study, we analyzed the nature of the SS-31 interaction with biomimetic model membranes and intact mitochondria to understand the forces that drive the peptide-membrane interaction and how SS-31 binding affects physical properties of model and natural lipid bilayers. Our results support a molecular MOA that involves alteration of bulk lipid bilayer properties, most notably the membrane surface electrostatic profile. We discuss multiple nonexclusive mechanisms by which the regulated tuning of surface electrostatics could underpin the broad therapeutic efficacy of SS peptides.

## Results

### Binding of SS-31 to model membranes

The interaction of basic amphipathic peptides with negatively charged membranes depends on the electric field originating from the bilayer surface as well as specific electrostatic and hydrophobic interactions between the peptide and lipids (**Fig. S2**). The first objective of this study was to quantitatively evaluate the binding of SS-31 to model membranes with specific lipid composition. For spectroscopic analysis of SS-31 binding, we made use of the endogenous spectral properties of the 2’,6’-dimethyltyrosine (2’,6’-Dmt) side chain at the second position in the peptide. Our initial spectral characterization of SS-31 confirmed the feasibility of using endogenous peptide fluorescence to quantify peptide-membrane interactions (**SI Results A,B** and **Figs. S3, S4**). We therefore proceeded to measure the binding of SS-31 to large unilamellar vesicles (LUVs) containing anionic lipids in a host background of the zwitterionic lipid palmitoyloleoyl-phosphatidylcholine (POPC, 16:0/18:1 PC). The anionic lipids included tetraoleoylcardiolipin (TOCL, all acyl chains 18:1), monolysocardiolipin (MLCL, all acyl chains 18:1), or palmitoleoyl-phosphatidylglycerol (POPG, 16:0/18:1 PG). Binding measurements were performed using two complementary approaches: (i) titration of fixed concentrations of LUVs with peptide, and (ii) titration of fixed concentrations of peptide with LUVs (**SI Results C** and **Fig. S5**). Note that in our analyses, we assume that peptide does not cross the bilayer of our model systems and can only access the outer leaflet; for LUVs of this size, the ‘effective lipid concentration’ of the outer leaflet ([L]^eff^) is taken as half of the total lipid concentration in the system (26).

Binding isotherms based on SS-31 emission intensity are shown in **Fig. 2**. First, peptide titrations were conducted by the progressive addition of SS-31 to LUVs of different composition at lipid concentrations ranging from 25-125 μM (**Fig. 2A**, see also **SI Results D and Figs. S6 and S7**). Second, lipid titrations were conducted by the progressive addition of LUVs to SS-31 at concentrations up to 15 μM (**Fig. 2B**). Taken together, these binding curves reveal the following: (i) in the absence of anionic lipid (LUVs with POPC only), SS-31 binding is negligible even at the highest lipid concentrations used, consistent with the established requirement of anionic lipids for binding (10–13); (ii) in the presence of anionic membranes, SS-31 binding displays saturation binding behavior; and (iii) membranes containing dianionic lipid (TOCL and MLCL) have a higher SS-31 binding capacity than membranes containing monoanionic POPG, reflected in the roughly twofold higher [SS-31] required to saturate binding in peptide titrations and in the roughly twofold lower [lipid] required to saturate binding in lipid titrations.

**Fig. 2.**
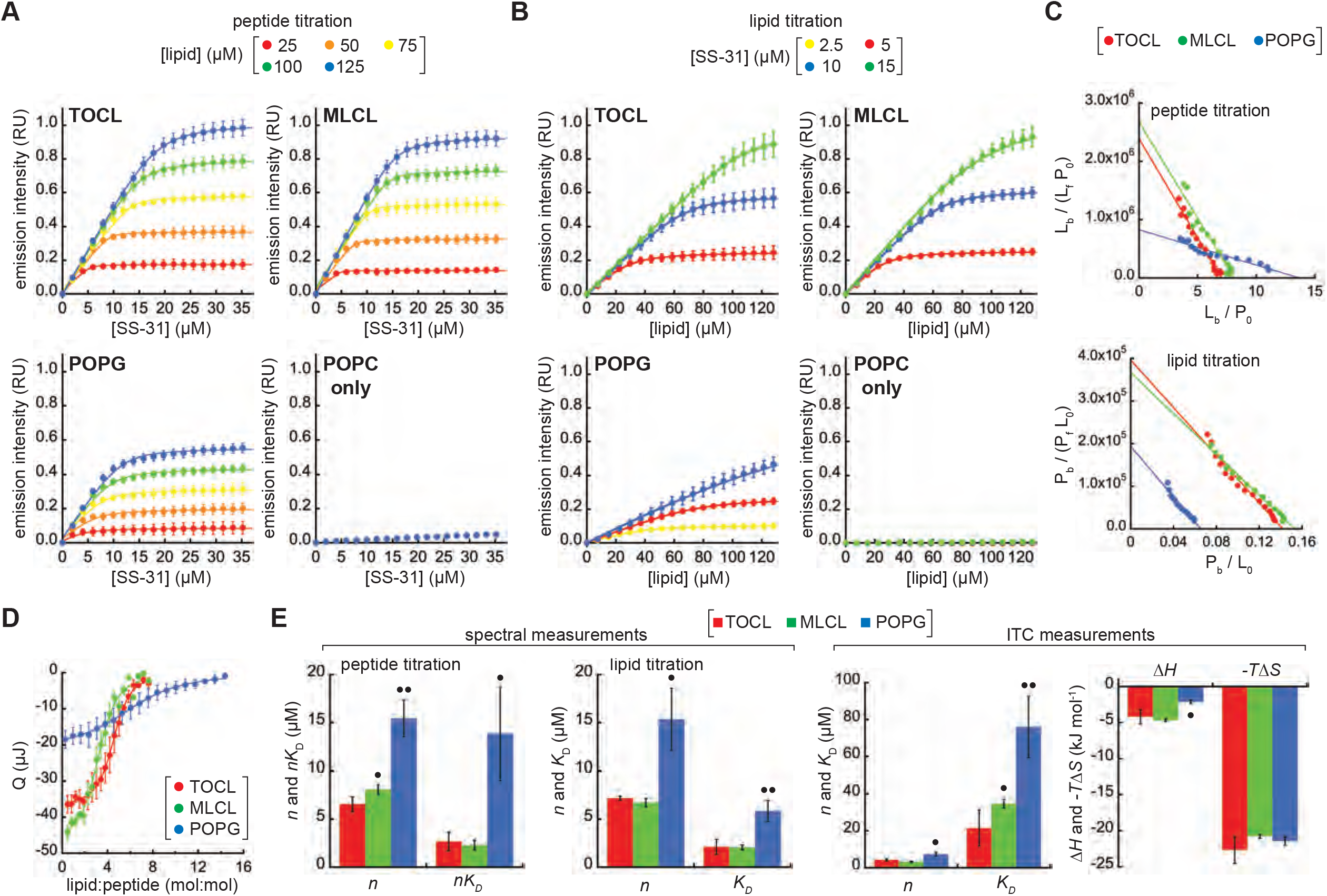
SS-31 binding isotherms. (*A-C*) Fluorescence-based measurements. (*A*) Peptide titrations. Liposomes containing anionic lipid (20 mol% TOCL, MLCL, or POPG) or POPC only at the lipid concentrations indicated were titrated with SS-31 (4 nmol increments). Values shown are means (n=3 ± SD) and lines show data fits based on Eq. S2. (*B*) Lipid titrations. Peptides at the concentrations indicated were titrated with liposomes (30 nmol lipid increments). Values shown are means (n=3 ± SD) and lines show data fits based on Eq. S3. (*C*) Scatchard analyses. Titration data with liposomes containing 20 mol% TOCL, MLCL, and POPG shown as Scatchard plots. *Upper panel*, peptide titrations (125 μM lipid) fit according to Eq. S6; *lower panel*, lipid titrations (10 μM SS-31, TOCL and MLCL; 5 μM SS-31, POPG) fit according to Eq. S8. (*D*) ITC analyses. Wiseman plot showing blank-subtracted average integrated heats (n=3 ± SD) as a function of lipid:SS-31 molar ratio for LUVs containing TOCL, MLCL, and POPG. (E) Summary of binding parameter data. Statistical differences within each dataset determined relative to TOCL-containing membranes: • *P* < 0.05; •• *P* < 0.01.

To obtain equilibrium binding parameters *n* (the number of lipid molecules per peptide bound) and *K*_D_ (the dissociation constant), we analyzed binding isotherm data by Scatchard analysis (**Fig. 2C**) and by fits of the data to Langmuir adsorption models (Eq. S2 and Eq. S3 for peptide and lipid titrations, respectively). The equilibrium binding parameters from this work (**Fig. 2E**, left panels) reveal two key points. First, for membranes containing cardiolipin variants, the average lipid:peptide stoichiometry (*n* = 6.9 and 7.4 for TOCL and MLCL, respectively) is roughly half the value for membranes with POPG (*n* = 15.4). Hence, if we assume ideal lipid mixing and approximate lipid cross-sectional areas (70 Å^2^ for POPC and POPG; 130 Å^2^ for TOCL and 110 Å^2^ MLCL (27, 28)), then for membranes composed of 20% TOCL or MLCL, the SS-31 “binding site” comprises an area of ~560-580 Å^2^ in which each peptide associates, on average, with 1.4-1.5 dianionic lipids. By comparison, for membranes containing 20% POPG, SS-31 binds an area of roughly 1080 Å^2^ and associates on average with 3 monoanionic lipids. Based on these stoichiometric relationships, then, there is an approximate charge balance between each peptide and its corresponding lipid binding site. Second, the affinity of SS-31 for membranes containing TOCL and MLCL is significantly higher than it is for membranes containing POPG. Specifically, the average dissociation constants for lipid monomers (reflected in *K*_D_ values for lipid titrations and in *nK*_D_ values for peptide titrations) are 2.0-2.9 μM for membranes composed of 20% TOCL and MLCL, whereas they range from 6.0-13 μM for membranes composed of 20% POPG. These values of *n* and *K*_D_ are consistent with parameters measured for other amphipathic membrane-active peptides (29, 30). As an independent measure of peptide binding, the molar partition coefficients (*K*_P_, Eq. S9) for membranes containing 20% TOCL (*K*_P_ = 5.07 ± 0.76 x10^4^) and 20% MLCL (*K*_P_ = 5.90 ± 0.44 x 10^4^) are significantly higher than that of 20% POPG-containing membranes (*K*_P_ = 1.28 ± 0.63 x 10^4^).

As a complementary approach for measuring SS-31 membrane interactions, we performed isothermal titration calorimetry (ITC) measurements by the progressive injection of LUVs into solutions of peptide in the sample cell (31). The interaction of SS-31 with membranes containing anionic lipid was shown to be exothermic in all cases (**Fig. S8**), and fits of complete binding isotherms (**Fig. 2D**) yielded the thermodynamic binding parameters shown (**Fig. 2E**, right panels). The values of *n* and *K*_D_ from fits to the ITC data showed consistent differences between TOCL/MLCL- and POPG-containing membranes when compared with our spectral-based measurements (Fig. 2). However, in comparison with our spectral analyses, there were differences in the absolute values from ITC-derived parameters (mean *n* values slightly lower and *K*_D_ values ~2-3 times higher). This difference could be related to different phenomena being measured: our spectral analyses may only reflect the binding event *per se*, whereas ITC measurements could be reflecting binding as well as peptide-dependent alterations of bilayer properties. Apropos of this point, we did observe a slight asymmetry in our exothermic peaks, particularly for TOCL- and MLCL-containing membranes, which could be related to thermotropic alterations in the bilayers following peptide interaction.

Comparison of the binding-associated changes in enthalpy (Δ*H*) and entropy (Δ*S*) provide insights into the membrane binding mechanism. Namely, exothermic (Δ*H*<0) values of peptide-lipid interaction originate mainly from the establishment of polar contacts; by contrast, increases in the system entropy (Δ*S*>0) originate mainly from the burial of hydrophobic side chains in the acyl chain region with concurrent release of water and ions from nonpolar surfaces of the peptide and bilayer (32). For membranes containing TOCL, MLCL, and POPG, the interaction of SS-31 was associated with a favorable enthalpic change (Δ*H* = −4.2, −4.7, and −2.1 kJ mol^−1^, respectively); the twofold difference in Δ*H* between TOCL/MLCL- and POPG-containing membranes is consistent with the difference in formal charges of these anionic lipids. The peptide-bilayer association, however, was predominantly entropy-driven (−*T*Δ*S* = −22.7, −20.8, and −21.5 kJ mol^−1^, respectively), showing that peptide association is stabilized mostly by burial of the nonpolar residues. Given that the hydrophobic contribution to peptide binding energy is proportional to the exposed side chain area (~80 J mol^−1^ Å^−2^) (33) the burial of Tyr (similar to 2’,6’-Dmt) and Phe side chains (accessible surface areas 187 and 175 Å^2^, respectively) would theoretically contribute a combined ~28 kJ mol^−1^, in reasonable agreement with our measurements. Moreover, because SS peptides are short and cannot form any appreciable secondary structure upon membrane binding, their membrane binding does not likely incur an entropic penalty that would be associated with stabilizing a secondary structure.

Taken together, our spectral and calorimetric binding data show that the bilayer binding density and affinity of SS-31 are directly related to the anionic lipid composition, that the lipid-dependent differences in interaction energy are mostly on the net charge of anionic lipids, and that membrane interactions are enthalpically and entropically favorable.

### Dependence of SS-31 binding on model membrane surface electrostatics

The electrostatic profile of biomembranes consists of multiple potential energy functions (**Fig. S2**). Among them, the surface potential (ψ_s_) originates from charge distribution at the membrane interface based on ionizable functional groups (e.g., lipid headgroups) and adsorbed ions, creating a strong electrostatic driving force for the binding of charged moieties to the bilayer surface (34). Electrolytes partitioned at the bilayer interface can have a complex effect on membrane interactions of amphipathic peptides. By attenuating the ψ_s_, solution cations can reduce the electrostatic attraction of basic peptides to the anionic lipid surface; however, increasing solution ionic strength may also favor the burial of hydrophobic side chains in the nonpolar core of the bilayer. We therefore explored the relationship between SS-31 binding and the ψ_s_ of model membranes and the influence of ionic strength (**Fig. 3**).

**Fig. 3.**
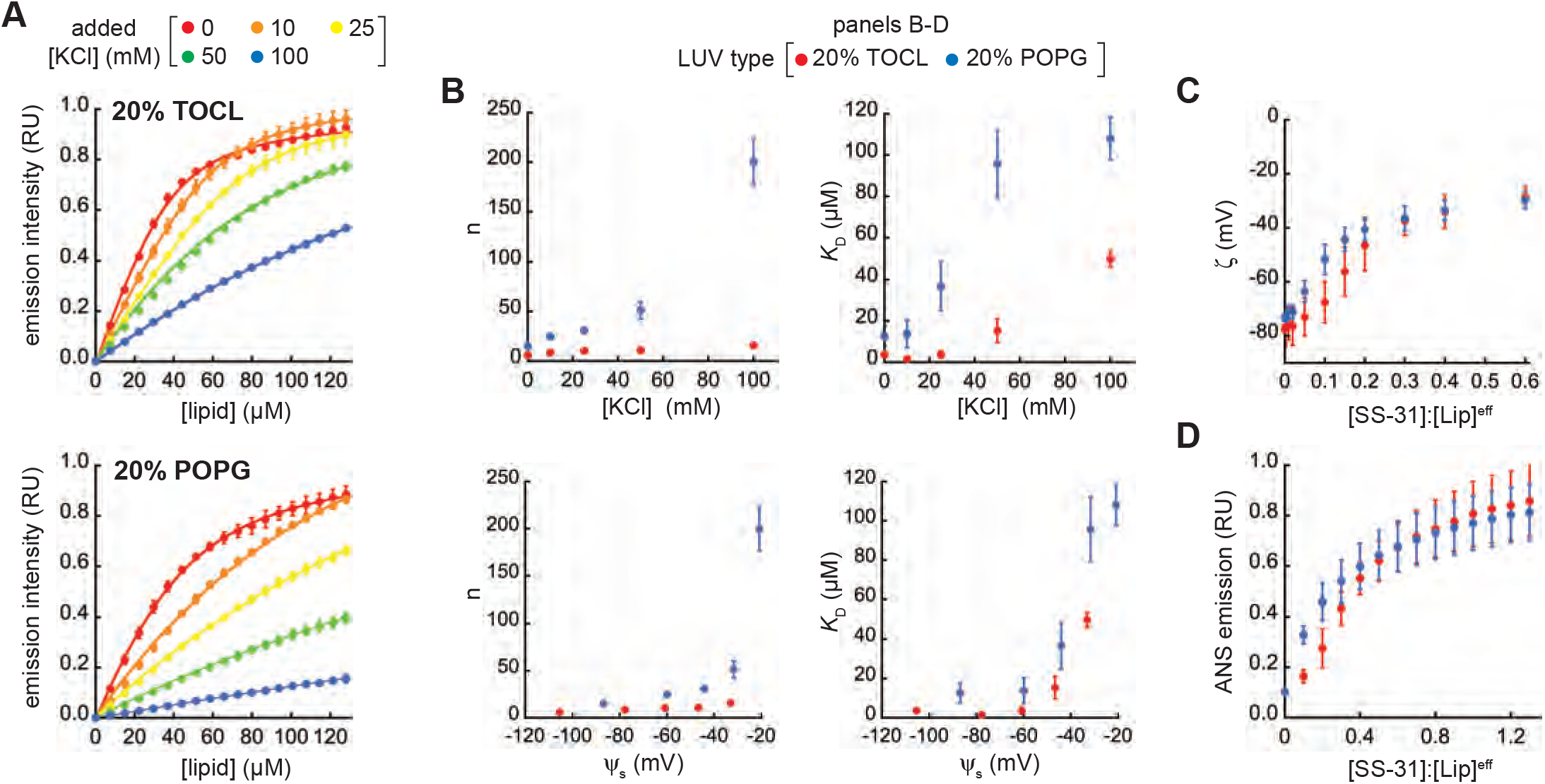
SS-31 binding and surface electrostatics. (*A*) Binding isotherms. Peptide was titrated with LUVs of the indicated composition (20% TOCL, [SS-31] = 7.5 μM and 20% POPG, [SS-31] = 3.8 μM) at 30 nmol lipid increments in the presence of different concentrations of added KCl. Values shown are means (n=3 ± SD) and lines show data fits based on Eq. S3. Because isotherms at higher [KCl] did not reach saturation, curves within each membrane set were fit globally assuming a common saturation point. (*B*) Equilibrium binding parameters *n* and *K*_D_ at different ionic strengths as a function of [KCl] (*upper panels*) and as a function of ψ_s_, calculated from GCS formalism, Eq. S10-S12 (*lower panels*) based on binding isotherms of panel A. (*C,D*) Effect of SS-31 binding on surface charge. Measurements of (*C*) zeta potential (ζ) and (*D*) ANS fluorescence shown as a function of [SS-31]:[Lip]^eff^ for membranes containing 20% TOCL and 20% POPG as indicated (values are means, n=3 ± SD).

SS-31 binding curves measured by titration of SS-31 with LUVs containing 20% TOCL or POPG revealed a strong effect of monovalent electrolyte (**Fig. 3A**). We confirmed that SS-31 binding to model bilayers was reversible in a manner that is dependent upon solution ionic strength used for these binding isotherms (**SI Results E** and **Fig. S9A,B**). When binding parameters *n* and *K*_D_ calculated from the curves of Fig. 3A are considered as a function of salt concentration, there is a large disparity between membranes containing divalent TOCL and monovalent POPG (**Fig. 3B**, *upper panels*). However, recasting these parameters as a function of ψ_s_, calculated from Gouy-Chapman-Stern (GCS) formalism (35, 36) (Eq. S10-S12), reveals a much smaller difference between membranes containing the two anionic lipids (**Fig. 3B**, *lower panels*). This GCS analysis, which accounts for the roughly twofold larger intrinsic surface charge (σ^max^) of TOCL-*vs*. POPG-containing bilayers, indicates that SS-31 binding is more related to ψ_s_ *per se*, rather than to any specific features of CL.

To continue this analysis, we determined the relationship between SS-31 binding and surface electrostatics by two complementary approaches. We first performed zeta potential (ζ) measurements of LUVs containing 20 mol% TOCL or POPG with increasing [SS-31] (**Fig. 3C**). The ζ represents the electrostatic potential at the hydrodynamic shear plane of membranes, and is related to the ψ_s_ by the Debye constant (κ), which describes the position-dependent decay of electrostatic potential from the membrane surface to the bulk solution. As expected for the binding of a polybasic peptide to a negatively charged surface, SS-31 caused a saturable reduction in the magnitude of ζ for both anionic bilayers. As an independent technique, we measured the effect of SS-31 on surface electrostatics using 1,8-ANS, a fluorescent reporter whose binding to anionic bilayers increases as the magnitude of the ψ_s_ is reduced, measured as an increase in probe emission intensity (37) (**Fig. 3D** and **Fig. S9C**). Consistent with our ζ data, SS-31 caused a saturable 1,8-ANS-detected reduction in the ψ_s_ of anionic model membranes. We observed the same response with SS-20 (**Fig. S9D**), supporting that this effect on membrane surface electrostatics is a general feature of SS peptides. Beyond the observation that SS peptides attenuate membrane surface potential, two features are notable from these experiments. First, for both ζ and 1,8-ANS measurements, SS-31 elicited a hyperbolic decay of ψ_s_ for PG-containing bilayers, whereas the response was more sigmoidal for CL-containing bilayers (Fig. S9C,D blue vs. red). This suggests that at low peptide concentration, there is not a linear correspondence between peptide binding and charge attenuation for CL-containing membranes. Second, the ζ responses, which provide an absolute measure of ψ_s_, consistently saturated near −30 mV with increasing peptide. This indicates that when the model bilayers are maximally bound with peptide, there remains an appreciable negative surface charge density. This observation is in contrast to other basic peptides (38) and multivalent cations (39) that cause charge overcompensation (lead to positive surface charge density) upon binding anionic bilayers at high concentration. Two nonexclusive explanations may account for this: (i) SS-31 binding may render some ionized phosphate groups unavailable for further interaction, and/or (ii) the formal charge on SS-31 may become reduced upon binding. Regardless of the mechanism, the fact that SS-31 attenuates the ψ_s_, but does not completely reverse it, is a fundamental feature of its effect on the membrane surface electrostatic profile.

### SS-31 maintains the lamellarity of model membranes but alters lipid interactions at the interface

Amphipathic molecules such as antimicrobial peptides can induce large structural changes in membranes, including micellization, pore formation, and induction of inverted topologies (40). We therefore addressed whether SS-31 induced structural polymorphism within model bilayers using a range of approaches (**Fig. 4**). Based on synchrotron small angle x-ray scattering (SAXS) measurements, LUVs composed of POPC alone or with 20% TOCL, MLCL, or POPG yielded scattering profiles typical of lipid vesicles (41), and the presence of SS-31 did not cause structural perturbation (**Fig. 4A, SI Results F**, and **Fig. S10A**). Similarly, based on ^31^P solid state NMR (ssNMR) measurements, these model membranes yielded lineshapes consistent with lamellar bilayers (42), and we observed no effect of SS-31 on bilayer structure (**Fig. 4B, SI Results F**, and **Fig. S10B**). We conclude that even at the highest peptide concentration used in this study ([P]:[L]^eff^ = 1:5), SS-31 did not cause major topological alterations of our model membranes. Thus, these membranes are expected to exist stably in the liquid crystalline lamellar mesophase over all peptide concentrations.

**Fig. 4.**
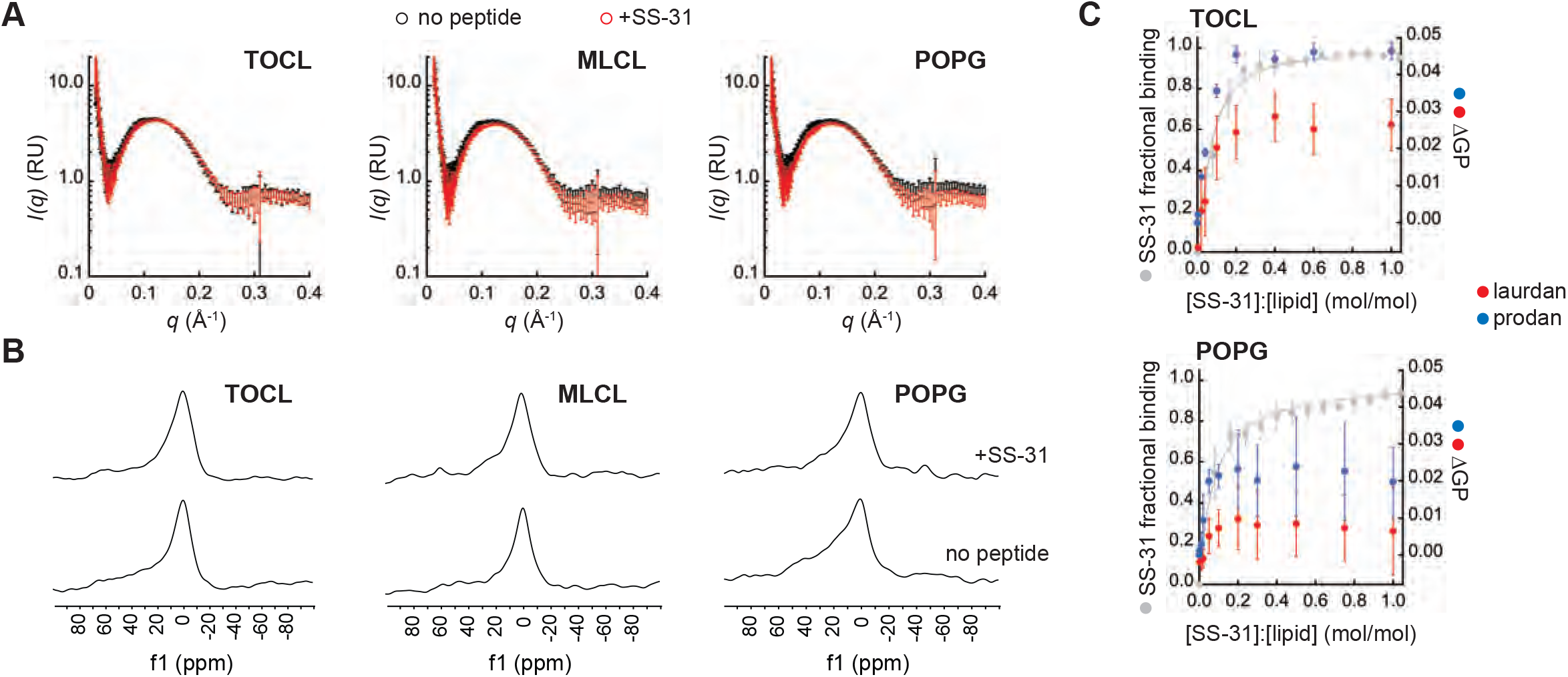
Effects of SS-31 on structural properties of model membranes. (*A,B*) SS-31 does not cause major structural perturbation of model membranes. LUVs composed of 20% anionic lipid (TOCL, MLCL, or POPG) in POPC background were measured in the absence or presence of peptide ([P]:[L]^eff^ = 1:5) as indicated. (*A*) Synchrotron SAXS measurements showing background-subtracted scattering density profiles for LUVs in the absence (black) or presence (red) of SS-31. (*B*) ^31^P solid state NMR spectra of LUVs in the absence (*lower spectra*) or presence (*upper spectra*) of SS-31. (*C*) SS-31 alters lipid packing interactions. Changes in laurdan/prodan GP values (ΔGP) (n=3 ± SD) with increasing [SS-31] for LUVs containing 20% TOCL or POPG in POPC background as indicated. Data and fits in grey show SS-31 fractional saturation (calculated from Fig. 2A, 25 μM lipid).

We therefore addressed whether SS-31 altered lipid interactions within these model bilayers. To this end, we used membrane-bound fluorescent reporters of lipid dynamics and packing that partition into the bilayer at different depths (**Fig. S10C**). Using DPH anisotropy (<*r*>^DPH^) as a readout of the fluid dynamics of hydrocarbon tails, we observed no effect of SS-31 in TOCL- or POPG-containing membranes (**Figs. S10D,E**). We then used solvatochromic probes laurdan and prodan, whose spectral properties are quantified as the generalized polarization (GP^LAU^ and GP^pro^, respectively) that increases with enhanced lipid packing (reduced interfacial hydration) (43) (**SI Results F and Fig S10F**). We observed modest, yet repeatable and saturable increases in GP^LAU^ and GP^PRO^ that corresponded to the fractional occupancy of peptide on the membrane (**Fig. 4C**). Notably, the response of GP values in TOCL-containing bilayers was much higher in magnitude compared to the response in those with POPG. Based on these results, SS-31 binding causes a change in the hydration/polarity of the bilayer interface that likely results from an increase in lipid headgroup packing density.

### Molecular dynamics analysis of the interaction between SS-31 and lipid bilayers

We next used all-atom molecular dynamics (MD) simulations to analyze the interaction between SS-31 and lipid bilayers (**Fig. 5, SI Results G, and Figs. S11-S14**). These simulation systems contained solvated bilayers composed of 20 mol% TOCL, MLCL, or POPG in a host background of POPC, and were conducted in the presence or absence of SS-31 to evaluate SS-31-bilayer interactions and how the presence of SS-31 may affect bilayer properties (**Fig. 5A**). In our initial investigations, we performed spontaneous membrane binding simulations by placing ten SS-31 peptides in the aqueous phase in random orientations at distances of 1-3 nm from the bilayer surface and conducting simulations for up to 1.6 μs. Throughout each trajectory, we quantified the distance between each side chain and the bilayer center of mass along an axis perpendicular to the membrane (the z-coordinate, **Fig. S11** and **Fig. 5B**). We observed rapid association (<200 ns) of all peptides to the bilayer surface, in which all side chains had docked to the membrane near the interfacial region, presumably driven by the electrostatic attraction between the polybasic SS-31 and the negative surface charge of the lipid bilayer. However, the membrane association time differed among the side chains. For example, the N-terminal Arg adsorbed to the bilayer rapidly (within 50 ns), whereas the C-terminal Phe took significantly longer to localize to the bilayer (within 180 ns) (**Fig. 5B**).

**Fig. 5.**
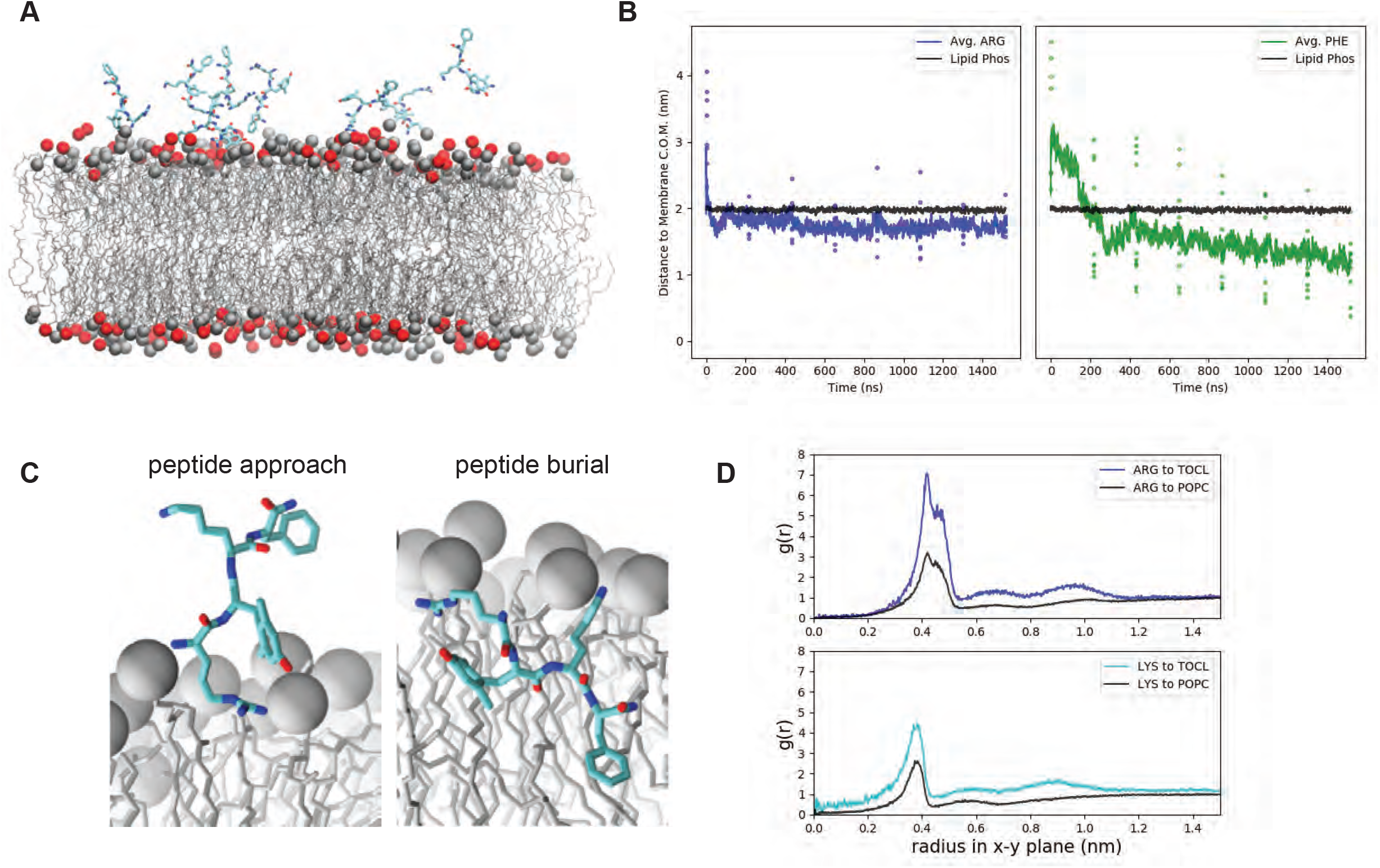
Molecular dynamics simulations of SS-31 bilayer interactions. (*A*) Snapshot of a typical MD simulation with SS-31 peptides in the aqueous phase relative to the upper leaflet of a 20% TOCL bilayer. Peptides are shown in licorice representation (cyan, carbon; blue, nitrogen; red, oxygen), lipid acyl chains are in wireframe, and lipid phosphates are shown as van der Waals spheres for POPC (gray) and TOCL (red). (*B*) Temporal evolution of SS-31 Arg (blue) and Phe (green) side chain positions relative to the membrane center of mass (C.O.M.). Black lines show the average position of all phosphates in the upper leaflet, colored dots show z coordinates (z^POS^) of individual side chains, and colored lines represent average z^POS^ values. (*C*) Orientation of SS-31 bound to a lipid bilayer consisting of 20 mol% TOCL in a POPC background in the peptide approach (*left*) and buried (*right*) states. Peptide representation as in (*A*), with all lipid phosphates shown as gray van der Waals spheres. (*D*) Lipid distributions around basic SS-31 residues. Lateral (x-y plane) radial distribution profiles from Arg and Lys side chains to lipid headgroup phosphate atoms of TOCL or POPC as indicated.

From these simulations, we observed two main types of SS-31 poses on the membrane (**Fig. 5C**): a *peptide approach* state in which the primary points of contact with the membrane are the N-terminus and Arg side chain anchored to phosphate headgroups, and a *peptide burial* state in which the aromatic side chains have partitioned into the nonpolar core while the basic side chains remain electrostatically tethered to headgroup phosphates. Notably, the burial of the aromatic side chains of SS-31 in the acyl chain region is consistent with previous NMR studies (10). Once the peptides reached the burial state, each side chain assumed a distinct, stable membrane insertion depth, quantified as the average z-coordinate between the side chain and the membrane center of mass (z^pos^) (**Fig. S11B,C**). In the 20:80 TOCL:POPC system, the basic side chains occupied the interfacial region, with Arg (z^pos Arg^ = 1.75 nm) residing just below the average position of lipid phosphates (z^pos Phos^ = 1.98 nm) and Lys residing at a slightly more distal position near the headgroups (z^pos Lys^ = 1.99 nm). By comparison, the aromatic residues assumed stable positions within the nonpolar core with the Phe side chain burying deeper (z^pos^ ^Phe^ = 1.12 nm) than the 2’,6’-Dmt side chain (z^pos DmT^ = 1.37 nm). These residue-specific membrane insertion depths were consistent across the three lipid compositions tested.

To evaluate the association of SS-31 side chains with lipid headgroups, we analyzed radial distribution profiles from our MD trajectories (**Fig. 5D, Fig. S12A**). This analysis revealed that basic side chains of SS-31 preferentially interacted with anionic lipids. For example, based on g(r) peak height, in lipid systems with 20% TOCL, Arg and Lys showed 2.25- and 1.72-fold relative increases in local concentration of phosphate groups from TOCL compared with those from POPC (**Fig. 5C**). These results are consistent with analysis of peptide-lipid co-diffusion, suggesting that the SS-31 Arg can concurrently associate with multiple lipid phosphates with a preference for TOCL over POPC (**Fig. S12B**). As a complement to these analyses, we evaluated how the presence of SS-31 may influence lipid bilayer properties. First, lipid-to-lipid radial distribution profiles showed a slight peptide-dependent local clustering of anionic lipids (**Fig. S12C**). Second, based on the lateral mean square displacement of lipid phosphates, we observed marked peptide-dependent decreases in the lateral diffusivity (*D*_xy_) of all tested lipids (**Fig. S13**). Finally, to study how SS-31 binding could affect membrane packing defects, we measured the solvent accessible surface area (SASA) of acyl chains in systems with and without SS-31 (**Fig. S14**). This analysis showed peptide-dependent decreases in acyl chain solvent accessibility of all tested lipids, consistent with our observed effects of SS-31 on interfacial hydration and packing density (Fig. 4C). Hence, taken together, our MD simulations elucidate the nature of polar and nonpolar interactions that mediate the SS-31-lipid bilayer interaction, and how peptide binding may affect lipid distribution, reduce lateral lipid mobility, and decrease the accessibility of solvent (and solvated ions) to acyl chains that may otherwise be exposed to the interface by packing defects.

### SS peptide interactions with mitochondrial membranes

Having evaluated the binding of SS-31 with model membranes, the next objective was to quantitatively assess how this peptide interacts with mitochondria. We first evaluated peptide binding interactions. The spectral complexity of mitochondria precludes analysis of SS-31 binding by endogenous peptide fluorescence, so we used instead the variant [ald]SS-31, which contains the environment-sensitive fluorophore aladan (44) in place of the C-terminal Phe residue (**Fig. 1C**). As shown previously, [ald]SS-31 displays a blue-shifted emission spectrum and an increase in fluorescence intensity when bound to bilayers, making it an excellent reporter for membrane interaction. This peptide variant has also been shown to target mitochondria in a manner similar to SS-31 (11, 12). We first thoroughly evaluated the binding of [ald]SS-31 to model membranes composed of POPC with TOCL, MLCL, and POPG in different molar ratios and under different ionic conditions (**Fig. S15 A,B**). Equilibrium binding parameters for [ald]SS-31 with model membranes compared favorably with those of SS-31 (**Fig. S15C**). Hence, this variant served as a suitable model for SS-31 binding to mitochondria.

In our evaluation of SS-31 mitochondrial interactions, our first goal was to assess the CL-dependence of peptide binding. Because of the relative simplicity of the CL biosynthesis and remodeling pathway in yeast compared with higher eukaryotes (Fig. S1B), *S. cerevisiae* is an excellent model organism to directly compare effects of altered CL metabolism among otherwise isogenic systems. We therefore used mitochondria isolated from *S. cerevisiae* strains with normal CL metabolism, or with knockouts in cardiolipin synthase (Δ*crd1*) or the transacylase tafazzin (Δ*taz1*) (45). Shotgun lipidomics data confirmed the expected phospholipidome of mitochondria from these three strains (**Fig. 6A**, *left*) - namely, that Δ*crd1* mitochondria lack CL and have a buildup of PG, the immediate biosynthetic precursor of CL; and that Δ*taz1* mitochondria show an accumulation of MLCL and reduced CL. Further, the acyl chain distribution of anionic lipids among these strains shows the expected patterns of fatty acid saturation (45) (**Fig. S16A**). Emission scans of [ald]SS-31 revealed spectral changes with increasing mitochondria consistent with membrane interaction (**Fig. S16B** and **Fig. 6A**, *right*), showing two important features. First, as shown previously (8), collapse of the transmembrane potential (Δψ_m_) by the ionophore valinomycin had no effect on peptide interaction (Fig. 6A, *inset*). Second, the relative binding of [ald]SS-31 was nearly identical among the WT, Δ*crd1*, and Δ*taz1* strains. Similar peptide binding between WT and Δ*taz1* strains is expected, given that tetra-acyl TOCL and tri-acyl MLCL equally support SS-31 binding in our reductionist systems (Figs. 2 and 3). Uninhibited peptide binding by the Δ*crd1* strain is perhaps surprising given that POPG supports significantly less peptide binding than CL (Figs. 2 and 3). However, our lipidomics data indicate that PG levels are dramatically increased when CL synthesis is blocked. This increase in PG likely creates a negative surface charge density of the MIM large enough to promote peptide binding at WT levels. Again, this supports a model in which SS-31 interaction is related to surface charge, not headgroup identity.

**Fig. 6.**
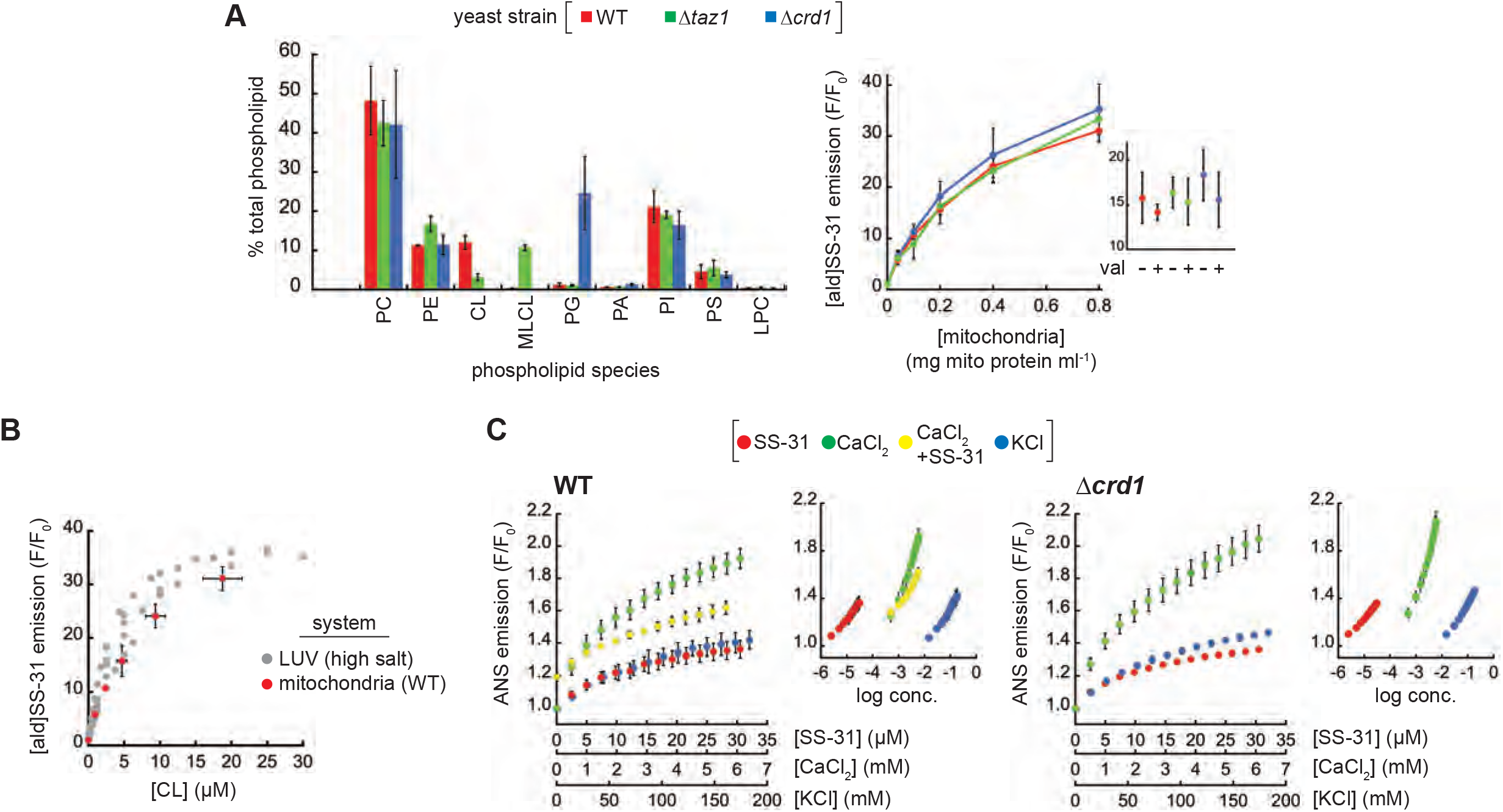
Interaction of SS-31 with isolated mitochondria. (*A*) [ald]SS-31 binding with differing phospholipid composition. *Left*, Percentages of total phospholipid of each species (PC, phosphatidylcholine; PE, phosphatidylethanolamine; CL, cardiolipin; MLCL, monolyso-CL; PG, phosphatidylglycerol; PA, phosphatidic acid; PI, phosphatidylinositol; PS, phosphatidylserine; LPC, lyso-phosphatidylcholine) in mitochondria from WT, Δ*taz1*, and Δ*crd1* strains measured by shotgun lipidomics. *Right*, binding of 1 μM [ald]SS-31 with mitochondria isolated from WT, Δ*taz1*, and Δ*crd1* yeast strains quantified as the mean fractional increase (*F*/*F*_0_) in aladan emission (n=3 ± SD) with increasing mitochondria concentration for fully energized mitochondria. *Inset*, relative binding of [ald]SS-31 to mitochondria (0.2 mg ml^−1^) in the presence or absence of 1 μM valinomycin (“val”) as indicated. (*B*) Comparison of [ald]SS-31 binding to mitochondria and model membranes. [ald]SS-31 binding is shown as a function of CL concentration for: (i) LUVs of defined TOCL composition under high salt conditions (gray, Fig. S15) and (ii) mitochondria isolated from yeast WT strain (red, Figs. S16 and 6A). Calculation of [CL] for mitochondrial samples was based on lipidomics analyses of Fig. 6A. (*C*) Effects of SS-31 and salt cations on mitochondrial surface potential. Fractional change in ANS emission is shown for titration mitoplasts from WT (*left*) and Δ*crd1* (*right*) yeast with SS-31, CaCl_2_, or KCl at the indicated concentrations (n=3 ± SD). For WT samples, CaCl_2_ titrations were also conducted following the addition of 20 μM SS-31 (“CaCl_2_+SS-31”). *Insets*, ANS emission data plotted as a function of log concentration for all three titrants.

The second goal of these binding experiments was to directly compare peptide binding between model membranes and mitochondria. We reasoned that the best comparison would come from quantifying [ald]SS-31 binding as a function of a common independent variable (CL composition), made possible using the known mitochondrial CL concentration from our lipidomics data. When evaluated in this way, the binding curves of [ald]SS-31 to LUVs and WT mitochondria are strikingly similar (**Fig. 6B**). This comparison, although indirect, is consistent with a model in which binding of SS-31 to mitochondria is governed by the same interactions in our model systems; that is, through lipid bilayer interactions.

The final goal of these experiments was to evaluate the effect of SS peptides on the surface electrostatic properties of mitochondrial membranes. Based on our observation that SS peptides attenuated the ψ_s_ of model membranes (Fig. 3C,D), we again used the ANS fluorescent reporter to measure the ψ_s_ of mitoplasts (mitochondria with osmotically ruptured outer membrane to allow probe access to the MIM). We therefore compared the titration of mitoplasts with SS-31 and mono- and divalent cations (K^+^ and Ca^2+^, respectively) to evaluate the relative effects of each species on ANS-detected ψ_s_ (**Fig. 6C**). All three cationic species caused an increase in ANS fluorescence, reflecting a decrease in the ψ_s_ of mitochondrial membranes from WT (*left*) and Δ*crd1* (*right*) yeast. Consistent with the exponential dependence of formal charge on accumulation of ionized species in the electric field of membranes (Eq. S11), the effective concentration ranges of SS-31, Ca^2+^, and K^+^ progressively differed by about two orders of magnitude (Fig. 6C, *insets*). The first notable feature of this analysis is the relative effect of each species on surface potential. Compared with K^+^, Ca^2+^ ions caused a much stronger impact on the relative ψ_s_, as expected. Interestingly, however, within its effective concentration range, SS-31 had a reduced effect on the relative ψ_s_ that was closer to that of the monovalent cation – an observation consistent with the fact that SS-31 does not completely reverse negative surface potential of model membranes even upon binding saturation (Fig. 3C). The second notable feature of this analysis is the effect of SS-31 on Ca^2+^-mediated decreases in the relative ψ_s_. SS-31 addition to WT mitoplasts prior to CaCl_2_ titration strongly blunted the effects of Ca^2+^ on surface potential (Fig. 6C, *left, compare green and yellow symbols*), supporting a binding model in which SS-31 preferentially binds to mitochondrial membranes over Ca^2+^ ions. Taken together, we conclude from the results of Fig. 6 that the binding of SS peptides to mitochondrial membranes is strongly dependent on surface charge of the lipid bilayer, that SS peptide binding causes a “controlled down-tuning” of the electric field originating from the membrane surface, and that SS peptides at the membrane interface can mitigate the effects of polyvalent cations on membrane surface electrostatics.

### The effects of SS peptides on cation accumulation at the membrane interface

The final analyses of this study were focused on testing the functional implications of our observed effects of SS peptides on membrane surface electrostatics. We began with a theoretical consideration of how SS peptides may affect the decay of the electric field and ion distribution at the membrane surface based on GCS formalism (Eq. S10-S12) (**Fig. 7A**). By this analysis, SS peptides (modeled as trivalent ions at a bulk concentration in the nM range) accumulate at the membrane surface with the effect of: (i) reducing the magnitude of the surface charge, and (ii) decreasing accumulation of other cationic species at the interface. This effect is particularly relevant for Ca^2+^, because mitochondria serve as high-capacity calcium stores in mediating cellular ion homeostasis and Ca^2+^ overload can cause severe damage to anionic lipid bilayers, particularly within mitochondria (see Discussion). The analysis of Fig. 7A shows that SS peptides cause a strong reduction in surface accumulation of divalent cations, by over an order of magnitude. Hence, this model, albeit a simplified one, makes testable predictions about how SS peptides may mitigate divalent cation stress by reducing Ca^2+^ accumulation at the membrane interface. Our model was tested as follows.

**Fig. 7.**
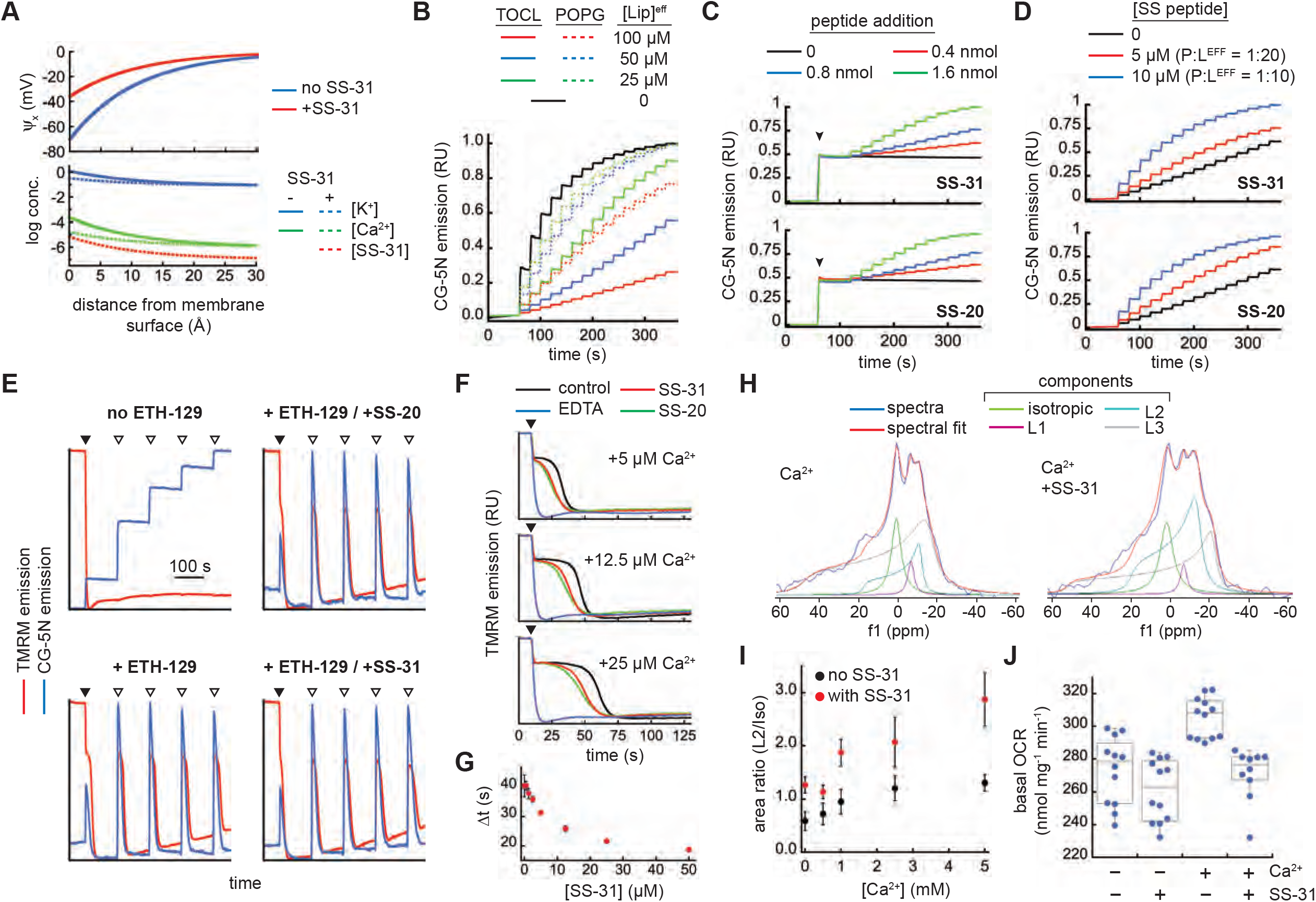
SS peptide effects on Ca^2+^-membrane interactions and on mitochondrial cation stress. (*A*) Profiles of surface potential decay (*upper panel*) and ion distribution (*lower panel*) based on GCS theory in the presence or absence of SS-31 assuming a lipid bilayer of 20% CL and bulk concentrations as follows: monovalent 1:1 electrolyte (e.g., Na^+^Cl^−^), 80 mM; divalent 2:1 electrolyte (e.g., Ca^2+^Cl^−^_2_), 1 μM; and SS-31 (where present), 100 nM. (*B-D*) Effect of SS peptides on Ca^2+^ accumulation at model membrane interfaces. Time course measurements of CG-5N emission were performed as follows. (*B*) The effect of anionic bilayers on the CG-5N response to added Ca^2+^ was evaluated by preparing solutions in the absence or presence of LUVs (20 mol% TOCL or 20 mol% POPG in a host POPC background) at the concentrations indicated. Starting at t=60 s, CaCl_2_ was titrated in increments of 1 nmol at 20 s intervals. (*C,D*) The effect of SS peptides on interfacial Ca^2+^ binding was evaluated by preparing solutions with LUVs (20 mol% TOCL, [Lip]^eff^ = 100 μM) and (*C*) subjecting samples to 5 nmol CaCl_2_ addition (arrowhead, t=60 s) followed by addition of SS-31 or SS-20 at the indicated amounts at 20 s intervals, or (D) pre-binding SS peptide at the indicated concentrations followed by titration of CaCl_2_ in increments of 1 nmol at 20 s intervals starting at t=60s. (*E-G*) Effect of SS peptides on mitochondrial Ca^2+^ uptake. Time courses of mitochondria (50 μg ml^−1^) were performed in buffer containing TMRM (Δψ_m_ measurements) or CG-5N (external [Ca^2+^] measurements). To facilitate comparison, both TMRM and CG-5N emission data were normalized relative to the full range of the probe response over the course of each measurement. (*E*) TMRM and CG-5N traces of Δψ_m_ generation following addition of 1 mM NADH (black arrowheads) and of calcium transients following addition of 25 μM CaCl_2_ (open arrowheads) in samples pre-treated with or without 5 μM ETH-129, 25 μM SS-20, and/or 25 μM SS-31 as indicated. (*F*) TMRM traces of Δψ_m_ generation following addition of 1 mM NADH (black arrowheads) in samples containing the indicated [CaCl_2_] and supplemented with vehicle only (control), 25 μM EDTA, SS-20, or SS-31 as indicated. (*G*) Dose response of SS-31 on Δψ_m_ generation with 25 μM CaCl_2_ (defined in Fig. S18B, n=3 ± SD). (*H-J*) Effect of SS-31 on external Ca^2+^ stress. (*H,l*) ^31^P ssNMR analysis. Mitochondria (70 mg ml^−1^) were given acute treatment of SS-31 (0.6 mM) or vehicle only followed by calcium treatment (up to 5 mM CaCl_2_). (*H*) Static wide-line ^31^P NMR spectra of mitochondria subject to calcium stress (5 mM CaCl_2_) following treatment without (*left*) or with (*right*) SS-31. Spectral deconvolution yielded the indicated components (see Fig. S19), which were used to quantify the area ratios of L2:iso (n=3 ± SD) in panel I. (*J*) Respirometry analysis. Mitochondria (1 mg ml^−1^) were treated in the absence or presence of SS-31 (54.4 μM) or CaCl_2_ (1 mM) as indicated and measured for state 2 respiration (n=10-12).

First, we measured the effects of SS peptides on the accumulation of Ca^2+^ in the interfacial region of model membranes (**Fig. 7B-D**). To this end, we used a fluorescence-based approach with Calcium Green-5N (CG-5N), a membrane-impermeant probe that exhibits an increase in emission intensity upon binding Ca^2+^ (characterized for our system in **SI Results H** and **Fig. S17A,B**). We reasoned that the binding of Ca^2+^ to anionic lipid bilayers in the headgroup region would reduce the bulk Ca^2+^ detected by the CG-5N probe, providing a direct readout of Ca^2+^ accumulation at the membrane interface. As a proof of principle, we titrated Ca^2+^ into solutions of CG-5N containing LUVs with monoanionic (POPG) or dianionic (TOCL) lipids at different concentrations (**Fig. 7B**). In the absence of LUVs, there was a saturating increase in CG-5N emission as the probe became maximally bound with Ca^2+^. However, the presence of POPG- and TOCL-containing LUVs caused a reduction in the CG-5N response to Ca^2+^ that was directly related to the amount of bilayer surface charge in the system. Hence, this assay provided a direct measure of Ca^2+^ sequestration within the headgroup region of anionic bilayers. To test the effects of SS peptides on Ca^2+^ binding to membrane surfaces, we first titrated SS-20 and SS-31 into solutions containing LUVs with 20% TOCL pre-bound with Ca^2+^ (**Fig. 7C**). These results showed that SS-20 and SS-31 released Ca^2+^ from the membrane interface in a dose-dependent manner that was identical between the two peptides. Importantly, the CG-5N response began to saturate near P:L^eff^ of ~1:10 (Fig. 7C, green traces), corresponding to the peptide binding saturation point, therefore supporting that Ca^2+^ release into the bulk corresponded to peptide bilayer binding. As a complementary approach, we performed Ca^2+^ titration on solutions of LUVs that had been pre-incubated without peptide or with SS peptides at molar P:L^eff^ of 1:20 and 1:10 (**Fig. 7D**). These results confirmed that the presence of SS-20 and SS-31 both reduced the equilibrium binding of Ca^2+^ to anionic bilayer surfaces in a dose-dependent manner. Taken together, these results support a model in which SS peptide binding to bilayer surfaces reduces the accumulation of Ca^2+^ ions within the interfacial region of model membranes.

Second, we measured the effects of SS peptides on mitochondrial Ca^2+^ flux (**Fig. 7E-G**). External Ca^2+^ is taken up into the mitochondrial matrix electrophoretically by the Δψ_m_ (matrix negative). We reasoned that if SS peptides altered the accumulation of cations at the bilayer surface (Fig. 7B-D), then they may have a measureable effect on mitochondrial Ca^2+^ uptake. We therefore performed time course measurements of isolated mitochondria supplemented with respiratory substrate NADH followed by additions of calcium. In these experiments, we monitored the relative Δψ_m_ using the potentiometric probe TMRM (the emission of which is inversely related to membrane potential, **Fig. S17C**) and in parallel experiments we monitored the external [Ca^2+^] using CG-5N (**Fig. 7E**). Note that because S. *cerevisiae* mitochondria do not contain a Ca^2+^ uniporter, Ca^2+^ uptake does not occur spontaneously but can occur in the presence of a calcium ionophore, for which we used ETH-129. These time courses show two key processes: (i) following NADH addition, there is a TMRM-detected establishment of the Δψ_m_, and (ii) after each calcium addition, there is a CG-5N-detected spike in external [Ca^2+^] followed by a decrease in CG-5N emission as Ca^2+^ is taken into the matrix, which is accompanied by a transient TMRM-detected partial depolarization and re-establishment of the Δψ_m_ (see **Fig. S18A** for expanded views of each of these transient processes). The absence of dynamic changes in TMRM and CG-5N emission in samples lacking ETH-129 (Fig. 7E, “no ETH-129”) confirmed that these spectral changes did indeed reflect Ca^2+^ uptake. These experiments revealed two notable effects of SS peptides. The first was related to the establishment of the Δψ_m_ in samples containing ETH-129. Upon reaching a threshold potential, there was a temporal lag in potential generation, after which the mitochondria resumed establishing the maximal Δψ_m_. This is attributable to the energetic demands coupled to the uptake of calcium that exists in the external buffer of these mitochondrial samples (see also Fig. S17D). Interestingly, both SS-20 and SS-31 reduced the extent of this temporal delay in mitochondrial energization observed in the presence of increasing [Ca^2+^] (**Fig. 7F** and Fig. S18A, “Δψ_m_ establishment”). Quantitative analysis of Ca^2+^-dependent Δψ_m_ generation (described in **Fig. S18B**) revealed that SS-31 reduced this temporal delay in a dose-dependent and saturable manner (**Fig. 7G**). The second related effect was associated with subsequent calcium spikes. Following calcium additions, mitochondria transiently depolarized to near the same Δψ_m_ associated with the temporal lag. The kinetics of repolarization during the first calcium additions were faster in the presence of SS peptides; however, with increasing calcium load, the peptides no longer enhanced repolarization (see Fig. S18A, “Ca^2+^ transient 1-4”). By contrast, external calcium flux, judging by the extent of the CG-5N signal and by the kinetics of Ca^2+^ uptake, was not affected at all by SS peptides. Hence, under these conditions, SS peptides do not measurably alter the calcium storage capacity of mitochondria, but do improve the ability to respond energetically to calcium load. These results may provide a mechanistic underpinning to previous observations that acute treatment of isolated mitochondria with SS peptides protected against Ca^2+^-induced onset of the mitochondrial permeability transition (MPT) (8).

Finally, we measured the effects of SS peptides on mitochondrial calcium stress that may occur independently of Ca^2+^ uptake into the matrix (**Fig. 7H-J**). To this end, we used a ^31^P ssNMR-based strategy to monitor mitochondrial membrane structural integrity, based on previous studies using NMR analysis of membranes from prokaryotic cells (46) or isolated mitochondria (47–49). We first characterized ^31^P ssNMR spectra of our isolated mitochondria in comparison to model membranes (**SI Results H** and **Fig. S18**), showing that mitochondria spectra contained a complex superposition of lineshapes that could be deconvoluted for the direct analysis of phospholipid bilayers. Based on previous work using NMR analysis to measure the effects of calcium (up to 10 mM) on mitochondrial membranes (49), we then tested the ability of SS-31 to mitigate damage to mitochondrial membranes by calcium stress. We found that acute treatment of mitochondria with SS-31 (8.5 nmol peptide per mg mitochondrial protein) reduced NMR-detected membrane degradation caused by high calcium load (up to 5 mM CaCl_2_) (**Fig. 7H**). By spectral deconvolution, we observed that SS-31 treatment resulted in a significant preservation of the L2 peak, which corresponds to lamellar phospholipid bilayers (**Fig. 7I, Fig. S19**). To corroborate these findings, we measured mitochondrial oxygen consumption rates under similar conditions of SS peptide treatment and calcium stress. Under these conditions, we observed a significant increase in basal respiration caused by calcium addition that was prevented by SS-31 pretreatment (**Fig. 7J**). Hence, acute SS-31 treatment rendered protective effects under these harsh conditions of calcium stress manifested as the preservation of membrane integrity and OXPHOS activity.

## Discussion

It is well established that SS-31 specifically targets mitochondria and ameliorates the decrease in mitochondrial function associated with aging, cellular stress, and heritable diseases. Further, SS-31 is known to have affinity for aqueous dispersions of anionic lipids (12), particularly membranes containing CL (10). However, the nature of the interaction between SS-31 and mitochondrial membranes has remained elusive. Toward the goal of understanding the molecular MOA of SS-31, the present study addressed how this peptide interacts with lipid bilayers and the effect that it has on membrane properties. Our results support a mechanism in which the modulation of mitochondrial membrane surface electrostatic properties plays a critical role.

The first key finding of this study was on the lipid determinants of SS-31 membrane interactions, based on empirical binding isotherms with model membranes (Figs. 2, S15) and MD simulations (Figs. 5, S11). Consistent with previous results (10, 12), we found that SS-31 had negligible interaction with membranes composed solely of zwitterionic lipids, and required anionic lipids for saturable binding. Independent lines of evidence in this study support that the primary determinant of SS-31 binding is the number of negative charges within the lipid headgroup region. First, the binding of SS-31 to membranes containing tetra-acyl CL (TOCL) or its lysolipid variant (MLCL) was virtually indistinguishable, as demonstrated in model membranes (Fig. 2) and isolated mitochondria (Fig. 6A, WT *vs*. Δ*taz1*). Notably, this supports the ability of SS-31 to act as a therapeutic for the treatment of Barth syndrome, which is characterized by an increase in the MLCL/CL ratio of the MIM (24). Second, membranes containing monoanionic PG supported roughly half of the SS-31 binding capacity observed in membranes with equal concentrations of CL (Fig. 2). In this regard, mitochondria lacking CL biosynthesis sustained near-WT levels of peptide binding because PG was increased to roughly twice the normal CL concentration (Fig. 6A, WT vs. Δ*crd1*). Together, these results support a model in which SS peptide binding to lipid bilayers depends on the anionic surface charge density, not the identity of a particular component lipid. This model is supported by our atomistic MD work, showing that bilayer-docked SS-31 resides stably in the interfacial region with similar residue-specific penetration depths and lipid interactions among TOCL, MLCL, and POPG-containing bilayers (Figs. 5, S11, S12). Yet based on ITC measurements, the noncovalent interactions that mediate SS-31 binding appear to be dominated by entropy rather than by the energy of polar contacts (Fig. 2D,E), underscoring the importance of nonpolar interactions between aromatic side chains of the peptide and the acyl chain region in determining binding stability. Therefore, a key role of the bilayer surface charge is likely the establishment of a strong electric field (see Fig. S2) that increases the effective concentration of peptides in the interfacial region, where they are then able to establish binding interactions, which are dominated by hydrophobic contacts (**Fig. 8A**). We note that although CL is not strictly required for SS-31 binding, it is likely essential for peptide action within mitochondrial membranes for two main reasons. First, under conditions of normal lipid metabolism, CL is the lipid that contributes the most to the strong negative surface charge of the MIM. As shown by our lipidomics data (Fig. 6A), the concentration of dianionic CL is greater than that of other monoanionic phospholipids (PS, PA, PG); other highly charged lipids including phosphoinositides (the phosphorylated variants of PI) are generally confined to cytoplasmic leaflets of the endomembrane system (50) and not abundant in mitochondria. Second, the inverted conical molecular geometry of CL, promoted by small headgroup volume and highly unsaturated fatty acids (cf. Fig. S16A), creates packing defects in CL-rich bilayers, which may be critical for providing peptide side chain access to acyl chains. As a corollary, specific physicochemical features of CL may be essential for downstream (post-binding) effects of SS peptides on membrane properties (see below). Hence, considering that polypeptide binding to lipid bilayers is determined by a combination of surface charge density and packing defects (51), we propose that SS peptides target the specialized environment of the MIM through a combination of a strong surface electrostatics and lipid packing defects imparted by features such as high [CL], highly unsaturated acyl chains, and low sterol content.

**Fig. 8.**
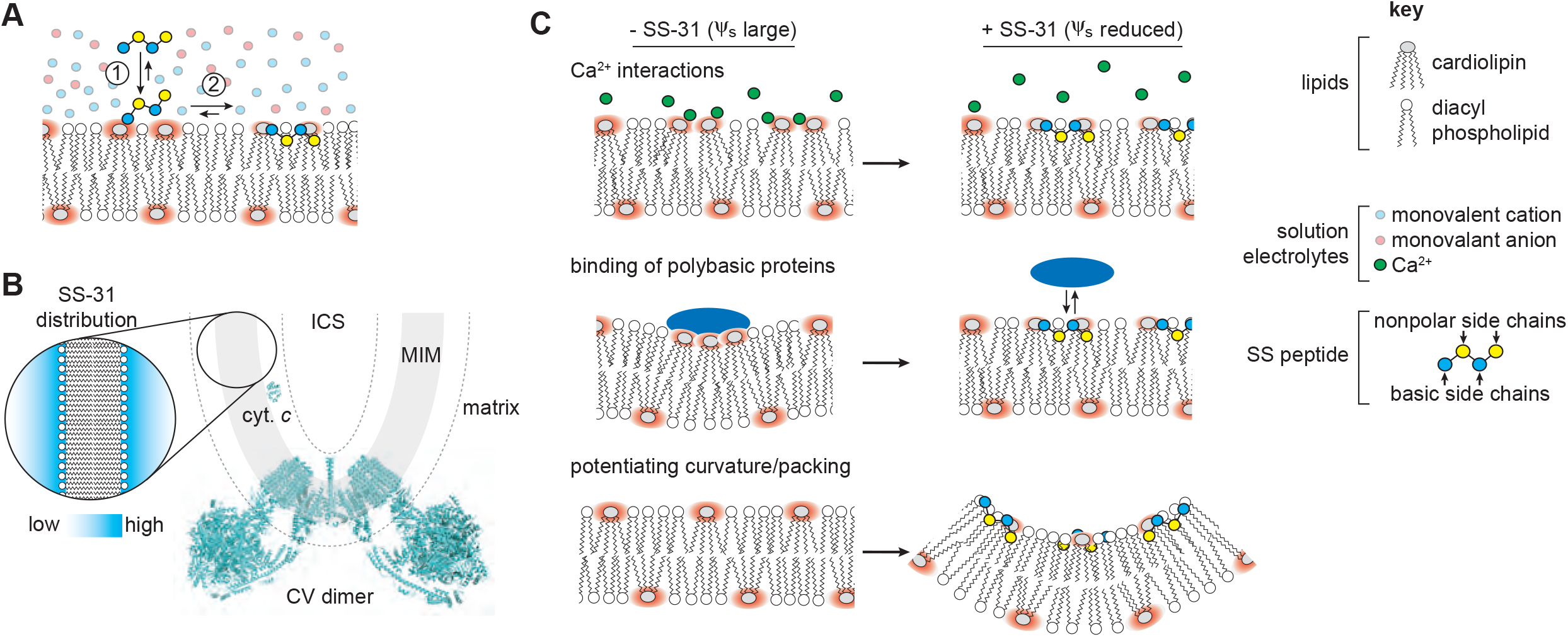
SS peptide distribution and modulation of membrane surface electrostatics. (*A*) Equilibrium binding of SS-31: (1) Polybasic peptide is drawn to bilayer surface under the force of the electric field originating from negative surface charge density. (2) Peptide interaction stabilized by polar contacts (basic side chains and headgroup phosphates) and hydrophobic interactions (burial of nonpolar side chains in acyl chain region). (*B*) Depiction of SS peptide equilibrium distribution at mitochondrial membranes showing mitochondrial cristae region, including the mitochondrial inner membrane (MIM), which delineates aqueous matrix and intracristal space (ICS). Topological positioning of the amphitropic protein cytochrome *c* (cyt. *c*, PDB 1ycc) and F_∘_F_1_ ATP synthase (CV dimer, PDB 6b8h) are shown for comparison. Dashed line represents region of ~30Å (2-3 Debye lengths) from the membrane surface. *Inset*, representation of SS peptide distribution within the diffuse double layer. (*C*) Working models for how regulation of mitochondrial surface charge may underpin the efficacy of SS peptides. (1 and 2) divalent cations and basic protein binding cause lipid demixing and sequestration of CL with implications for membrane integrity, lipid-protein interactions, and lipid peroxidation. The binding of SS peptides to CL-containing bilayers shifts the equilibrium binding of cations and basic proteins away from the membrane surface. (3) SS peptide binding alters physical properties of CL-containing bilayers. The ionized CL headgroup creates a strong local electric field (red) that alters lateral lipid interactions and creates a strong ψ_s_. Interfacial binding of SS-31 reduces ψ_s_, thereby enhancing lipid packing and promoting local curvature.

The second key finding of this study was the equilibrium membrane binding parameters of SS-31 and how they elucidate the molecular MOA. First, the lipid:peptide stoichiometry (*n*) serves as an index for peptide binding density (*n* scales inversely with the number of peptides bound per unit membrane area). We show that under conditions of maximal peptide binding density, model membranes composed of 20 mol% CL and 20 mol% PG bind SS-31 with n ≈ 6-7 (area occupancy ~500-600 Å^2^) and n ≈ 15 (area occupancy ~ 1000 Å^2^), respectively. To put these values into context, consider a typical cell with a total mitochondrial volume of 2×10^3^ μm^3^ and a total MIM surface area of 2×10^5^ μm^2^ (52). Assuming, conservatively, that 10% of the MIM surface is available for peptide binding, this translates into a mitochondrial SS-31 concentration of 400 μM at maximal binding density. This estimate, although approximate, is consistent with the observed 1000 to 5000-fold mitochondrial accumulation in cells incubated with low nM amounts of peptide (8). Second, the binding affinity (*K*_D_) of SS-31 for membranes was found to be in the low μM range. Compared with typical drug compounds that bind molecular targets with much higher affinity (e.g., *K*_D_ in the nM and lower range), the binding affinity of SS peptides to membranes is very weak. This low affinity range is to be expected, considering that the primary mode of peptide interaction is with the thermally disordered lipid bilayer rather than with a molecular binding pocket with a defined covalent structure. Furthermore, the low μM affinity of SS-31 for membranes is consistent the pharmacokinetic profile of SS peptides, showing rapid clearance of the peptide after administration (13) as well as peptide release from cultured cells upon media exchange (7). Notably, the correspondence between SS-31 binding parameters with model membranes and with isolated mitochondria observed in this study supports a model in which the primary mode of SS peptide interaction is with lipid bilayers. Notably, it remains possible (and perhaps likely) that a fraction of SS peptides may interact with proteins (e.g., within acidic pockets) and/or at protein-lipid interfaces, particularly given the protein-rich nature of mitochondrial membranes.

The third key finding of this study addressed the effects of SS peptides on membrane physical properties. Using ^31^P NMR and SAXS, we found that SS-31 did not disturb the lamellar bilayers of model LUVs, even at high peptide binding densities (Fig. 4A,B). This is consistent with previous work (8) and the present study (Fig. S17E), showing that SS peptides, even at high concentrations, do not dissipate ion gradients (ΔΨ_m_) of respiring mitochondria. As noted above, given their amphipathic character, SS peptides resemble AMPs. This shared amphipathicity likely drives the interactions with bacterial membranes (AMPs) and membranes of bacterial origin (SS peptides). We propose that the short (four-residue) length of SS peptides is the predominant feature that keeps them from disrupting membrane structure, unlike AMPs, which are significantly longer (40). SS-31 did, however, affect properties of the interfacial membrane region in a dose-dependent manner. First, based on membrane-bound solvatochromic probes, SS-31 caused a modest but repeatable increase in lipid packing, particularly for CL-containing bilayers (Fig. 4C), a result consistent with our analysis of peptide-dependent reduction in acyl chain solvent accessible surface area (Fig. S14). Due to its conical molecular geometry, CL can cause localized exposure of acyl chains to the aqueous interface (27). Intercalation of SS-31 aromatic chains into these pockets could fill such interfacial “voids” (packing defects), which could in turn have implications for bilayer properties such as mechanostability and elasticity. Second, based on computational approaches, we found that SS-31 caused a reduction in the lateral diffusivity of lipids (Fig. S14), consistent with a model in which peptide binds tightly enough to anionic lipids that they co-diffuse for significant (at least μsec) time scales. Third, using electrokinetic and fluorescence-based approaches, we found that SS-31 decreases the surface charge (and hence the ψ_s_) of both model (Fig. 3C,D) and mitochondrial (Fig. 6C) membranes. All of these observed effects are likely interrelated. For example, by partially neutralizing the charge of anionic lipids, the binding of SS-31 will reduce the hydration and effective cross-sectional area of headgroups, as well as decrease inter-lipid electrostatic repulsion, all of which could increase lipid packing. We therefore propose a model for the molecular MOA of SS-31 in which the primary effect of bilayer interaction is the down-regulation of the ψ_s_ of mitochondrial membranes. In support of this model, we found that SS peptides caused the partial neutralization of the strong negative surface charge density of mitochondrial membranes, but appeared to do so in a way that does not cause positive charge overcompensation (Figs. 3C and 6C). A critical corollary of this model is that this effect on surface electrostatics should be shared by all SS peptides due to their common positive charge density, and not depend on specific side chain features such as the ROS scavenging potential of the 2’,6’-Dmt moiety of SS-31. Indeed, we found that SS-20 and SS-31 are indistinguishable in terms of their effects on tuning membrane surface potential.

In this model, it is important to consider how peptides may distribute within the membrane interface. SS peptide occupancy at or near the surface of anionic membranes is driven by the high (+3) positive charge density that dictates peptide equilibrium distribution in the electrostatic field of the bilayer and promotes ionic interactions with negatively charged membrane-bound groups. Hence, SS peptides bind the membrane surface with micromolar affinity that is entropically and enthalpically favored (Fig. 2); however, given that the binding is relatively weak and reversible depending on local ionic strength (Fig. 3), a fraction of these small polybasic amphiphiles are likely to exist untethered to the membrane surface, but stably reside within the electric field of the double diffuse layer of the anionic mitochondrial membranes (**Fig. 8B**). This distribution is likely to vary laterally along the MIM in different mitochondrial subcompartments that have different surface charge characteristics. Noting that ion distribution near a charged membrane surface is an exponential function of formal charge (Eq. S11), the polybasic nature of SS peptides is critical in allowing them to bind anionic bilayers in competition with physiological concentrations of mono- and divalent cations (trivalent cations are rare in biological systems). Hence, in accordance with GCS theory (Fig. 7A), the accumulation of SS peptides at the interface will have the effect of shifting the balance of other ionic species (e.g., solvated metal ions and amphitropic basic proteins) away form the membrane surface. Finally, given that CL may be enriched on the inner leaflet of the MIM (53), SS peptides may preferentially accumulate on the matrix-facing side of this membrane, provided that a pathway exists for them to traverse the MIM. Ongoing work in our group is aimed toward understanding the dynamic distribution of SS peptides among mitochondrial subcompartments.

How might the down-regulation of the ψ_s_ of anionic mitochondrial membranes underpin the broad therapeutic efficacy of SS peptides? We propose several nonexclusive possibilities (**Fig. 8C**). First, reducing the magnitude of the membrane surface charge will alter cation distribution at the interfacial region. This is particularly relevant for Ca^2+^ because mitochondria must accumulate large concentrations of this divalent cation in their role as cellular calcium stores. Ca^2+^ binding to the lipid headgroup region severely alters properties of anionic bilayers (48). More relevant to mitochondrial membranes, Ca^2+^ interacts strongly with CL, altering bilayer properties (e.g., demixing, phase behavior, and headgroup conformation) (19); Ca^2+^ can disrupt protein-lipid interactions, causing effects such as respiratory complex disintegration (54); and mitochondrial Ca^2+^ dyshomeostasis underpins a wide variety of mitochondrial diseases (55). In this study, we have directly shown that the binding of SS peptides reduces the effects of Ca^2+^ on mitochondrial surface electrostatics (Fig. 6C) and reduces the equilibrium binding of Ca^2+^ to the interface of anionic synthetic membranes, which act as a “sink” for divalent cations (Fig. 7B-D). We also demonstrated that SS peptides appear to reduce the energetic burden associated with calcium uptake (Fig. 7E-F) and preserve mitochondrial function and membrane integrity with calcium stress (Fig. 7G-I). Although we do not directly show the latter two physiological effects originate specifically from the reduction of Ca^2+^ accumulation at the bilayer surface, evidence that the site of action of SS peptides is at the membrane interface supports this as a likely mechanism. This proposed effect of SS peptides on mitigating bilayer Ca^2+^ accumulation is consistent with their demonstrated efficacy toward processes coupled to pathogenic mitochondrial Ca^2+^ overload, including ischemia-reperfusion, induction of the MPT, and apoptosis/necrosis (5). A second related mechanism is that reducing the magnitude of the ψ_s_ could reduce the mitotoxic interaction of basic proteins with CL-rich mitochondrial membranes. These include cytochrome *c* (cyt. c), whose binding to the MIM leads to lipid peroxidation and initiates the intrinsic apoptotic pathway (56), and neurotoxic amyloid peptides, known to aggregate upon and damage mitochondrial membranes (57, 58). To this point it is noteworthy that SS-31 has been shown to alter cyt. *c* interactions with CL-containing bilayers (10, 11) and to reduce mitochondrial damage in models of Alzheimer’s and Parkinson’s disease (59, 60). Third, by downregulating the surface charge density of mitochondrial membranes, SS peptides may act in part by altering the physical properties of CL-containing bilayers. Full ionization of CL phosphate groups creates strong electrostatic repulsion in the lipid headgroup region and increases the effective headgroup area, imparting a more cylindrical lipid geometry. Reduction of CL headgroup charge promotes more conical molecular geometry, thereby altering lipid packing interactions and making the establishment of membrane curvature more energetically accessible (20, 61–63). SS peptides may therefore help stabilize local regions of high curvature in the morphologically complex MIM. A final potential mechanism could relate to recent findings that dysfunctional mitochondrial protein complex assembly is coupled to the release of CL into the bulk lipid phase (64); under such pathological conditions, SS peptides could help regulate the resulting increase in membrane surface charge.

With these proposed mechanisms in mind, there are two ways to consider the broad efficacy of SS peptides in treating mitochondrial dysfunction. First, the controlled down-regulation of the mitochondrial ψ_s_ could have direct and specific impacts on different molecular pathomechanisms: for instance, by altering the molecular geometry of MLCL to impart more conical-like character, SS peptides could help stabilize cristae curvature in patients with Barth syndrome. But from a more general standpoint, SS peptides could also serve to reduce the constant stresses that mitochondria face under normal physiological conditions, including spikes in calcium levels and mitotoxic interactions between CL and polybasic proteins. Such stresses could present a manageable burden on mitochondria of healthy individuals, but an excessively taxing one on mitochondria whose functional integrity is compromised, as occurs with ageing and heritable mitochondrial disease. Hence, the remarkably broad effects of SS peptides in treating mitochondrial dysfunction could also be based partly on their ability to mitigate general molecular stressors at the membrane.

In summary, the results of this study support a model in which SS-31 binding to mitochondrial membranes is largely driven by, and subsequently downregulates, the ψ_s_ as a core component of its molecular MOA. Considering SS peptides as mitochondria-targeted hydrophobic polyvalent cations, this model explains how these compounds act upon the general property of bilayer surface electrostatics to serve as broad-based therapeutic compounds for a wide array of mitochondrial disorders.

## Acknowledgements

The authors would like to thank Dr. Steven Claypool (Johns Hopkins University School of Medicine) for the original synthesis of yeast mutants used in this study, Drs. Mu-Ping Nieh and Kuo-Chih Shih (University of Connecticut) for technical assistance with SAXS measurements, Dr. Georg Pabst (University of Graz) for providing the GAP program for analysis of SAXS data, and the UT Health-San Antonio Mass Spectrometry Core Facility. This work was supported by the National Institutes of Health (R01-GM113092 to N.N.A., R35-GM1197623 to E.R.M., and RF1-AG061872 to X.H.), by the National Science Foundation (GRFP award 1247393 to K.B.) and by a charitable contribution from the Social Profit Network (to N.N.A.)

