## Supplementary Information for "Molecular Mechanism of Action of Mitochondrial Therapeutic SS-31 (Elamipretide): Membrane Interactions and Effects on Surface Electrostatics"

### Table of Contents

**SI Methods and Materials** — **Page**
A. Reagents and sample preparation 1
B. Experimental and analytical methods 2
C. Molecular dynamics methods 6
D. Quantitative analysis of peptide binding isotherms and surface electrostatics 9

**SI Results**
A. Spectral characterization of SS-31 in solution 13
B. Fluorescence-based analysis of SS-31 membrane interactions 14
C. Quantitative analysis of SS-31 membrane binding isotherms 16
D. SS-31 titration time course analysis 17
E. Model membrane surface electrostatics and SS-31 binding 18
F. Assays for peptide-induced structural polymorphism of model membranes 19
G. Molecular dynamics analyses 22
H. Assays for measuring $Ca^{2+}$ interactions with model membranes and mitochondria 25

**SI References** 29

**SI Figures**
Figure S1. Cardiolipin structure and biogenesis 34
Figure S2. The binding of amphipathic peptides to anionic membranes 35
Figure S3. Fluorescence characterization of SS-31 in aqueous buffer 36
Figure S4. Fluorescence characterization of SS-31 membrane interactions 37
Figure S5. Quantitative analysis of peptide membrane binding 38
Figure S6. Kinetic measurements of SS-31 addition to LUVs 39
Figure S7. Analysis of time course binding saturation measurements 40
Figure S8. Microcalorimetry analysis of SS-31 membrane binding 41
Figure S9. Relationship between SS peptide binding, ionic strength, and surface potential 42
Figure S10. Characterization of the effects of SS-31 on model membranes 43
Figure S11. Molecular dynamics simulations of SS-31 side chain insertion depths 44
Figure S12. Lipid radial distribution profiles from MD simulations 45
Figure S13. MSD of lipids with and without SS-31 from MD simulations 46
Figure S14. Acyl chain solvent accessible surface area from MD simulations 47
Figure S15. [ald]SS-31 binding isotherms with model membranes 48
Figure S16. Binding of [ald]SS-31 to isolated mitochondria 49
Figure S17. Characterization of fluorescence-based assays of calcium dynamics 50
Figure S18. Kinetic responses of $\Delta\Psi_m$ and external [$Ca^{2+}$] 51
Figure S19. NMR and respirometry measurements with calcium stress 52

### Supplementary Methods and Materials

#### A. Reagents and sample preparation

*1. Reagents*

Peptides SS-31 and SS-20 were prepared by solid phase synthesis by Phoenix Pharmaceuticals (Burlingame, CA) and [ald]SS-31 was synthesized as described (1). All peptides were prepared as aqueous stocks. Synthetic phospholipids were purchased as chloroform stocks from Avanti Polar Lipids (Alabaster, AL), including 1-palmitoyl-2-oleoyl-*sn*-glycero-3-phosphocholine (POPC), 1-palmitoyl-2-oleoyl-*sn*-glycero-3-phosphoglycerol (POPG), and variants of cardiolipin with 18:1 acyl chains, including 1',3'-bis[1,2-dioleoyl-*sn*-glycero-3-phospho-]-*sn*-glycerol (TOCL) and trioleoyl monolysocardiolipin (MLCL). All lipid stocks were stored at -20°C in clear vials with Teflon-lined cap closures until use. Fluorescent probes TMRM (tetramethylrhodamine methyl ester), Laurdan (6-dodecanoyl-2-dimethylaminonaphthalene), Prodan (6-propinoyl-2-dimethylaminonaphthalene), DPH (1,6-diphenyl-1,3,5-hexatriene), ANS (1-anilinonaphthalene-8-sulfonic acid), and Calcium Green-5N (CG-5N, glycine, N-[2-[2-[2-[bis(carboxymethyl)amino]-5-[[(2',7'-dichloro-3',6'-dihydroxy-3-oxospiro[isobenzofuran-1(3H),9'-[9H]xanthen]-5-yl) carbonyl] amino] phenoxy]ethoxy]-4-nitrophenyl]-N-(carboxymethyl)-hexapotassium salt) were purchased from Molecular Probes (ThermoFisher Scientific). Solutions were prepared with deionized ultrapure water and all buffers, salts and other reagents were obtained from Sigma-Aldrich (St. Louis, MO, USA).

*2. Liposome preparation*

Liposomes of a specific lipid composition were prepared by mixing chloroform stocks at the appropriate molar ratios in a glass vials and drying under $N_2$ flow, followed by incubation in a vacuum desiccator for a minimum of two hours to remove all organic solvent. Lipid films were hydrated in aqueous solutions to produce dispersions of multilamellar vesicles (MLVs). To prepare large unilamellar vesicles (LUVs), lipid suspensions were extruded through polycarbonate membranes with 0.1 μM pores (Whatman, Maidstone, UK) (2). MLVs and LUVs were prepared in aqueous solutions of buffer (20 mM HEPES-KOH, pH 7.5) in the presence or absence of different concentrations of salt (KCl) depending on the application. Hydration and extrusion steps were all performed at a temperature above the main (gel to liquid phase) temperature of the highest-melting lipid. The phospholipid concentration of each preparation was determined by colorimetric ammonium ferrothiocyanate assay (3) against standards based on the mole fraction of lipids of a given preparation. Liposome suspensions were routinely monitored for size and monodispersity by dynamic light scattering.

*3. Yeast strains and isolation of mitochondria*

The yeast strains used in this study are isogenic to GA74-1A, including genetic knockouts Δ*crd1* and Δ*taz1*, which were described previously (4). Yeast cells were grown at 30°C in lactate medium containing 0.1% glucose. For the preparation of active mitochondria, yeast cells were cultivated to mid-log phase and mitochondria were isolated by dounce homogenization of spheroplasts and differential centrifugation as described (5). Isolated mitochondria were resuspended in buffer (600 mM sorbitol, 20 mM HEPES-KOH, 1 mM EDTA, pH 7.5) and aliquots (1-2 mg mitochondrial protein each) were snap frozen in liquid nitrogen and stored at -80ºC. Mitoplasts were prepared by osmotically rupturing the OM as described (5) by incubating intact mitochondria in buffer (60 mM sorbitol, 20 mM HEPES, pH 7.5) on ice for 30 min.

**B. Experimental and analytical methods**

*1. Analytical fluorescence spectroscopy*

Steady-state fluorescence measurements were performed with a Fluorolog 3-22 spectrofluorometer (HORIBA Jobin-Yvon) equipped with photon-counting electronics, double-grating excitation and emission monochromators, automated Glan-Thompson polarizers, and a 450-W xenon lamp. Measurements were made either in 4 x 4-mm quartz microcells or in 1 x 1-cm quartz cuvettes with stir disc seated in a thermostated cell holder. For all spectral measurements, samples containing the fluorescent moiety (peptide or extrinsic fluorophore) were recorded in the L-configuration. Endogenous fluorescence measurements were made in parallel with blanks (lacking fluorophore but otherwise identical), which were used for background subtraction to obtain the signal originating solely from the fluorescent molecule. The following steady-state measurements were made:

(a) Endogenous fluorescence from the 2´,6´- dimethyltyrosine (2´,6´-Dmt) moiety of the native SS-31 peptide was measured by emission ($\lambda_{ex}$ = 283 nm; $\lambda_{em}$ = 290 to 400 nm) or excitation ($\lambda_{ex}$ = 260 to 300 nm; $\lambda_{em}$ = 308 nm) scans using 1 nm increments and 2 s integration times. To suppress scattered light from lipid-containing samples and eliminate spectral distortions, in-line polarizers were used in the cross-oriented configuration ($Ex^{pol}$ = 90°, $Em^{pol}$ = 0°) as described (6).

(b) Endogenous steady-state anisotropy (<*r*>) of 2´,6´-Dmt was determined by measuring emission intensities ($\lambda_{ex}$ = 280 nm; $\lambda_{em}$ = 308 nm) at four different polarizer orientations (excitation with vertical polarization, $I_{VV}$ and $I_{VH}$, and excitation with horizontal polarization, $I_{HV}$ and $I_{HH}$). Each sample was background-subtracted with the cognate blank to measure the G-factor ($G = I_{HV}/I_{HH}$) and the steady-state anisotropy $\bar{r} = \frac{I_{VV} - GI_{HV}}{I_{VV} - 2GI_{VH}}$. The steady-state anisotropy of DPH was determined using $\lambda_{ex}$ = 360 nm; $\lambda_{em}$ = 430 nm.

(c) Fluorescence measurements of extrinsic fluorophores were performed by emission scans (1 nm increments with 1-2 s integration times) or by kinetic measurements (0.5-1 s integration times) as follows. Aladan fluorescence from the [ald]SS-31 peptide was measured by emission scans ($\lambda_{ex}$ = 360 nm; $\lambda_{em}$ = 400 to 600 nm) with excitation and emission bandpass at 4 nm. The fluorescence of laurdan and prodan-containing samples was measured by emission scans ($\lambda_{ex}$ = 340 nm; $\lambda_{em}$ = 370 to 600 nm) with excitation and emission bandpass at 2 nm. Emission intensities at 440 nm and 490 nm ($I_{440}$ and $I_{490}$, representing emission maxima of pure $L_\beta$ and $L_\alpha$ phases, respectively) were used to calculate the Generalized Polarization (GP) of laurdan as $GP^{LAU} = \frac{(I_{440} - I_{490})}{(I_{440} + I_{490})}$. The GP of prodan was calculated by the three-wavelength excitation GP (3wGP) (7) as $GP^{PRO} = \frac{R_{12} - 1}{R_{12} + 1}$ where $R_{12} = \frac{I_{420}k}{(I_{480}k - I_{530})}$, $I_{420}$, $I_{480}$, and $I_{530}$ are emission intensities at 420, 480, and 530 nm, and $k$ is a constant independently verified to be 2.8. The fluorescence of ANS was measured by time course ($\lambda_{ex}$ = 380 nm; $\lambda_{em}$ = 460 nm) or emission scan ($\lambda_{ex}$ = 380 nm; $\lambda_{em}$ = 400 to 500 nm) measurements with excitation and emission bandpass at 4 nm. The fluorescence of CG-5N was measured by time course ($\lambda_{ex}$ = 506 nm; $\lambda_{em}$ = 532 nm) or emission scan ($\lambda_{ex}$ = 506 nm; $\lambda_{em}$ = 512 to 562 nm) measurements with excitation and emission bandpass at 2 nm. And the fluorescence of TMRM was measured by time course measurements ($\lambda_{ex}$ = 546 nm; $\lambda_{em}$ = 573 nm) with excitation and emission bandpass at 4 nm.

Time-resolved fluorescence lifetime measurements were performed with a Fluorolog 3-22 instrument equipped with a TCSPC module and pulsed NanoLED light source (peak wavelength 280 nm +/- 10 nm; pulse width <1.2 ns).

*2. Microcalorimetry*

Isothermal titration calorimetry (ITC) measurements were performed as described (8) with a Low Volume Nano-ITC microcalorimeter (TA Instruments, New Castle, DE). Solutions of SS-31 (titrate) and LUVs (titrant) were prepared in degassed buffer, equivalent to that used for liposome preparation. Lipid-into-peptide titrations were performed with 87.5-175 μM SS-31 in the calorimeter cell (volume 170 μl) and LUVs (8 mM total lipid) were injected in aliquots of 2.5 μl (20 total injections) at time intervals of 300 s at 25°C. To account for heats of dilution, experiments were performed by addition of titrant into solutions of buffer only, which were used for baseline subtraction. Data from dilution-corrected and integrated heat flow time courses were fit as Wiseman plots, from which equilibrium binding and thermodynamic parameters ($K_A$, $n$, $\Delta H$, $\Delta S$) were determined by nonlinear regression fits (NanoAnalyze software v. 3.10.0, TA Instruments).

*3. NMR spectroscopy*

NMR data were recorded at 162 MHz on a Bruker AVANCE III 400 MHz WB spectrometer equipped with an HXY probe tuned for $^{1}H$ and $^{31}P$. High power decoupled spectra were acquired with a recycle delay of 2 s, a 90º pulse of 5 μs, and a 50 kHz decoupling field. All $^{31}P$ NMR spectra were externally referenced by assigning the 85% $H_3PO_4$ peak to 0 ppm. Model membranes (LUVs) were measured at 25ºC (total 10,240 scans per sample) with a total lipid concentration of 2 mM lipid in the presence of different concentrations of SS-31 (0, 1:500, 1:50, and 1:5 $[P]:[L]^{eff}$). Isolated mitochondria were measured in minimal buffer (600 mM sorbitol, 20 mM HEPES-KOH, pH 7.5) at 4ºC (total 512 scans per sample) at a total concentration of 70 mg mitochondrial protein $ml^{-1}$ with different concentrations of SS-31 (up to 3 mM) and/or $[CaCl_2]$ (up to 5 mM). Peak fitting and deconvolution was performed using the software dmFit (9).

*4. Small angle x-ray scattering (SAXS)*

SAXS scattering curves (scattered intensity, $I(q)$ as a function of moment transfer, $q$) were recorded at the LiX beamline of National Synchrotron Light Source II (NSLS-II), Brookhaven National Laboratory (BNL) operating at 13 keV. LUV samples (2 mM total lipid) were measured in 1.5 mm diameter quartz flow-through capillaries using a one-second exposure and the $q$-range covered was 0.005 to 3.2 $Å^{-1}$.

*5. Electrokinetic measurements*

Measurements of zeta potential (ζ) of LUVs were performed as we have described (10) using a Zetasizer Nano ZS (Malvern). LUVs (100 μM total lipid) were added to the sample cell (final volume, 1 ml) with different concentrations of SS-31. Values of ζ represent the electrostatic potential at the hydrodynamic shear plane of lipid vesicles based on the measured electrophoretic mobility ($\mu$) by the Helmholtz-Smoluchowski relationship:

$$\xi = \frac{\mu\eta}{\varepsilon_r \varepsilon_0}$$

where η is the viscosity of the solution, $\varepsilon_r$ is the dielectric constant of the aqueous phase, and $\varepsilon_0$ is the permittivity of free space. Calculation of surface potential ($\psi_0$) was based on the distance-dependent decay of electrostatic potential from the membrane surface:

$$\kappa x = \left\{ \frac{\left[\exp\left(ZF\zeta/2RT\right)+1\right]\left[\exp\left(ZF\psi_0/2RT\right)-1\right]}{\left[\exp\left(ZF\zeta/2RT\right)-1\right]\left[\exp\left(ZF\psi_0/2RT\right)+1\right]} \right\}$$

where *R* is the molar gas constant, *T* is the absolute temperature, Z is ionic charge of solution electrolyte, x is the distance from the membrane surface, and κ (the Debye constant) is:

$$\kappa = \left( \frac{2Z^2F^2C_{i\infty}}{\varepsilon_r \varepsilon_0 RT} \right)^{1/2}$$

The electrostatic shear plane was assumed to be at x = 2Å.

*6. Mitochondrial Activity Assays*

Mitochondrial calcium uptake was measured by monitoring the bulk [$Ca^{2+}$] using the cell-impermeant calcium-indicating fluorophore calcium green 5N (CG-5N, prepared as an aqueous stock), adapted from a protocol described for measuring $Ca^{2+}$ uptake into *S. cerevisiae* mitochondria (11). Reactions containing mitochondria (100 μg) in Calcium Flux Buffer (CFB, 300 mM sucrose, 20 mM HEPES-KOH, 2 mM $KH_2PO_4$, 0.5 mg $ml^{-1}$ BSA, pH 7.5) were prepared with the probe (1 μM CG-5N) in the presence or absence of calcium ionophore (5 μM ETH-129) and variable concentrations of $CaCl_2$. Measurements of mitochondrial membrane potential were made using the potentiometric fluorophore TMRM under identical conditions, but in the presence of 50 nM TMRM (prepared as a methanol stock). CG-5N and TMRM measurements were performed as time course measurements in stirred cuvettes with the progressive addition of respiratory substrate (1 mM NADH) and $CaCl_2$ at specified time intervals in the presence or absence of the indicated concentrations of SS peptides.

Oxygen consumption rates of isolated mitochondria were measured using a closed, thermostated chamber equipped with a Clark-type oxygen electrode (Oxygraph Plus System, Hansatech). Reactions were set up with 100 μg of mitochondria in one of two respiration measurement buffers. Standard Respiration Buffer (SRB, 300 mM sucrose, 20 mM HEPES-KOH, 10 mM $KH_2PO_4$, 10 mM KCl, 5 mM $MgCl_2$, 1 mg $ml^{-1}$ BSA, 0.5 mM EDTA, pH 7.5) was used for routine testing of mitochondrial activity and coupling. Minimal Respiration Buffer (MRB, 300 mM sucrose, 20 mM HEPES-KOH, 1 mg $ml^{-1}$ BSA, 10 mM $KH_2PO_4$, pH 7.5) was used for calcium stress analyses. To test the effects of calcium stress and/or SS peptide on mitochondrial respiration, a concentrated stock of mitochondria (1 mg $ml^{-1}$) was preincubated with $CaCl_2$ (1 mM, equivalent to 1 μmol $Ca^{2+}$ per mg mitochondria) and/or SS-31 (54.4 μM, equivalent to 54.4 nmol SS-31 per mg mitochondria) on ice for 20 min prior to polarographic measurements. The effect of calcium stress on mitochondrial respiration was measured by monitoring state 2 respiration for three min following the addition of respiratory substrate (19 mM succinate and 1 mM NADH) and by monitoring respiration without non-catalytic $H^+$ flux through the $F_O$ of ATP synthase for three min following the addition of 14.4 μM oligomycin.

*7. Lipidomics*

Multidimensional mass spectrometry-based shotgun lipidomic analysis of mitochondrial lipids was performed as described (4, 12). In brief, a mixture of internal standards was added to isolated mitochondria for quantitation of phospholipid species based on the content of mitochondrial protein, lipids were extracted using a modified Bligh and Dyer procedure (13), and each lipid extract was reconstituted in 1:1 (v/v) chloroform/methanol at a volume of 200 μl $mg^{-1}$ mitochondrial protein. For shotgun lipidomics, lipid extracts were diluted to a final concentration of ~500 fmol total lipids $\mu l^{-1}$. Mass spectrometric analysis was performed on a triple quadrupole mass spectrometer (TSQ Altis, Thermo Fisher Scientific, San Jose, CA) and a Q Exactive mass spectrometer (Thermo Scientific, San Jose, CA), both of which were equipped with an automated nanospray device (TriVersa NanoMate, Advion Bioscience Ltd., Ithaca, NY) as described (14). Identification and quantification of phospholipid species were performed using an automated software program (15, 16). Data processing (ion peak selection, baseline correction, data transfer, peak intensity comparison and quantitation) was performed as described (16).

*8. Statistical Analysis*

All reported means are the average values of a minimum of three measurements from independent experimental samples. Statistical comparisons were performed by t-test (ns, $P>0.05$; • $P<0.05$; •• $P<0.01$; ••• $P<0.001$; •••• $P<0.0001$).

**C. Molecular dynamics methods**

*1. Simulation methods*

The SS-31 peptide structure was generated by modifying an extended tetrapeptide with sequence Arg-Tyr-Lys-Phe. Coordinates were modified using the VMD Molefacture Plugin (17) to invert the stereochemistry of the N-terminal Arg residue (from L to D) and to replace the 2′

and 6´ hydrogen atoms of the Tyr side chain with methyl groups. Parameters for the 2´,6´-Dmt were modeled after the output parameters of 3´,5´-dimethylphenol after running its structure through ParamChem's CGenFF server (18). The fully parameterized SS-31 peptide was then run through the CGenFF server again to check for energy penalties.

MD simulations were then used to characterize the binding mechanism of SS-31 and to investigate its effects on membrane dynamics. All-atom systems with explicit membrane and solvent were prepared using CHARMM-GUI with the CHARMM-36m forcefield and the TIP3 water model (19-25). Bilayers were generated with molar ratios of TOCL:POPC (20:80), MLCL:POPC (20:80), and POPG:POPC (20:80). Each system contained a total of 300 lipids (150 per leaflet). As done in a previous study (26), the MLCL system was created from the TOCL system by replacing an *sn*-2 acyl chain with a hydroxyl, appending the topology, and using a GROMACS-compatible topology file (.itp) for MLCL. Following the CHARMM-GUI standard protocol for membranes, the bilayer systems were energy-minimized using the steepest-descent algorithm for 5000 steps, followed by canonical ensemble equilibration for 50 ps with 1 fs timestep, and 325 ps of NPT equilibration with a 2 fs timestep accomplished using the semiisotropic pressure coupling scheme and the Berendsen barostat. Position and dihedral restraints were used during equilibration on the lipids to maintain lipid geometry and bilayer morphology. Production simulations of the bilayers alone were each run for 1 μs in an NVT ensemble with a timestep of 2 fs. In our simulations, we define the "upper leaflet" as the side of the membrane exposed to peptides and the "lower leaflet" as the opposing side, which was not exposed to peptides.

Peptide-bilayer systems were constructed by removing solvent from the equilibrated bilayers, placing 10 peptides 1-3 nm away from the upper leaflet headgroup region, and then re-solvating and adding neutralizing sodium ions. The sizes of the systems were ~88,000 atoms for the TOCL systems, ~85,000 atoms for the MLCL systems, and ~73,000 for the POPG systems. The systems were then energy minimized for 5000 steepest descent steps, followed by canonical ensemble equilibration for 50 ps with 1 fs timestep, and 425 ps of NPT equilibration with a 2 fs timestep accomplished using the semiisotropic pressure coupling scheme and the Berendsen barostat. Position and dihedral restraints were used during equilibration on the lipids and peptides to maintain lipid geometry and bilayer morphology and to prevent the peptides from interacting with the bilayers during equilibration. Production simulations were run for 1.6 μs and saved every 50 ps for the GROMACS mean-squared-displacement (MSD) and radial distribution function (RDF) analyses, every 100 ps for GROMACS solvent accessible surface area (SASA) analyses, and every 250 ps for the MDTraj analyses. The first 50 ns of the production simulations were omitted from all analyses except the binding time-dependence (Figs. 5A and S11A). To enforce peptide binding to only the upper leaflet (i.e., to impose an energy penalty on peptides crossing the z-dimension periodic boundary and binding to lower leaflet), two inverted flat-bottom restraints in the Z-direction were placed ~1 nm away from the headgroup region of the lower leaflet. A restraint was placed on the peptides with a force constant of 1000 kJ $mol^{-1}$ $nm^{-1}$ and a radius of 2 nm, which served to prevent peptides from binding to the lower leaflet. A second restraint was placed on the lower leaflet phosphates of POPC with a force constant of 200 kJ $mol^{-1}$ $nm^{-1}$ and a radius of 0.5 nm to prevent the lower leaflet from drifting in the negative z-direction while still allowing for natural membrane deformations.

All simulations were performed using GROMACS 2016 (27, 28). Electrostatic and Lennard-Jones (LJ) interactions were cut off at 1.2 nm, with electrostatics shifted from 0 nm to the cutoff, and LJ interactions shifted from 1.0 nm to the cutoff. All productions runs were simulated in the NPT ensemble using the Parrinello-Rahman coupling scheme, with the temperature maintained at 303.15 K and pressure kept at 1.0 bar with semi-isotropic coupling. The time constants for pressure and temperature couplings were 5.0 and 1.0 ps, respectively, and the compressibility value was set to $4.5x10^{-5}$ $bar^{-1}$. Simulations were performed using periodic boundary conditions in all dimensions and the simulation time step was 2 fs.

*2. Analysis of MD simulations*

From our simulations, we ascertained: (i) binding time-dependence and membrane insertion depths for each residue (**Fig. 5B**, **Fig. S11**); (ii) binding poses of the peptide within the interfacial region (**Fig. 5C**); (iii) radial distribution profiles of sidechain to lipid headgroup for each residue (**Fig. 5D**, **Fig. 12A**) and between lipid headgroups (**Fig. 12C**) for each system; (iv) co-diffusion of Arg and coordinated lipid phosphates (**Fig. S12B**); (v) mean square displacement plots for each lipid with and without SS-31 and their corresponding diffusion coefficients (**Fig. S13**); and (vi) mean solvent accessible surface area measurements of the acyl chain regions for each lipid with and without SS-31 (**Fig. S14**).

The binding time-dependence, membrane insertion depth, and co-diffusion of Arg and coordinated lipid phosphates were analyzed using the MDTraj Python module (29) to process the trajectories and in-house Python scripts to conduct the analysis. The average and standard errors of the mean membrane insertion depths were calculated from selected residues (n=10 for Arg and Lys residues in all three lipid systems; n≥7 for 2´,6´-Dmt and Phe residues in all three lipid systems) which achieved stable positions within the bilayer and remained bound for the remainder of the trajectory. Radial distribution profiles were generated using the GROMACS *gmx rdf* function (27, 28). Mean square displacements were calculated using the *gmx* msd function in GROMACS (27, 28) by tracking the phosphates of POPG and POPC and the central glycerol C2´ carbon of TOCL and MLCL. Area per lipid calculations were made using the GROMACS *gmx energy* function to extract the average x-y dimensions from three separate segments of the trajectory and dividing by the number of lipids per leaflet. Bilayer thicknesses were calculated using in-house scripts to determine differences in the average z-position of lipid phosphates in the upper and lower leaflets of each system. To calculate the average lipid phosphate z-positions, each lipid phosphate was treated as an independent sample to calculate mean z-positions over each trajectory. Solvent accessible surface area (SASA) measurements were calculated using the *gmx sasa* function in GROMACS (27, 28), which uses the double cubic lattice method (30). For the SASA analyses, we defined the acyl chain region of a lipid as the carbon and hydrogens below the ester carbon. The upper leaflet of each bilayer that contained the SS-31 peptides was used for both the radial distribution profiles, the mean square displacement, and SASA analyses for the "with SS-31, upper leaflet" condition. The lower leaflet of each bilayer in "+SS-31" trajectories, but that was not in contact with peptides, was used for the diffusion coefficients and the SASA analyses for the "with SS-31, lower leaflet" condition. Both leaflets of the systems without any peptide were used for the mean square displacement analysis, radial distribution profiles (normalized for lipid count), and SASA analyses (normalized for lipid count) for the "without SS-31" condition. Diffusion coefficients were calculated from the mean square displacement analysis and fit to the linear portion of the curve (10 to 50 ns) using the Einstein relation:

$$\lim_{t\to\infty}\left\langle \left\| r_i(t) - r_i(0) \right\|^2 \right\rangle_{i\in A} = 6D_{\mathrm{A}}t$$

The average diffusion coefficients (Fig. S13B) and average acyl chain SASA measurements (Fig. S14) were calculated from the averages over three equal intervals of ~515 ns in the systems with peptide, and over three equal intervals of ~315 ns for the systems without peptide. 95% confidence intervals were calculated from the standard error of the averages from each interval of the respective trajectories. All images of the systems were created using VMD (17). All figures were created using the Matplotlib Python module (31).

### D. Quantitative analysis of peptide binding isotherms and surface electrostatics

*1. Fitting saturation binding curves to obtain parameters n and $K_D$*

Models for the binding of SS-31 to membranes are based on the independent and reversible binding of free peptide in solution ($P_f$) to an accessible lipid binding site ($M_f$) that is composed of *n* free lipids ($L_f$) to form a peptide-membrane complex (*PM*):

$$[P]_f + [M]_f \rightleftarrows PM \qquad \text{Eq. S1}$$

Here, the total number of peptide binding sites is related to the lipid concentration by $[M] = [L]_0/n$, where $[L]_0$ is the total concentration of accessible lipids. The equilibrium dissociation constant ($K_D$, in units of *M*) associated with this binding is:

$$K_D = \frac{[P]_f[M]_f}{[PM]}$$

Binging isotherm data were fit to quadratic expansions of the Langmuir adsorption isotherm equation (32) to extract equilibrium binding data as follows:

For experiments in which a fixed concentration of *L* was titrated with *P*, binding data were fit by the following equation:

$$\theta = \frac{([P]_0 + [L]_0/n + K_D) - \sqrt{([P]_0 + [L]_0/n + K_D)^2 - (4[P]_0[L]_0/n)}}{2([L]_0/n)} \qquad \text{Eq. S2}$$

In this equation, $\theta$ is the fractional saturation, $[P]_0$ and $[L]_0$ are the total *P* and accessible *L* concentrations, respectively, *n* is the lipid:peptide stoichiometry, and $K_D$ is the equilibrium dissociation constant. In these experiments, the total number of peptide binding sites ($[L]_0/n$) is constant and $\theta$ is a measure of the fraction of lipid binding sites bound to peptide. Note that when binding saturation data are fit to Eq. S2, $K_D$ is expressed in terms of molarity of peptide *binding sites* (not individual lipids); therefore, the term $nK_D$ expresses the affinity of peptide for lipid monomers.

For experiments in which a fixed concentration of *P* was titrated with *L*, binding data were fit by:

$$\theta = \frac{(n[P]_0 + [L]_0 + K_D) - \sqrt{(n[P]_0 + [L]_0 + K_D)^2 - (4n[P]_0[L]_0)}}{2(n[P]_0)} \qquad \text{Eq. S3}$$

In this equation, the number of accessible lipids required to bind all peptide ($n[P]_0$) is constant and θ is a measure of the fraction of peptides bound. Example data and analysis of data fits to Langmuir isotherms are shown for peptide titration (**Fig. S5 A,B**) and lipid titration (**Fig. S5 D,E**).

*2. Scatchard analysis*

As an independent means of analysis, binding data were also evaluated by Scatchard analysis (33) as follows. Given the definition of the equilibrium association constant ($K_A$, in units of $M^{-1}$) in terms of peptide-lipid complex ($[PL]$), free peptide ($[P]_f$) and free lipid ($[L]_f$):

$$K_A = \frac{[PL]}{[P]_f[L]_f} \qquad \text{Eq. S4}$$

This equation can be recast to include bound peptide ($[P]_b$) and bound lipid ($[L]_b$). In the case of peptide titrations (with constant lipid), given that $[L]_b/n = [PL]$ and that $[P]_f = [P]_0 - [P]_b$, Eq. S4 is rearranged such that:

$$\frac{[L]_b}{[L]_f} = nK_A[P]_0 - nK_A[P]_b \qquad \text{Eq. S5}$$

Given the relationship $[P]_b = [L]_b/n$, Eq. S5 becomes

$$\frac{[L]_b}{[L]_f} = nK_A[P]_0 - K_A[L]_b$$

which can be rearranged as

$$\frac{[L]_b}{[L]_f[P]_0} = K_A\left(n - \frac{[L]_b}{[P]_0}\right) \qquad \text{Eq. S6}$$

Therefore, a plot of $[L]_b/[L]_f[P]_0$ *versus* $[L]_b/[P]_0$ yields a line with slope of $-K_A$, a y-intercept of $nK_A$, and an x-intercept of $n$ (**Fig. S5C**).

In the case of lipid titrations (with constant peptide), given that $[P]_b = [PL]$ and that $[L]_f = [L]_0 - [L]_b$, Eq. S4 is rearranged such that:

$$\frac{[P]_b}{[P]_f} = K_A[L]_0 - K_A[L]_b \qquad \text{Eq. S7}$$

Given the relationship $[L]_b = n[P]_b$, Eq. S7 becomes

$$\frac{[P]_b}{[P]_f} = K_A[L]_0 - K_A n[P]_b$$

which can be rearranged as

$$\frac{[P]_b}{[P]_f[L]_0} = K_A\left(1 - n\frac{[P]_b}{[L]_0}\right) \qquad \text{Eq. S8}$$

Therefore, a plot of $[P]_b/[P]_f[L]_0$ *versus* $[P]_b/[L]_0$ yields a line with slope of $-nK_A$, a y-intercept of $K_A$, and an x-intercept of $n^{-1}$ (**Fig. S5F**).

*3. Partition coefficients*

Peptide binding isotherms from lipid titrations were independently fit to obtain the molar partition coefficient $K_P$ as described (34), in accordance with the equation

$$F = \frac{F_0 + K_P \gamma_L [L]_0 F_L}{1 + K_P \gamma_L [L]_0} \qquad \text{Eq. S9}$$

in which $F$ is the measured fluorescence intensity at a given lipid concentration, $[L]_0$; $F_0$ and $F_L$ are, respectively, the peptide fluorescence intensities in the absence of lipid (water only) and the fluorescence when completely lipid-bound (determined from $F$ vs. $[L]_0$ plots); and $\gamma_L$ is the mol fraction-weighted average of lipid molar volume. The partial molar volumes used in this equation were $\gamma_{POPC}$ = 0.7564 $M^{-1}$, $\gamma_{POPG}$ = 0.7238 $M^{-1}$, and $\gamma_{TOCL}$ = 1.4453 $M^{-1}$ from published values (35-37). The corresponding value for MLCL is not available, so we made the assumption that $\gamma_{MLCL} = \gamma_{TOCL}$.

*4. Quantitative analysis of surface electrostatics*

The electrostatic profile within the diffuse double layer of our model membranes was evaluated as we have described (10) using Gouy-Chapman-Stern formalism (38, 39). The relationship between the charge density at the membrane surface ($\sigma$, in C $m^{-2}$) and the electrostatic potential of the aqueous phase at the surface of the membrane ($\psi_0$, in V) is described by a variation of the Gouy-Chapman equation:

$$\sigma = \frac{\psi_0}{|\psi_0|}\left[2\varepsilon_r\varepsilon_0 RT\sum_i C_{i\infty}\left(\exp\left(\frac{-Z_i F\psi_0}{RT}\right)-1\right)\right]^{1/2} \qquad \text{Eq. S10}$$

where $\varepsilon_r$ is the dielectric constant, $\varepsilon_0$ is the permittivity of free space, $R$ is the molar gas constant, $T$ is the absolute temperature, $C_{i\infty}$ is the bulk concentration of ionic species $i$, $F$ is the Faraday constant, and $Z_i$ is the formal charge on ionic species $i$. The distribution of ions within the interfacial region and normal to the membrane surface is determined by Boltzmann statistics, wherein the relationship between $C_{i\infty}$ and its surface concentration ($C_{0,i}$) is given by:

$$C_{0,i} = C_{i\infty}\exp\left(\frac{-Z_i F\psi_0}{RT}\right) \qquad \text{Eq. S11}$$

The adsorption of ions to the charged membrane surface is described by a Langmuir isotherm:

$$\sigma = \frac{\sigma^{\max}}{1 + K_M C_{0,i}} \qquad \text{Eq. S12}$$

where $\sigma^{max}$ is the maximum charge density of the membrane surface, determined by the mole fraction-weighted lipid headgroup formal charge and cross-sectional areas, and $K_M$ is the association constant that describes the adsorption of solution ions.

The simultaneous evaluation of Eq. S10-S12 allows for the determination of $\psi_0$ as a function of $\sigma^{max}$, $C_{i\infty}$, and $K_M$. This analysis was used to evaluate $\psi_s$ (surface potential) under conditions of different lipid composition and bulk ionic strength (Fig. 3B) and to evaluate SS-31 binding isotherms based on measurements of zeta potential (Fig. 3C).

*5. Nonlinear least squares fitting*

Evaluation of binding data to simultaneously solve for *n* and $K_D$ (Eq. S2 and S3), to solve for $K_P$ (Eq. S9), and to evaluate membrane electrostatics (Eq. S10-S12) was performed as described (10) using *Wolfram Mathematica 10*. Note that some binding experiment series in this study included conditions in which binding isotherms did not reach saturation (e.g., Fig. 3A), rendering binding responses that could not, individually, be adequately fit by Eq. S2 or Eq. S3. In such cases, we performed a global analysis of all binding curves within a given experiment set, including those that did reach saturation, with the assumption that all binding curves will asymptote toward the same maximal value.

### <u>SI Results</u>

#### A. Spectral characterization of SS-31 in solution (*pertaining to Fig. S3*)

SS-31 contains two fluorescent side chains (**Fig. 1A**): a phenylalanine (Phe) at position four and a 2´,6´- dimethyltyrosine (2´,6´-Dmt, tyrosine [Tyr] with methyl groups at the $\delta^1$ and $\delta^2$ positions of the aromatic ring) at position two. We selected the 2´,6´-Dmt moiety as a potential reporter for SS-31 membrane interactions because (i) Tyr has a significantly higher extinction coefficient and quantum yield than Phe, and (ii) fluorescence measurements can be designed to selectively measure Tyr fluorescence in the presence of Phe, because absorption and emission spectra of the two side chains are sufficiently separated (Phe absorbing and emitting at lower wavelengths) (40). The aromatic side chain of tyrosine contains a phenol chromophore with fluorescence properties that are sensitive to its microenvironment (41-43) and may be used to measure peptide and protein interactions with membranes (44-46).

Initially we characterized the spectral features of the 2´,6´-Dmt side chain of SS-31 in comparison with L-Tyr in aqueous buffer (**Fig. S3**). We first measured absorbance and fluorescence spectra of L-Tyr and SS-31. In comparison with free tyrosine, SS-31 displayed a more structured and red-shifted absorbance with a concurrent shift in the wavelength of maximal fluorescence excitation ($\lambda_{ex}^{SS\text{-}31}$ = 281 nm) (**Fig. S3A**, compare red and blue traces). We next measured the relationship between fluorescence yield and concentrations of L-Tyr or SS-31. The emission intensity of SS-31 linearly increased with concentration up to 100 μM (**Fig. S3B**, red) and its emission was reduced in comparison with free L-Tyr (**Fig. S3B**, blue). These steady-state measurements were consistent with shorter measured fluorescence lifetime of SS-31 in solution ($<\tau^{SS\text{-}31}>$ = 1.07 ns) in comparison with L-Tyr ($<\tau^{Tyr}>$ = 3.33 ns). To address the solvent accessibility of the 2´,6´-Dmt side chain of SS-31, we measured dynamic quenching of L-Tyr and SS-31 by the collisional quenching agent acrylamide. Stern-Volmer analysis showed a reduced solvent accessibility of the 2´,6´-Dmt in SS-31 ($K_{SV}^{SS\text{-}31}$ = 4.8 $M^{-1}$; $k_q$ = 4.49 x $10^9$ $M^{-1}$ $s^{-1}$) in comparison with free L-Tyr ($K_{SV}^{Tyr}$= 17.1 $M^{-1}$; $k_q$ = 5.14 x$10^9$ $M^{-1}$ $s^{-1}$) (**Fig. S3C**). Finally, to confirm that the Phe residue of SS-31 did not contribute to our measured signals, we compared emission spectra of SS-31 with SS-20, a peptide variant with two Phe residues (**Fig. 1B**). Under the conditions used for our fluorescence measurements, SS-20 is spectrally silent (**Fig. S3D**, note that the intensity of SS-20 emission overlaps with the abscissa of the graph).

Thus, taken together, the data of Fig. S3 reveal the following: (i) compared with L-Tyr, the 2´,6´-Dmt side chain of SS-31 in solution is less solvent accessible, possibly due to steric shielding by neighboring side chains; (ii) in comparison with L -Tyr, SS-31 has a decreased fluorescence lifetime and emission intensity, likely due in part to proximity of the ε-$NH_3$ group of lysine, which is a strong quencher of phenolic fluorescence (47); and (iii) within μM concentrations, the intrinsic fluorescence emission of SS-31 is in the linear dynamic range.

**B. Fluorescence-based analysis of SS-31 membrane interactions** (*pertaining to Fig. S4*)

We next addressed the ability of the 2´,6´-Dmt side chain of SS-31 to serve as a reporter for membrane interaction of the peptide (**Fig. S4**). Membrane binding measurements of Figs. S4A and B were performed by incubating 20 μM of SS-31 with large unilamellar vesicles (LUVs) in a low- or high-salt buffer (buffer with no added salt or with 100 mM KCl added, respectively) for 20 min, followed by spectroscopic analysis. Binding of SS-31 to these model membranes is believed to be restricted to the external lipid leaflet in these studies. We therefore quantified added lipid as the effective lipid concentration ($[L]^{eff}$) of the outer leaflet, which for LUVs of this size is half of the total lipid (0.5x$[L]^{tot}$) as described (48). In the experiments of Fig. S4 A and B, the $[L]^{eff}$ was 100 μM; hence, the peptide to effective lipid concentration ($[P]:[L]^{eff}$) was 1:5. For simplicity, we base all data analyses using this outer leaflet lipid concentration; in cases where the absolute total amount of lipid used, it will be reported as $[L]^{tot}$.

LUVs consisted of either pure POPC or 20 mol% anionic lipids (TOCL, MLCL, or POPG) in a host background of POPC. Emission scans of SS-31 in the absence and presence of LUVs revealed three important features (**Fig. S4A**). First, although incubation of SS-31 with membranes did not cause a spectral shift in 2´,6´-Dmt fluorescence, it did effect an increase in emission intensity. We attribute this phenomenon to a transfer of the 2´,6´-Dmt moiety from the aqueous environment to a bilayer-inserted state. This is not likely due to a simple change in the dielectric of the side chain microenvironment, but rather to other factors that include: (i) a change in solvent hydrogen bonding with the phenolic hydroxyl group, (ii) a reversal of quenching by another side of the peptide (e.g., the lysine at position three), and (iii) specific polar interactions between 2´,6´-Dmt and lipid headgroups. Second, the magnitude of LUV-induced emission increase is lipid-dependent. As shown previously (1, 49-51), SS-31 has a strong propensity to bind bilayers containing lipids with anionic headgroups, but binds very weakly to bilayers with a net zero charge. Our results show that the emission increase in the presence of LUVs containing only the zwitterionic POPC is very weak, and is much more robust in the presence of LUVs composed of 20 mol% anionic lipid. Further, binding to bilayers containing the dianionic TOCL and MLCL is significantly higher than it is to bilayers containing the monoanionic POPG. Finally, the LUV-induced emission increase is significantly reduced in the presence of high salt, particularly for bilayers with anionic lipids. This is explained by charge shielding of the anionic lipid headgroups by solution electrolytes, which decreases the interfacial surface charge (38, 39). Together, these results show that: (i) the intrinsic emission of the 2´,6´-Dmt side chain of SS-31 increases with membrane binding, and (ii) that SS-31 interaction with lipid bilayers is sensitive to the bilayer surface charge density and to the presence of electrolytes in solution.

To compare the solvent accessibility of the 2´,6´-Dmt reporter when SS-31 is free in solution and when SS-31 is membrane-bound, we performed dynamic quenching experiments in the presence and absence of LUVs (**Fig. S4B**). Note that due to the sensitivity of SS-31 membrane binding to the ionic strength of the solution, the use of a charged collisional quenching agent would have untoward effects on binding; we therefore used acrylamide (formal charge of zero) as a dynamic quencher in these experiments. In these experiments, the measured fluorescence was corrected for the absorbance of exciting light by acrylamide as described (52). For SS-31 in the absence of membranes, the 2´,6´-Dmt moiety was solvent exposed, as expected, under low salt ($K_{SV}$ = 5.7 $M^{-1}$) and high salt ($K_{SV}$ = 5.4 $M^{-1}$) conditions. In the presence of LUVs under low salt conditions, acrylamide accessibility was almost negligible ($K_{SV}$ = 0.15 $M^{-1}$, 0.02 $M^{-1}$, and 0.21 $M^{-1}$ for LUVs containing 20% TOCL, MLCL, and POPG, respectively). The presence of high salt reduced the LUV-dependent protection from acrylamide for vesicles with 20% TOCL and MLCL ($K_{SV}$ = 1.20 $M^{-1}$ and 1.04 $M^{-1}$, respectively) and more so for vesicles with 20% POPG ($K_{SV}$ = 2.19 $M^{-1}$). These results are consistent with the salt-sensitivity of membrane binding observed in our emission scans.

Reduction of the surface charge density of liposomes containing anionic lipids by the addition of cationic species can induce vesicle aggregation. To address whether the presence of peptides induced aggregation of our model membranes, we performed turbidity measurements over a range of peptide and lipid molar ratios (**Fig. S4C,D**). Optical absorbance is an established means of monitoring changes in vesicle size (e.g., by flocculation, fusion, micellezation, etc.) due to soluble factors such as membrane-active peptides (53). First, we measured absorbance changes by titrating solutions with LUVs (20% TOCL, MLCL, or POPG) with increasing [SS-31] (**Fig. S4C**). For comparison, we performed parallel experiments in which we titrated LUVs with divalent cations. Below a threshold [SS-31]:$[lipid]^{tot}$ ratio of 1:1 (or a $[Ca^{2+}]$:$[lipid]^{tot}$ ratio of 10:1), we observed no change in absorbance; above these thresholds, absorbance increases were observed within a narrow concentration range. These results are consistent with previous studies measuring $Ca^{2+}$-induced aggregation of cardiolipin-containing liposomes (54). Second, to account for any effects of lipid concentration, we measured the absorbance of solutions containing increasing concentrations of LUVs containing 20% TOCL in the absence or presence of SS-31 (**Fig. S4D**). For this assay we analyzed a range of LUV concentrations (up to $[lipid]^{tot}$ = 250 μM) corresponding to the range of concentrations used in the majority of our SS-31 binding experiments. We also used a [SS-31]:$[lipid]^{tot}$ of 1:10, a peptide concentration approaching binding saturation for this lipid composition. Based on this assay, we did not observe peptide-induced aggregation at any LUV concentration.

Taken together, we conclude from the analyses of Fig. S4 that: (i) when SS-31 is membrane-bound, the 2´,6´-Dmt side chain increases its emission yield, confirming that this is an excellent reporter for SS-31 bilayer interactions; (ii) the SS-31 bilayer interaction is strongly dependent on membrane surface charge and solution ionic strength; and (iii) over the range of [SS-31] and [LUV] typically used throughout this study, there is no observable aggregation of our model membranes.

**C. Quantitative analysis of SS-31 membrane binding isotherms** (*pertaining to Fig. S5*)

In **Fig. S5**, we illustrate our approaches for fitting SS-31-bilayer binding measurements with simulated data sets. By one approach, SS-31 binding to lipid bilayers was analyzed as a Langmuir-type isotherm. For experiments performed by titrating a fixed concentration of lipid

($[L]_0$) with peptide ($[P]_0$]), simulated binding curves ($\theta$ vs. $[P]_0$, fit with Eq. S2) originating from different $n$ and $K_D$ values are shown in Fig. S5A. The total amount of bound peptide over each titration range ($[P]_b$ vs. $[P]_0$) for each associated case is shown in Fig. S5B. By comparison, experiments performed by titrating a fixed concentration of peptide ($[P]_0$) with lipid ($[L]_0$) for the same $n$ and $K_D$ values produce binding curves ($\theta$ vs. $[L]_0$, fit with Eq. S3) shown in Fig. S5D, with the total amount of bound peptide over each titration range shown in Fig. S5E. Corresponding Scatchard plots for peptide (Eq. S6) and lipid (Eq. S9) titrations are shown in Fig. S5, panels C and F, respectively.

**D. SS-31 titration time course analysis and ITC** (*pertaining to Figs. S6 to S8*).

SS-31 binding isotherms measured by peptide titration were generally conducted by time course measurements in which a solution of LUVs at a fixed lipid concentration was titrated with SS-31. Such experiments typically entailed total lipid concentrations up to 250 μM (i.e., up to $[L]^{eff}$ of 125 μM) with the progressive addition of SS-31 in 2 μM increments. This assay is based on the measurable increase in emission from the 2´,6´-Dmt side chain upon the transition from bulk aqueous solution to the lipid bilayer (Fig. S4). To enhance the difference between emission intensity of the soluble vs. bound peptide, these assays included a fixed concentration of acrylamide in the aqueous medium (55).

Representative time courses of SS-31 titrations are shown in **Fig. S6**. As expected, these traces show that the SS-31 emission jumps are high when lipid is available for binding, and become measurably reduced at a point during the titration when lipid binding sites become saturated. Beyond this point, fluorescence increases with further peptide addition are small and linear with added SS-31 because all added peptide either remains soluble or dynamically replaces pre-bound peptide.

The time course data from these measurements can be analyzed as shown in **Fig. S7**. Plotting the stepwise emission intensities as a function of $[P]$:$[L]^{eff}$ and determining the best linear fits to pre- and post-binding site saturation phases of the titrations, the value of the abscissa at the intersection of the two lines corresponds to the $[P]$:$[L]^{eff}$ at saturation (i.e., 1/$n$) (**Fig. S7 A,B**). A summary of these analyses (**Fig. S7C**) reveals the following. Under low salt conditions, SS-31 binds to LUVs with 20 mol% TOCL with an $n$ value of ~5 (roughly one peptide bound per five lipid molecules, or one peptide per cardiolipin molecule on the external leaflet). For LUVs containing 10 mol% TOCL or 20 mol% POPG under low salt conditions, the $n$ value roughly doubles to values of ~12 and 11, respectively. Together, these results suggest that the number of lipids that constitute an SS-31 binding site at saturation is formally determined by the net charge at the bilayer surface (roughly one peptide per two lipid headgroup charges). This analysis also reveals the effect of ionic strength on the binding stoichiometry, as the *n* value for LUVs with 20 mol% TOCL increases to ~9 in the presence of high salt. Hence, the peptide-lipid stoichiometry is a function of surface charge density ($\sigma$), which is determined by both the intrinsic charge of the lipids and the electrolytes in the interfacial region.

Notably, the analysis of Fig. S7 works well under high affinity interactions when added peptide essentially binds quantitatively to available lipid sites and the pre- and post-saturation regimes of the titrations can be easily delineated. However, under conditions in which the binding affinity is reduced (e.g., lower lipid headgroup charge and/or higher ionic strength), the difference between the two regimes becomes more difficult to delineate and binding parameters must be evaluated by curve fits using nonlinear least squares analysis (Figs. 2 and 3).

Microcalorimetry experiments were conducted by the progressive addition of LUVs with different lipid composition (20% TOCL, MLCL, or POPG in a POPC background or 100% POPC as a control) at increments of 10 nmol total lipid to solutions of peptide. The starting concentrations of SS-31 in the sample cell were varied ([SS-31] = 175 μM for CL-containing membranes; [SS-31] = 87.5 μM for PG-containing membranes) so that a complete binding curve could be obtained over the injection course. Representative heat flow time courses corrected for heats of dilution (**Fig. S8**) show that the peptide-bilayer interactions are exothermic and saturable. ITC data were analyzed as Wiseman plots (**Fig. 2D**) to obtain $K_A$, *n*, $\Delta H$ and $\Delta S$.

**E. Model membrane surface electrostatics and SS-31 binding** (*pertaining to Fig. S9*)

*1. Effects of ionic strength on SS-31 binding*

To address the salt-dependent reversibility of SS-31 membrane interactions, we measured the relative SS-31 membrane binding when increasing concentrations of salt were added before or after peptide was bound to membranes (**Fig. S9A**). We found no difference in the ionic strength-dependent decrease in SS-31 bilayer interaction whether salt was added before or after peptide binding, quantified as either emission intensity (Fig. S9A, *left*) or anisotropy (Fig. S9A, *right*). To substantiate this observation, we performed kinetic measurements of relative SS-31 binding with progressive addition of monovalent or divalent cations (**Fig. S9B**). The addition of salt caused rapid fluorescence-detected desorption of SS-31 from the bilayer surface that was commensurate with cation charge. We conclude that the binding of SS-31 to model membranes is highly reversible in a manner that is dependent on solution salt concentration. Hence, based on GCS models, cations "compete" with SS-31 for binding by two mechanisms: (i) electrostatic shielding, and (ii) specific complexation with lipid functional groups containing formal (phosphates) or partial (e.g., carbonyl oxygens) negative charges.

*2. Effect of membrane binding on SS-31 formal charge*

Our SS-31 binding analyses (Fig. 2) suggest that when peptide is maximally bound to the surface of an anionic bilayer, there is a near balance of charge between peptide formal charge (+3) and headgroup phosphates (one peptide binds per every ~1.5 CL/MLCL molecules or per every ~3 PG molecules). However, our ζ profiles of Fig. 3C suggest that a measureable negative charge density exists among the headgroup phosphates even when SS-31 is maximally bound to membranes. This observation could originate from phenomena such as partial charge neutralization of SS-31 upon membrane binding (as found for other polybasic peptides, e.g., (56)) and/or changing conformations of ionized lipid headgroups that alter the electric potential sensed at the slip plane. To begin to address this question, we sought to determine the formal charge on bound SS-31 from the ζ profiles of Fig. 3C, based on the additive charge densities of: ionized lipids ($\sigma^{max}$), bound SS-31 ($\sigma^{SS\text{-}31}$, taken from peptide titration binding isotherms (Fig. 2A)), and adsorbed solution ions ($\sigma^{ion}$, based on $K_M$, the intrinsic association constant of ion complexation determined from LUVs in the absence of peptide). This analysis returned a formal charge on SS-31 of +1.7 and +2.3 for membranes containing 20% TOCL and POPG, respectively. Based on this analysis, bound SS-31 is approximately 30% charge-neutralized, which could, for example, result from the deionization of basic groups in the microenvironment of the bound state. Such partial charge neutralization of bound SS-31 could contribute to the persistence of negative surface potentials in model membranes even when peptide is maximally bound to the surface. Ongoing work in our group is aimed at using NMR- and electrokinetic-based approaches to address this question by examining the structural basis of the side chain-lipid headgroup interactions and the effect of peptide on headgroup orientation.

**F. Assays for peptide-induced structural polymorphism of model membranes** (*pertaining to Fig. S10*).

*1. SAXS measurements with model membranes*

SAXS measurements of lipid samples yield x-ray diffraction patterns that convey information about polymorphic state and bilayer dimensions (57). SAXS profiles for LUVs of different lipid composition in the presence of different molar ratios of SS-31:lipid are shown in **Fig. S10A**. These plots show scattering intensity $I(q)$ as a function of momentum transfer $q$, where $q = (4\pi/\lambda) \sin\theta$, λ is the wavelength of the x-ray beam, and θ is half of the angle between the incident beam and scattered radiation. Scattering profiles of our LUVs containing anionic lipid show a broad peak between $q$ = ~0.05 and 0.25 $Å^{-1}$, typical of unilamellar vesicles. By contrast, profiles of LUVs containing POPC only (not shown) display a superposition of this broad peak and two peaks corresponding to first- and second-order Bragg reflections (with q spacings of $n2\pi/d$, where n is the peak order and d is the repeat distance accounting for the bilayer and intervening water), suggesting the presence of some oligolamellar vesicles.

We fit our SAXS data using the Global Analysis Program (GAP) (58, 59), kindly provided by Dr. Georg Pabst (University of Graz). Fitting was performed using the equation

$$I(q) = \frac{(1 - N_{\mathrm{diff}}) S(q) [F(q)]^2 + N_{\mathrm{diff}} [F(q)]^2}{q^2} \qquad \text{Eq. S13}$$

where $N_{\mathrm{diff}}$ is the fraction number of uncorrelated bilayers per scattering domain, $F(q)$ is the bilayer form factor, and $S(q)$ is the interbilayer structure factor based on Caillé theory. From these fits, the following parameters are shown beneath each corresponding plot: $d$ (the lamellar repeat distance), $N_{\mathrm{diff}}$, $z_{\mathrm{H}}$ (the Gaussian distribution center of the polar heads), and $\eta$ (the fluctuation parameter related to bilayer bending rigidity). These values show no systematic trends that correlate with different molar amounts of SS-31.

*2. $^{31}$P ssNMR measurements with model membranes*

Phosphorous NMR allows for the analysis of lipid phosphate headgroups, yielding specific spectral lineshapes characteristic of gel or liquid crystalline lamellar, inverted $H_{II}$ and cubic phases (60). As a complement to our SAXS analyses, we used $^{31}$P solid state NMR to test for any peptide-induced changes in our model membranes. The major phospholipids used throughout this work all have main transition temperatures below freezing ($T_m^{POPC}$ = -2ºC; $T_m^{TOCL}$ = -8.4ºC; $T_m^{POPG}$ = -2ºC) (61-63). Therefore, at the temperatures used throughout this study, LUVs prepared as binary mixtures of these miscible lipids are expected to yield $^{31}$P NMR spectra with powder patterns consistent with the liquid crystalline (L$\alpha$) phase.

To characterize the $^{31}$P NMR spectra of the LUV samples used in this study, we hydrated a lipid film (20 mol% TOCL, 80 mol% POPC) in aqueous buffer and processed the dispersion in three ways: (i) preparation of multilamellar vesicles (MLVs); (ii) processing of MLVs by four freeze/thaw (liquid $N_2$ / 35ºC water bath) cycles; and (iii) extrusion through 0.1 $\mu$m polycarbonate membrane to produce LUVs. Analysis of these samples by $^{31}$P ssNMR (**Fig. S10B**) revealed that MLVs produced asymmetric lineshapes (low-field shoulder and high-field peak, CSA 54.4 ppm) characteristic of axially symmetric phospholipid in a lamellar phase. By comparison, MLVs subject to freeze/thaw cycles show some evidence of motional averaging (CSA 55.0 ppm), and samples that were extruded to produce unilamellar vesicles of uniform diameter (LUVs, ~100 nm) showed complete averaging of the NMR spectrum, yielding a single isotropic peak. This observation is consistent with greater rotational mobility/tumbling of the smaller and more homogeneous LUVs compared with larger MLVs with stacked lamellae (64). We conclude that the LUVs used with SS-31 titration (Fig. 4B) are indeed ~100 nm lamellar vesicles, although their size causes partial motional averaging of their spectral lineshapes.

*3. Membrane-bound fluorescent probes*

DPH is a rod-shaped fluorophore that partitions near the center of lipid bilayers and serves as an anisotropy probe for lipid dynamics and order due to the sensitivity of its rotational diffusivity to the microviscosity of its surroundings (65) (**Fig. S10C**). The steady-state anisotropy of DPH ($\langle r \rangle^{DPH}$) increases with microviscosity (i.e., with decreased fluid dynamics of the hydrocarbon tails) and can report changes in membrane phase state, as occurs with thermotropic transitions between lameller gel (Lβ) and liquid crystalline (Lα) phases. DMPC and TMCL are short acyl chain (14:0) variants of PC and CL whose Lβ to Lα phase transition temperatures (Tm) are within an experimentally tractable range ($Tm^{DMPC}$ = 24°C and $Tm^{TMCL}$ = 47°C). LUVs composed of a binary mixture of 20% TMCL and 80% DMPC and containing DPH yielded temperature-dependent maximum and minimum $\langle r \rangle^{DPH}$ values of 0.265 and 0.084 (corresponding to Lβ and Lα phases, respectively) with a sharp transition in $\langle r^{DPH} \rangle$ centered at 28.5°C, corresponding to a cooperative Lβ to Lα phase change (**Fig. S10D**). Having established the range of $\langle r \rangle^{DPH}$ expected of this PC/CL system, we then used DPH anisotropy measurements to address the SS-31 response of binary lipid systems of longer chain and more unsaturated lipids used throughout this study (20% TOCL or 20% POPG in a host background of POPC). With increasing SS-31, we observed low $\langle r \rangle^{DPH}$ values that corresponded to the Lα phase for both TOCL- and POPG-containing membranes, with no measurable effect of peptide, even at saturating concentrations (**Fig. S10E**). We conclude that SS-31 binding does not affect the rotational diffusion of the DPH probe, consistent with the binding of the peptide in the headgroup region with negligible effect on hydrocarbon chain dynamics.

Laurdan and prodan are interface-localized probes that contain the same naphthalene-based solvatochromic fluorescent moiety, but partition at different depths within in membrane due to differences in their nonpolar substituents: prodan (with a propionyl tail) resides near the level of phospholipid glycerol groups, whereas larudan (with a lauryl tail) resides deeper, near the level of the *sn*-1 carbonyl (66) (**Fig. S10C**). Emission scans of laurdan- and prodan-containing LUVs composed of 20 mol% TOCL or POPG in a POPC background in the presence of different SS-31:[Lip]$^{eff}$ ratios are shown in **Fig. S10F**. TOCL-containing membranes revealed modest, but consistent, peptide-dependent and saturable changes in the spectra of the probes. Specifically, laurdan fluorescence showed an SS-31-dependent decrease in $I_{490}$ vs. $I_{440}$ (equivalent to a blueshift), causing an increase in $GP^{LAU}$; prodan fluorescence showed an SS-31-dependent decrease in $I_{480}$ and $I_{520}$ relative to $I_{420}$, causing an increase in $3wGP^{PRO}$. These changes correlate with decreases in probe hydration (reduced probe dipolar relaxation), indicative of higher order/lipid packing in the interfacial region (67-69). Measurable peptide-dependent increases in $GP^{LAU}$ and in $3wGP^{PRO}$ were also observed for POPG-containing membranes, but were much reduced in comparison with TOCL-containing bilayers. Note that due to differences in the SS-31 binding capacities of POPG- and TOCL-containing bilayers (Fig. 2), the titration range of [SS-31] for POPG bilayers was half of that for TOCL bilayers in these experiments.

**G. Molecular dynamics analyses** (*pertaining to Figs. S11-S14*).

*1. Characterization of general membrane structural parameters*

For our simulated membrane systems, the average area per lipid for bilayers of different composition were (mean ± SE): 20% TOCL, 75.49 Å$^2$ ± 0.04; 20% MLCL, 71.91 Å$^2$ ± 0.01; and 20% POPG, 64.03 Å$^2$ ± 0.11. Bilayer thickness was calculated as the phosphate-to-phosphate (P-P) distance between leaflets in the presence and absence of SS-31 in the upper leaflet. Average P-P distances were (mean ± SE): 20% TOCL (no SS-31), 3.98 nm ± 0.04; 20% TOCL (with SS-31), 3.95 nm ± 0.04; 20% MLCL (no SS-31), 3.90 nm ± 0.04); 20% MLCL (with SS-31), 3.87 nm ± 0.04; 20% POPG (no SS-31), 3.90 nm ± 0.04; 20% POPG (with SS-31), 3.87 nm ± 0.05. These measures of area per lipid and bilayer thickness are consistent with MD work from our group (26, 70) and others (e.g., (71)). To complement these P-P distances that were calculated over the bilayer ensemble, we also calculated distances between upper leaflet phosphates and the bilayer C.O.M. (phosphate z-positions) for individual lipids (**Fig. S11A**). Given the acyl chain asymmetry of MLCL, we refer to the headgroup phosphate on the side of the lipid with the full acyl chain complement as "P1" and to the phosphate on the side lacking an *sn*-2 chain as "P3" (corresponding to the "native " and "lyso" phosphates, $P_N$ and $P_L$, respectively in (26)). Although the headgroup phosphates of TOCL are equivalent, we distinguish them as "P1" and "P3", corresponding to the side from which the acyl chain was cleaved in generating MLCL from TOCL. As expected, these phosphate z-positions are roughly half the distances of our calculated P-P bilayer thicknesses. Notably, however, the presence of

SS-31 caused a slight but consistent reduction in P-P bilayer thickness, as well as reduction in the z-position specifically of anionic lipid phosphates (Fig. S11A). We discuss the potential origin of this peptide-dependent effect on phosphate distances below.

*2. Measurement of side chain insertion depth*

In each MD binding trajectory (**Fig. S11B**), basic side chains revealed rapid membrane association with z-coordinates that remained generally consistent until the end of the simulation. By comparison, the 2´,6´-Dmt side chains, and to a greater extent the Phe side chains, displayed rapid transitions between the water- and nonpolar core-exposed states until becoming stably bound in the acyl chain region. Residue-specific membrane insertion depths (**Fig. S11C**) were determined for bound peptides, which were defined as those having both aromatic groups buried in the acyl chain region. For each bound peptide in the three different bilayer systems, we quantified the average z-coordinate of each side chain ($z^{pos}$) based on landmark atoms (the D-Arg Cζ atom; the 2´,6´-Dmt Oη atom; the Lys Nζ atom; or the Phe Cζ atom). Specifically, the average and standard errors of the mean membrane insertion depths were calculated from selected residues (n = 10 for Arg and Lys residues and n ≥ 7 for 2´,6´-Dmt and Phe residues in all three lipid systems) that achieved stable positions within the bilayer and remained bound throughout the trajectory. Position variability was quantified as the 95% confidence intervals of $z^{pos}$ values for each side chain. We observed no significant side-chain specific differences in penetration depth among the three lipid systems.

*3. Radial distribution profiles: lipid distributions around SS-31 residues*

The proximity of different lipid headgroups around each side chain landmark atom (defined above) was measured by lateral (x-y plane) radial distribution profiles (**Fig. S12A**). For basic side chains, we observed three distinct radial shells of phosphate groups: a primary density located ~4Å from the side chain atom, as well as secondary and tertiary densities located ~6Å and ~9-10Å from the side chain atom, respectively. The primary peak, consistent with the Arg-phosphate and Lys-phosphate hydrogen bonding distances (72), shows two features for TOCL-containing membranes. First, both basic side chains show a strong preference for TOCL phosphates over POPC phosphates in this primary shell, consistent with the greater accessibility of the charged phosphate group in the former (73). Second, the peaks are higher for Arg than they are for Lys, consistent the more extensive hydrogen bonding contacts possible by the guanidino group relative to the primary amine, as also seen with MD simulations of penetratin (74). Compared with TOCL, the magnitude of the primary phosphate shell around Arg and Lys is reduced for POPG, but the general trends are similar. Interestingly, however, for MLCL-containing bilayers, the primary phosphate shell around Arg does is not significantly enriched in MLCL phosphates over POPC phosphates. This is likely attributable, in part, to disruption of intermolecular phosphate contacts due to hydrogen bond contacts with the hydroxyl at the *sn*-2 position of the lyso side of MLCL (26). We also note that this reduction in Arg-phosphate contacts in MLCL-containing bilayers appears to be compensated by enhanced contacts between Lys and MLCL phosphates. The secondary and tertiary phosphate shells likely consist primarily of: (i) phosphates on the same headgroup as the phosphate in the primary density (for di-phosphate TOCL and MLCL) and/or (ii) phosphates corresponding to those in the primary density of the alternate basic residue of a given peptide. As a complement to this analysis, co-diffusion between Arg and lipid phosphates from six different peptides in a 100 ns block of a given simulation (**Fig. S12B**) supports preferential interaction between this basic side chain and TOCL phosphates. The hydrogen bond network that stabilizes the interaction between the Arg guanidino group and phosphates illustrates the potential for Arg to mediate interaction with multiple lipid headgroup phosphates (75).

*4. Effects of SS-31 on lipid bilayer properties*

Lipid-to-lipid radial distribution profiles (based on phosphate-phosphate radii) revealed two primary peaks in the absence of peptide, corresponding to successive lipid shells centered around 6Å and 9Å, respectively (**Fig. S12C** *black traces*). In comparison with this radial distribution based on lipid self-association without peptide, the presence of SS-31 caused a slight but measurable enhancement in the local concentration of anionic lipid (**Fig. S12C**, *red traces*). This effect most likely originates from localization of anionic lipids in the vicinity of bound peptide by specific interactions with basic side chains (Fig. 12A,B). We also note that the observed SS-31-dependent decrease in the average z-positions of anionic (but not zwitterionic) lipid phosphates (Fig. S11A) is likely attributable, at least in part, to the polar interactions that these phosphates make with the basic side chains of the peptide. That is, the presence of peptide causes anionic lipid phosphates to reside deeper in the bilayer (by ca. 0.4-0.5Å), perhaps reflecting a balance between maintaining the position of the headgroup within the aqueous interface and the interaction between the phosphate and the basic side chain.

Based on the lateral mean square displacement of lipid phosphates, we observed marked SS-31-dependent decreases in the diffusivity ($D_{xy}$) of all tested lipids (**Fig. S13**). Specifically, the presence of SS-31 caused a decrease in $D^{TOCL}$ from 0.52 x $10^{-7}$ $cm^2$ $s^{-1}$ to 0. 34 x $10^{-7}$ $cm^2$ $s^{-1}$, a decrease in $D^{MLCL}$ from 0.53 x $10^{-7}$ $cm^2$ $s^{-1}$ to 0.35 x $10^{-7}$ $cm^2$ $s^{-1}$, and a decrease in $D^{POPG}$ from 0.74 x $10^{-7}$ $cm^2$ $s^{-1}$ to 0. 48 x $10^{-7}$ $cm^2$ $s^{-1}$. The presence of peptide also caused a decrease in $D^{POPC}$ for all systems tested. Furthermore, we compared the diffusivity of lipids between upper (peptide-accessible) and lower (peptide non-accessible) leaflets in simulations containing SS-31. Although the lower leaflet displayed trends toward SS-31-dependent decrease in $D_{xy}$ in these simulations, they did not significantly differ from the no-peptide simulations. Therefore, we conclude from this analysis that the binding of SS-31 causes a marked reduction in lipid lateral diffusion on the leaflet to which peptide is bound.

Our analysis of the solvent accessible surface area (SASA) of acyl chains in our simulations (**Fig. S14**) revealed two key features. First, using increased SASA as a measure of disturbance in lipid packing (acyl chain exposure), we found that the presence of TOCL caused greater local packing defects than did MLCL, and that both CL variants caused greater packing defects than POPG. These results, consistent with our previous work (26), are attributable to the molecular geometries of these anionic lipids (TOCL, and to a lesser extent, MLCL, have lower effective headgroup volumes relative to acyl chain volume occupancy, than does POPG). Second, this analysis showed peptide-dependent decreases in solvent accessibility of all tested lipids; specifically, $SASA^{TOCL}$ decreased from 12.0 $nm^2$ to 9.8 $nm^2$, $SASA^{MLCL}$ decreased from 9.4 $nm^2$ to 7.8 $nm^2$, and $SASA^{POPG}$ decreased from 5.8 $nm^2$ to 4.7 $nm^2$. Consistent with our analyses of lipid diffusion, SASA was only measurably decreased on the leaflet to which SS-31 was bound. These results suggest that the binding of SS-31 has the effect of reducing the accessibility of solvent (and solvated ions) to acyl chains that would otherwise be exposed to the interface, largely by lipid packing defects.

**H. Assays for measuring $Ca^{2+}$ interactions with model membranes and mitochondria** (*pertaining to Figs. S17-S19*).

*1. Characterization of fluorescence-based analyses of calcium dynamics and $\Delta\psi_m$*

CG-5N is a membrane-impermeant probe whose fluorescence intensity increases upon binding calcium. To properly evaluate the responses of CG-5N under the different experimental conditions used in this study, we first measured SS peptide-dependent changes in probe emission under conditions used for experiments with model membranes (Fig. 7B-D) and under conditions used for experiments with isolated mitochondria (Fig. 7E-G). In these control experiments, calcium and SS peptides were added in concentrations commensurate with the respective assays of Fig. 7.

Given the potential for some peptides to bind metal ions (76), we first addressed the potential effects of SS-20 and SS-30 on CG-5N fluorescence. In the minimal buffer used for the measurement of model membranes supplemented with calcium, the addition of SS-20 caused a nearly negligible decrease in CG-5N emission in comparison with dilution effects of vehicle-only titration, whereas the addition of SS-31 caused a slightly greater decrease in CG-5N emission (**Fig. S17A**). Notably, however, the quenching effect of both peptides was significantly less than that of the metal ion chelator EDTA, which in these measurements was added in molar amounts 2.5x lower than peptide. The main conclusion from the analyses shown in Figs. 7B-D is that SS peptides bind to anionic lipid bilayers in the headgroup region, thereby shifting the equilibrium binding of $Ca^{2+}$ ions away from coordination at the membrane and toward bulk solution. This conclusion is predicated on the assumptions that CG-5N accurately reports [$Ca^{2+}$] in the bulk solution and that SS peptides do not significantly influence bulk [$Ca^{2+}$] in a manner other than by competing for $Ca^{2+}$ binding at the bilayer surface. Our control tests shown in Fig. S17A do indicate that, under these experimental conditions, SS peptides may have slight effects on CG-5N spectral properties (e.g., through a dynamic quenching mechanism) and on the bulk [$Ca^{2+}$]. However, they do not counter the conclusions from Figs. 7B-D for the following reasons: (i) If SS peptides have a slight quenching effect on CG-5N fluorescence, this effect is *opposite* to what is observed in Figs. 7C-D (that peptides show an increase CG-5N emission due to increased bulk [$Ca^{2+}$]; hence, any untoward quenching effect is likely negligible under these conditions; (ii) If SS-31 has the effect of chelating bulk phase $Ca^{2+}$ ions that is significant under our experimental conditions, the responses in Figs. 7C-D would be different between SS-31 and SS-20; the fact that they are nearly identical indicates that any putative chelation behavior by SS-31 is very unlikely to have an influence on these results.

The buffer used for calcium measurements with mitochondria (CFB, 20 mM HEPES, 300 mM sucrose, 2 mM potassium phosphate, 0.05% BSA, pH 7.5, supplemented with variable concentrations of $CaCl_2$) was adapted from a recent study on $Ca^{2+}$ uptake into yeast mitochondria (11). By comparing CG-5N emission intensity in CFB buffer with and without BSA, we observed that the presence of BSA caused a robust (10.7-fold) reduction in CG-5N fluorescence. We attribute this to the ability of albumin to bind $Ca^{2+}$ (as well as other metal) ions (77). We evaluated the response of the CG-5N probe to reagents used in our mitochondrial calcium transient measurements (**Fig. S17D**). The most notable results of these time traces are as follows. First, vehicle-only controls for additions of respiratory substrate NADH and calcium ionophore ETH-129 had minor effects on CG-5N emission in the absence (sample 1) and presence (sample 2) of mitochondria. Second, the presence of NADH itself increased CG-5N emission (samples 3 and 5). We attribute this NADH-dependent increase in CG-5N emission to the presence of $Ca^{2+}$ in stock concentrations of NADH obtained from the vendor (Sigma N8129) because the presence of EDTA suppressed this NADH-dependent fluorescence increase (not shown). Importantly, this effect of respiratory substrate on CG-5N emission did not confound the analysis of time courses and calcium transients of Fig. 7E-G because: (i) all samples were given identical concentrations of NADH, and (ii) all peptide treatments were performed prior to spectral analysis and fluorescence normalization allowed for direct comparison of all samples. Third, because *S. cerevisiae* mitochondria lack a calcium uniporter, in the absence of ETH-129 (sample 3), the addition of calcium resulted in an increase in CG-5N emission that was sustained over time (i.e., calcium remained in the bulk phase). Fourth, when mitochondria were provided respiratory substrate (NADH) to generate a membrane potential, addition of ETH-129 resulted in a decay of CG-5N emission (sample 5). This was due to the membrane potential-dependent uptake of calcium that was present in the buffer system containing isolated mitochondria. Also in sample 5, the addition of calcium results in a transient response in which the [$Ca^{2+}$] of the bulk solution immediately increases, followed by the electrophoretic uptake into the matrix, manifest as a return of CG-5N emission to the base value. The latter shows the complete uptake of solution calcium into energized mitochondria.

TMRM is a potentiometric probe that is used to measure relative transmembrane potential ($\Delta\psi_m$) of isolated mitochondria. Under the conditions used in this study, TMRM was analyzed in quench mode such that upon establishment of the $\Delta\psi_m$, this lipophilic cationic dye accumulates in mitochondria in accordance with a Nernstian distribution and emission intensity decreases. The response of TMRM to $\Delta\psi_m$ is shown in **Fig. S17E**: addition of respiratory substrate causes a decrease in TMRM emission as the membrane potential is established, and dissipation of the $\Delta\psi_m$ by the $K^+$ ionophore valinomycin causes an increase in the TMRM signal. Under our measurement conditions, the presence of SS peptide did not measurably affect the magnitude of the $\Delta\psi_m$ established in isolated mitochondria.

*2. NMR- and polarography-based analyses of mitochondrial calcium stress*

Our protocols to evaluate the $^{31}P$ NMR spectra of isolated mitochondria were adapted from previous studies (78-81). Isolated mitochondria were suspended at high concentration (70 mg $ml^{-1}$) in a buffer compatible with NMR measurements (600 mM sorbitol; 20 mM HEPES-KOH, pH 7.5, consistent with that used in (81)). For each sample, NMR measurements (512 total transients) were taken over 26 min with temperature maintained at 4°C. We first compared the $^{31}P$ spectra of isolated mitochondria and hydrated dispersions of mitochondria-extracted lipids (**Fig. S19A**). The lineshape of mitochondrial lipids revealed a powder pattern consistent with a lamellar bilayer of lipids with axial symmetry, with a CSA span of 31 ppm and a high-field peak at -12 ppm. The lineshape of our mitochondrial samples revealed multiple peaks, including an isotropic signal ("Iso") and three distinct high-field peaks ("L1-L3"). The large isotropic peak likely originates from molecules undergoing fast reorientation on the NMR timescale, including NTPs, inorganic phosphate, and soluble phosphoproteins. The mitochondrial L2 peak coincides with the peak from our extracted lipids, suggesting that this peak corresponds to a lipid bilayer signal. Mitochondrial peak L3, shifted ~10 ppm upfield from the L2 lamellar component, may correspond to motionally-restricted lipids associated with membrane proteins.

We used NMR analysis as a means of measuring the ability of SS-31 to mitigate stress caused by high levels of calcium. To this end, our highly concentrated samples of mitochondria isolated from WT *S. cerevisiae* were pre-incubated with or without 600 μM SS-31 on ice for 1h. This corresponds to a peptide concentration of 8.5 nmol SS-31 per mg mitochondrial protein, which is comparable to previous studies (e.g., (82)) in which 10-100 μM SS-31 was used with isolated mitochondria at ~0.4 mg $ml^{-1}$ (25 to 250 nmol SS-31 per mg mitochondrial protein). Samples were then incubated with vehicle only or up to 5 mM $CaCl_2$ (i.e., up to ~700 nmol $Ca^{2+}$ per mg mitochondrial protein) followed by NMR analysis. Compared with control samples that did not receive SS-31 or $CaCl_2$ treatment, incubation with SS-31 did not cause a major disruption of the $^{31}P$ spectra, although we did observe a slight shift of the L2 peak to a more shielded position (~1.0 ppm) in the presence of SS-31, whereas the other peaks did not change (**Fig. S19B**). Addition of $CaCl_2$ resulted in the complete loss of the L3 component of our spectra, regardless of the presence of SS-31 (compare **Fig. S19B** and **Fig. 7G**). However, SS-31 did offset the calcium-dependent decay of the L2 component (**Fig. 7G-H**).

Parallel measurements of mitochondrial oxygen consumption were based on conditions similar to our NMR measurements, but with the following differences. First, mitochondria received higher concentrations of SS-31 and $CaCl_2$ (54.4 nmol SS-31 and 1 μmol $Ca^{2+}$ per mg mitochondrial protein, respectively). Second, respirometry measurements were made in a buffer (Minimal Respiration Buffer) with reduced osmoticum (300 mM sucrose) and supplemented with 1 mg $ml^{-1}$ BSA and 10 mM $KH_2PO_4$, pH 7.5. Oxygen consumption rates in this buffer and in the absence of calcium stress were higher than normal (~280 nmol $O_2$ $min^{-1}$ $mg^{-1}$). To confirm that this elevated rate was due to the buffer conditions and not our isolated mitochondria, parallel respirometry measurements made in our Standard Respiration Buffer showed reduced state 2 oxygen consumption rate (~190 nmol $O_2$ $min^{-1}$ $mg^{-1}$) (**Fig. 19C**).

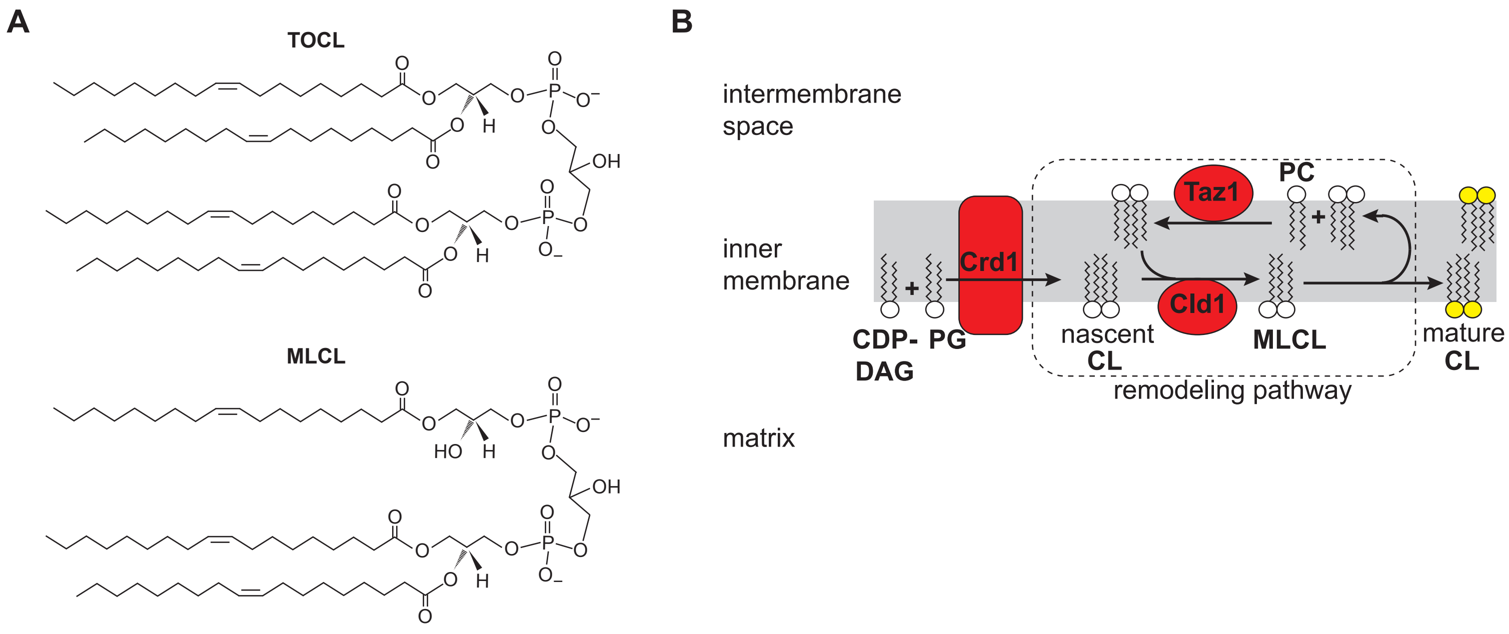


**Supplementary Figure S1. Cardiolipin structure and biogenesis**

**A.** Chemical structures of tetraoleoyl cardiolipin (TOCL) and monolysocardiolipin (MLCL).

**B.** Cardiolipin biosynthesis and remodeling in yeast. The committed step of CL synthesis is the condensation of phosphatidylglycerol (PG) and CDP-diacylglycerol (CDP-DAG) by cardiolipin synthase (Crd1) to produce nascent CL. The remodeling cycle that produces mature CL is initiated by the lipase Cld1, which removes an acyl chain to generate MLCL. The acyl-CoA independent transacylase tafazzin (Taz1) then mediates the transfer of an acyl chain from a donor phospholipid to regenerate tetra-acyl CL. This remodeling pathway yields CL species that are enriched in unsaturated fatty acids.

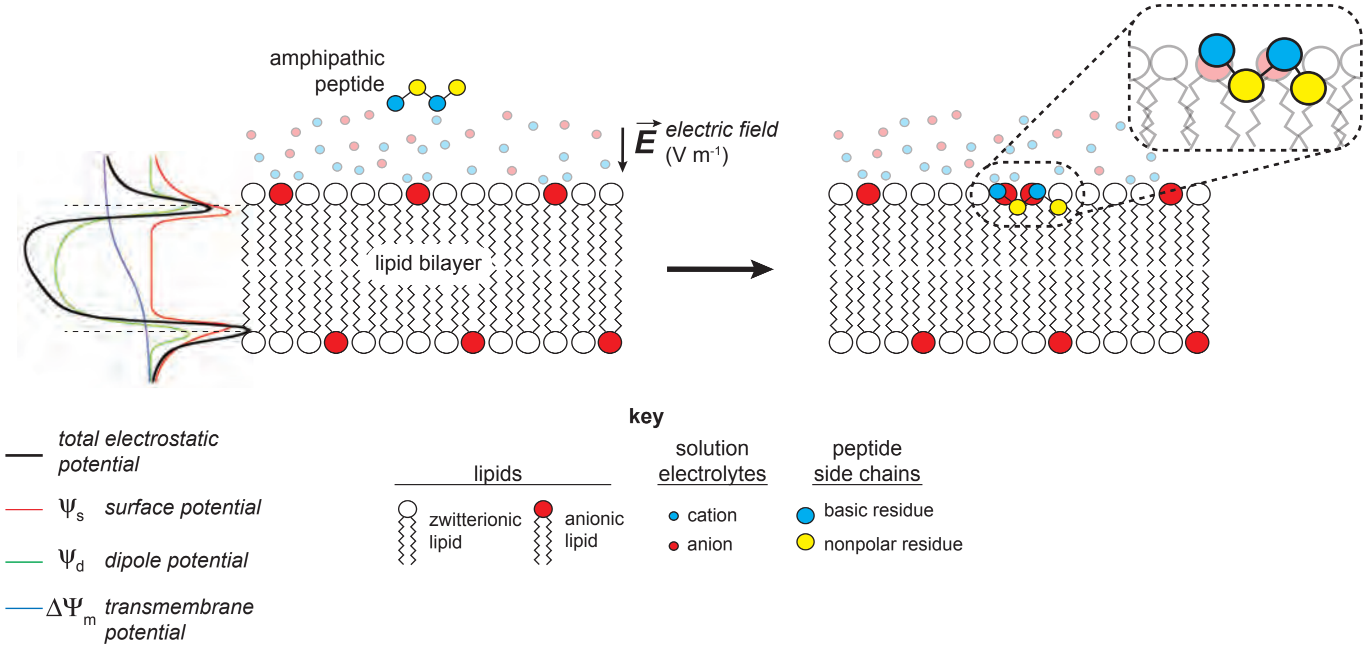


**Supplementary Figure S2. The binding of amphipathic peptides to anionic membranes**

The electrostatic potential of biomembranes consists of three superimposed potentials: the surface potential ($\psi_s$, arising from the net charges in the membrane interface); the dipole potential ($\psi_d$, from intramolecular dipole moments of interfacial chemical groups and water molecules), and the transmembrane potential ($\Delta\Psi_m$, from charge imbalance of aqueous solutions on either side of the membrane).

Peptides containing both basic and nonpolar side chains bind to lipid bilayers with a net negative charge *via* electrostatic and hydrophobic interactions. A lipid bilayer can be characterized by the surface charge density ($\sigma$, C $m^{-2}$), based on the number of formal charges imparted by ionized headgroups per membrane surface area. $\sigma$ is related to surface potential ($\psi_s$, mV) by Gouy Chapman-Stern formalism. The surface charge of anionic biliayers creates a strong electric field (***E***) that attracts positively charged ions and molecules (governed by Boltzmann statistics), creating a diffuse double layer with an enrichment of cations around near the bilayer surface. Cations can electrostatically shield the negative charge density of the bilayer and may undergo complexation with lipid headgroups with an appreciable association constant ($K^{ion}$). An amphipathic peptide such as SS-31 is attracted to the bilayer surface by long-range Coulombic forces originating from the lipid surface charges. This has the effect of accumulating such basic peptides at the bilayer surface at concentrations significantly larger than the bulk peptide concentration. At the bilayer, the peptide basic side chains can form electrostatic interactions with anionic lipid headgroups and the nonpolar side chains can intercalate into the acyl chain region, stabilized by the hydrophobic effect.

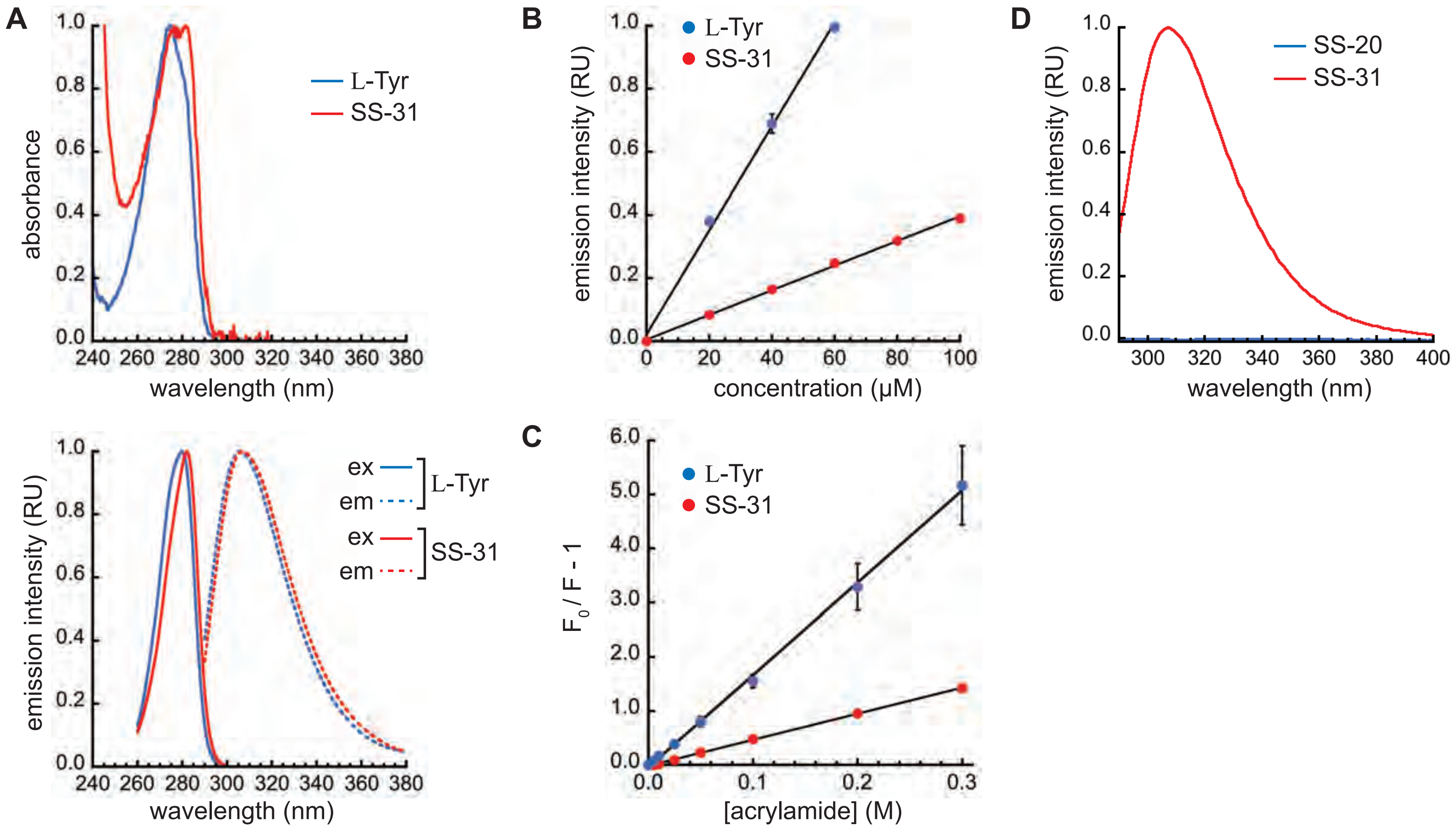


**Supplementary Figure S3. Fluorescence characterization of SS-31 in aqueous buffer**

**A.** Normalized absorption spectra (upper panel) and excitation/emission spectra (lower panel) of 20 μM L-Tyr (blue) and 20 μM SS-31 (red). All traces are signal averages of n=3 independent samples.

**B.** Emission intensity of increasing concentrations of L-Tyr (blue) and SS-31 (red) (n=3 ± SD). Lines represent linear fits to the data (R>0.99 for both L-Tyr and SS-31 regressions).

**C.** Stern-Volmer plots of fluorescence quenching of L-Tyr (blue) and SS-31 (red) by acrylamide (n=3 ± SD). Lines represent fits to the Stern-Volmer equation $F_0/F-1=K_{SV}$[acrylamide] (R>0.99 for both L-Tyr and SS-31 regressions).

**D.** Normalized emission spectra of 20 μM SS-31 (red) and 20 μM SS-20 (blue).

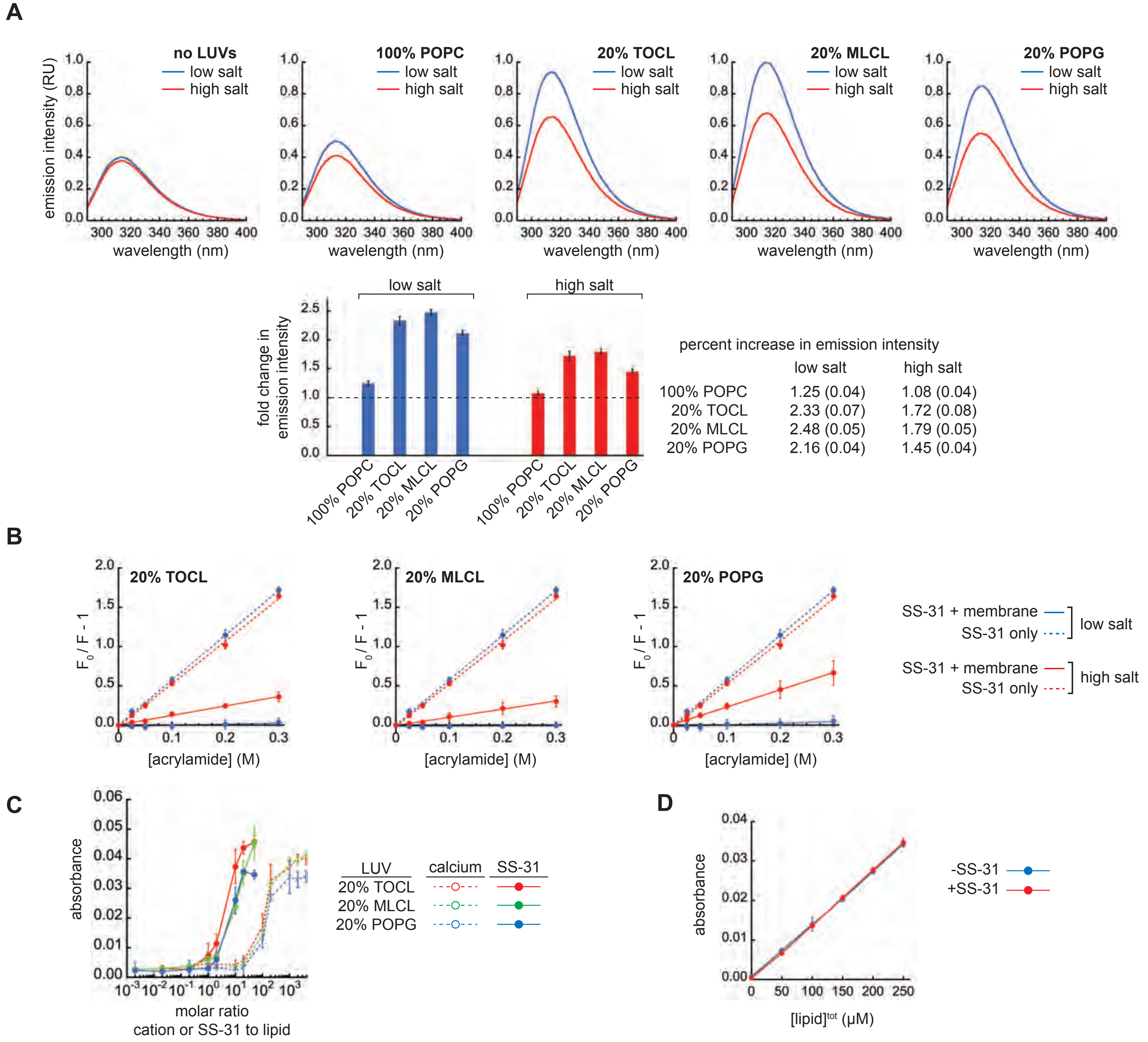


**Supplementary Figure S4. Fluorescence characterization of SS-31 membrane interactions**

**A.** *Above*, normalized emission scans of 20 µM SS-31 ($\lambda_{ex}$ = 281 nm, $\lambda_{em}$ = 290 to 400 nm) in solution (no LUVs) and in the presence of LUVs composed of 100% POPC, or containing 20 mol% TOCL, MLCL, or POPG in a background of POPC as indicated. Measurements were made under low salt (blue) or high salt (red) conditions. Traces are signal averages of n=6 independent samples. *Below*, fold increase in SS-31 emission for lipid-containing samples relative to the respective (low or high salt) emission in the absence of LUVs. Values shown are means (n=6 ± SD) determined from the emission scans above.

**B.** Stern-Volmer plots of acrylamide fluorescence quenching of 20 µM SS-31 in solution (dashed lines) or in the presence of LUVs containing 20 mol% TOCL, MLCL, or POPG as indicated (solid lines) under low salt (blue) or high salt (red) conditions (n=3 ± SD). Lines represent fits to the Stern-Volmer equation $F_0/F-1=K_{SV}$[acrylamide] (R>0.99 for all regressions). Note that data for SS-31 in solution (dashed line fits) are from a single set of experiments but are shown in all panels for comparison.

**C,D.** Absorbance-based analysis of liposome aggregation. LUVs were incubated with the indicated salts or peptide followed by measurements of absorbance at 440 nm. **C**) LUVs ([lipid]$^{tot}$ = 25 µM) with the indicated lipid composition were preincubated with $CaCl_2$ (“calcium”) or peptide (“SS-31”) at different molar ratios followed by turbidity measurements (n=3 ± SD). **D**) LUVs ([lipid]$^{tot}$ = 0 to 250 µM) were preincubated with or without SS-31 at a 1:10 molar ratio (peptide : [lipid]$^{tot}$) followed by turbidity measurements (n=3 ± SD).

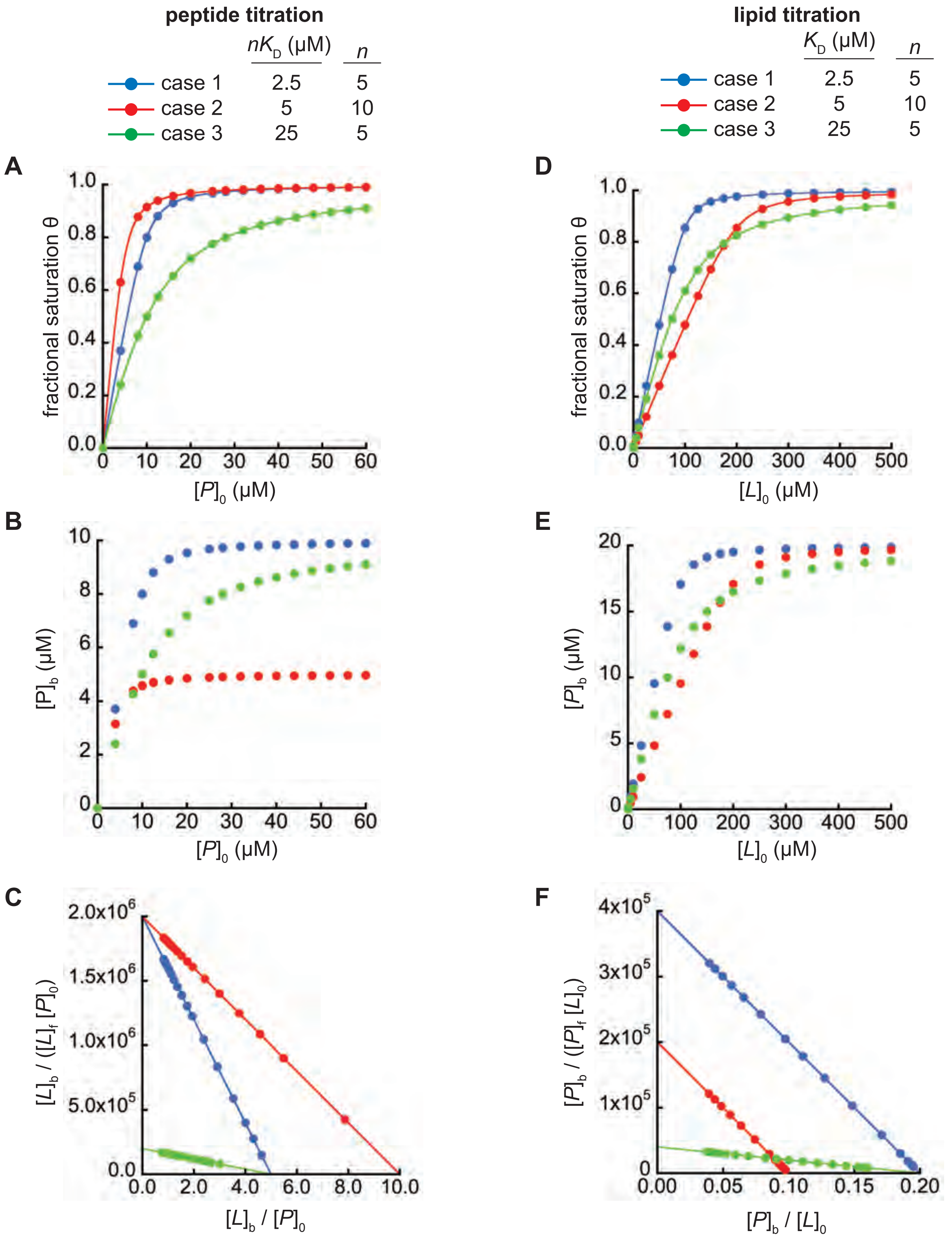


**Supplementary Figure S5. Quantitative analysis of peptide membrane binding**

Simulated peptide-bilayer binding data are shown for three scenarios: case 1 (blue: high affinity, low *n*), case 2 (red: mid affinity, high *n*), and case 3 (green: low affinity, low *n*). (A-C): titration of a fixed concentration of lipid ($[L]_0$=50 μM) with peptide ($[P]_0$, up to 60 μM); (D-F): titration of a fixed concentration of peptide ($[P]_0$=20 μM) with lipid ($[L]_0$, up to 500 μM).

**A,D.** Binding plots showing fractional saturation derived from Eq. S2 (panel A) or Eq. S3 (panel E)

**B,E.** Plots of bound peptide as a function of titrant *P* (panel B) or *L* (panel E)

**C,F.** Scatchard analyses derived from Eq. S6 (panel C) or Eq. S8 (panel F)

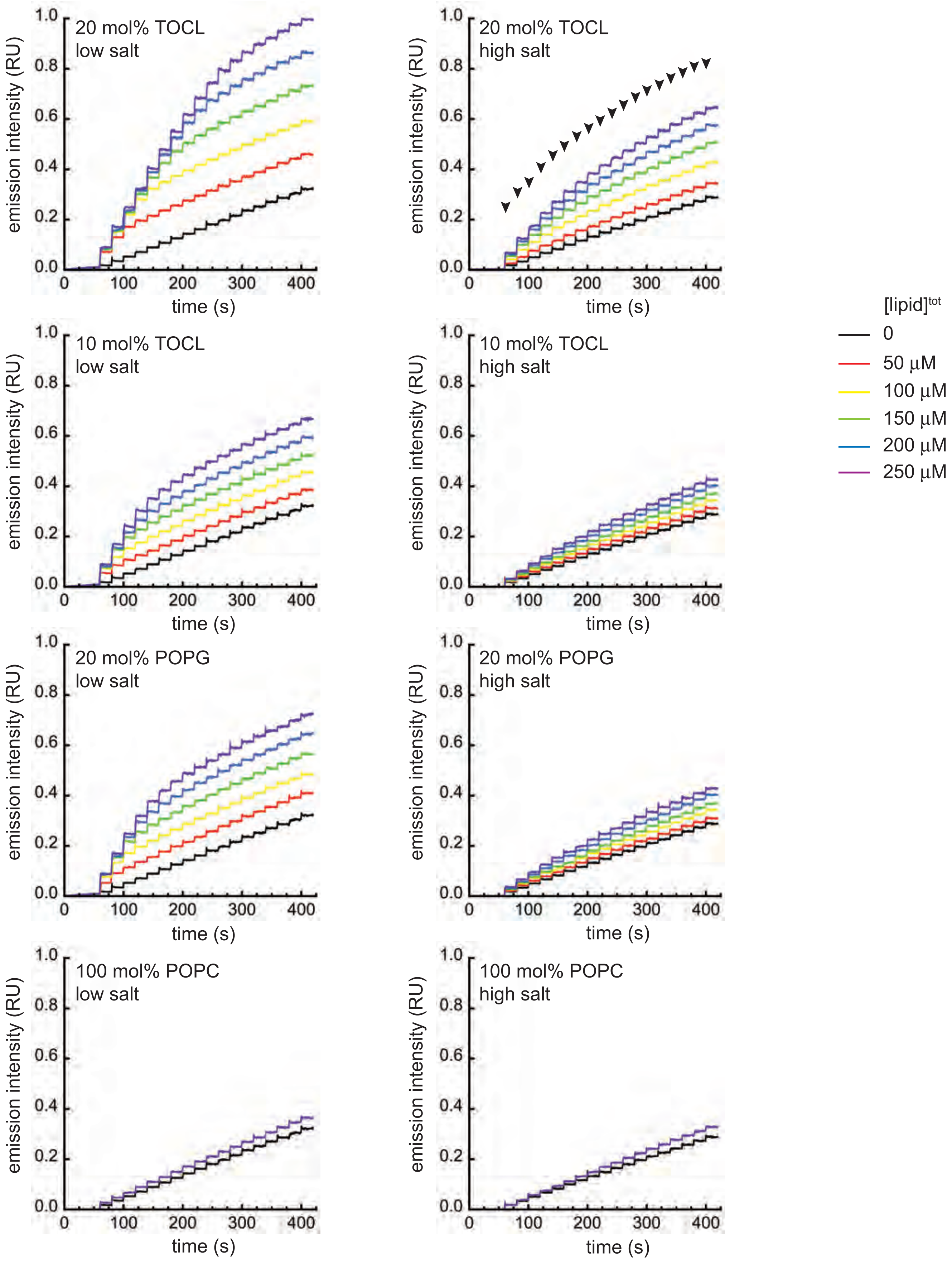


**Supplementary Figure S6. Kinetic measurements of SS-31 addition to LUVs.**
Titration of different concentrations of LUVs ($[\text{lipid}]^{tot}$ = 0, 50, 100, 150, 200, and 250 μM as indicated) with SS-31 at 2 μM increments. Peptide injections commenced at t=60 s and continued at 20 s intervals for a total of 18 injections ($[\text{SS-31}]^{final}$ = 36 μM). Injection points are displayed as arrowheads for the 20 mol% TOCL high salt sample. Note that the low- and high-salt time courses for peptide addition in the absence of membrane ($[\text{lipid}]^{tot}$ = 0, black traces) are identical, but are shown in each panel for comparison.

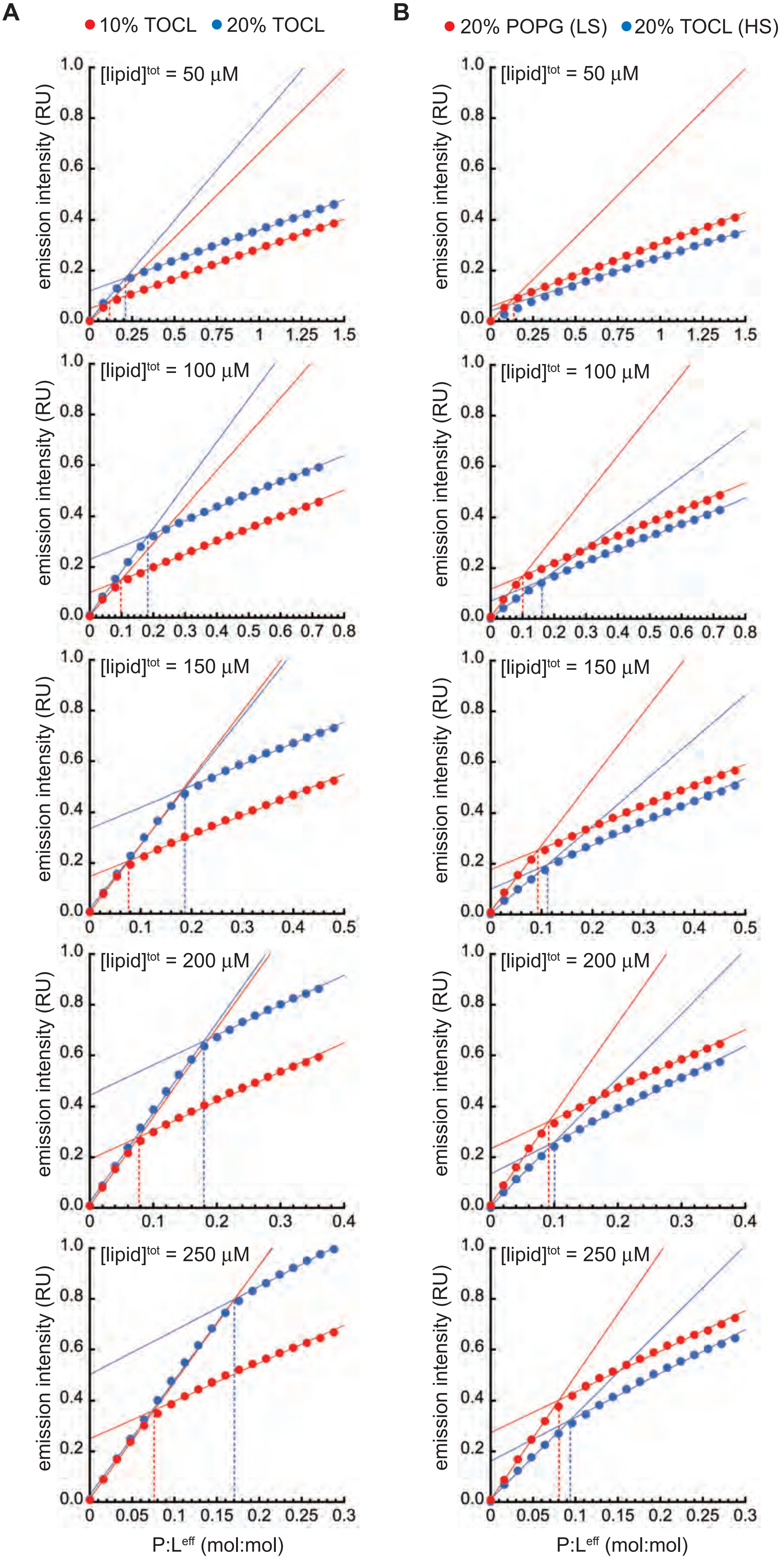


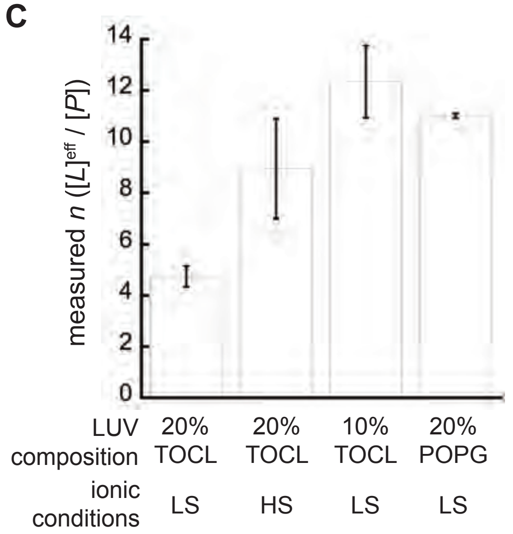


**Supplementary Figure S7. Analysis of time course binding saturation measurements.**
SS-31 fluorescence as a function of $[P]:[L]^{eff}$ (mol:mol) from peptide titrations into solutions of LUVs at the total lipid concentration indicated. Calculated inflection points for each titration delineate pre- and post-saturation stages of the titration, with each stage fit to a linear function shown by lines with cognate colors. Dashed drop lines to the abscissa mark the $P:L^{eff}$ at the inflection point for each titration.
**A.** SS-31 titrations of LUVs composed of 10% TOCL (red) or 20% TOCL (blue) under low salt conditions.
**B.** SS-31 titrations of LUVs composed of 20% POPG under low salt conditions (red) or 20% TOCL under high salt conditions (blue).
**C.** Measured values of $[L]^{eff}:[P]$ (=*n*) for each LUV type and salt condition shown in panels A and B (means +/- SD).

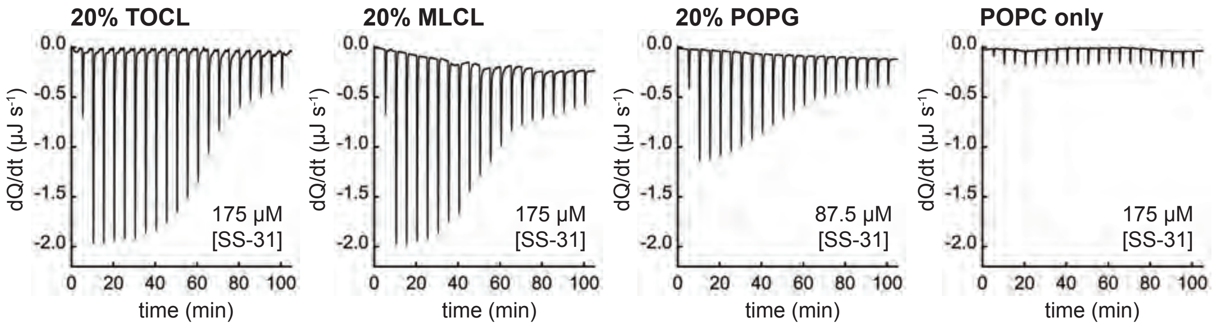


**Supplementary Figure S8. Microcalorimetry analysis of SS-31 membrane binding**

Representative ITC raw data (heat flow time courses) obtained by titration of SS-31 at the indicated concentrations with LUVs of different lipid composition. Downward peaks represent exothermic events from injections of LUVs (10 nmol lipid each).

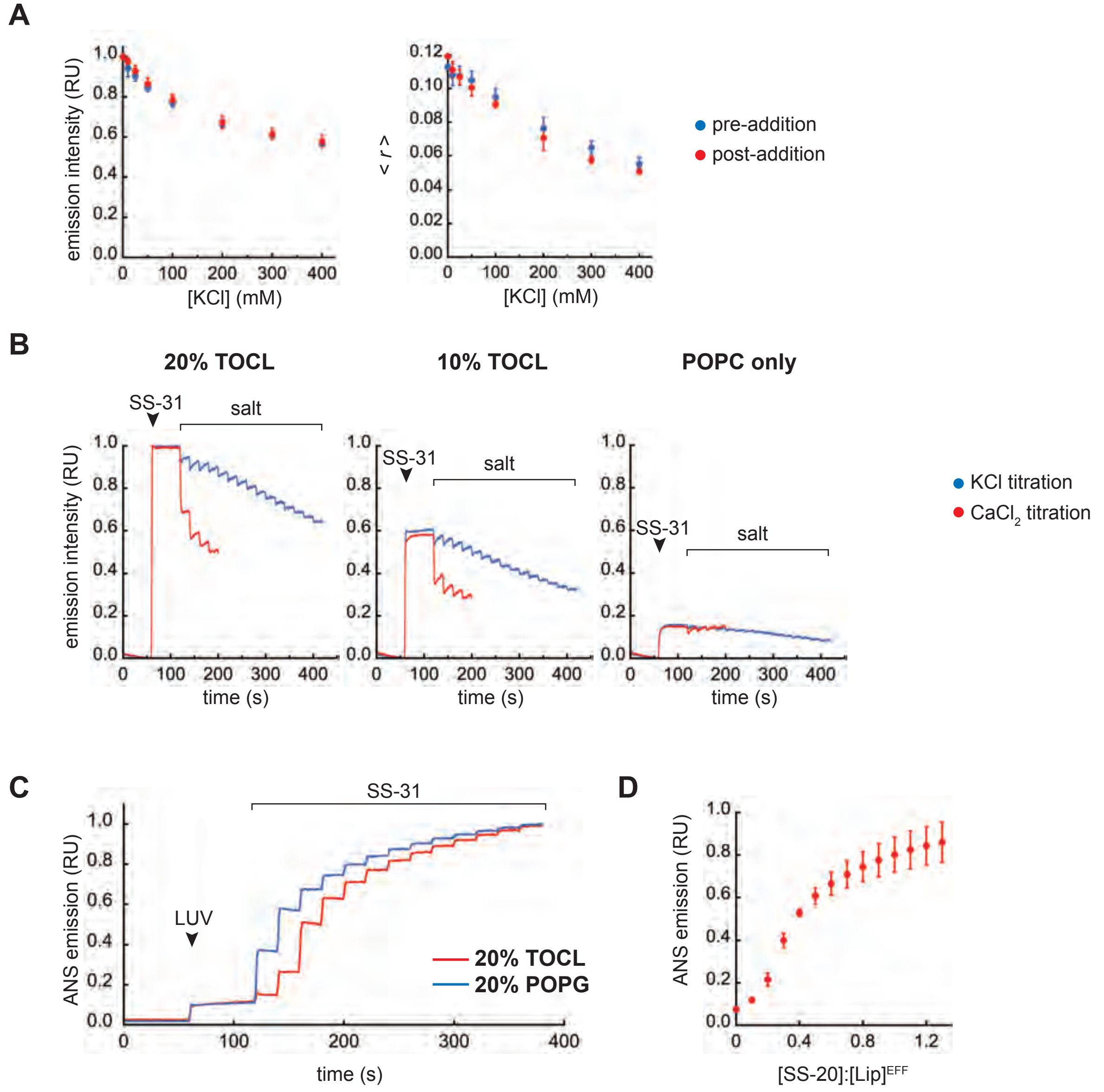


**Supplementary Figure S9. Relationship between SS peptide binding, ionic strength, and surface potential**

**A. Effect of pre- and post-salt addition.** Different concentrations of KCl were added to LUVs (20 mol% TOCL) before binding of SS-31 (pre-addition, blue) or after the binding of SS-31 (post-addition, red) followed by measurements of emission intensity (*left*) or steady state anisotropy (*right*) of SS-31.

**B. Time courses of ionic strength titration**. Representative time courses of endogenous SS-31 emission in the presence of LUVs with progressive addition of salts. At t=60 s (black arrowheads), SS-31 was added to suspensions of LUVs (20 mol% TOCL, 10 mol% TOCL, or POPC only). Titration of salts (starting at t=120 s) proceeded by addition of KCl (15 µmol increments, blue traces) or $CaCl_2$ (0.5 µmol increments, red traces) at 20 s intervals. $Ca^{2+}$ titrations were truncated prior to the point of calcium-induced lipid aggregation (determined from Fig. S4C).

**C. Time courses of 1,8-ANS emission with SS-31 titration.** Representative time courses of 1,8-ANS emission with SS-31 addition to suspensions of LUVs. At t=60 s (black arrowhead), LUVs composed of 20 mol% TOCL (red trace) or 20 mol% POPG (blue trace) were added to measurement buffer ([1,8-ANS] = 1 µM) at a concentration of $[Lip]^{eff}$ = 50 µM. Titration of SS-31 (starting at t=120 s) proceeded by peptide addition (10 nmol increments) at 20 s intervals.

**D. Effect of SS-20 on surface potential.** Measurements of ANS fluorescence are shown as a function of [SS-20]:$[Lip]^{eff}$ for membranes containing 20% TOCL (values are means, n=4 ± SD).

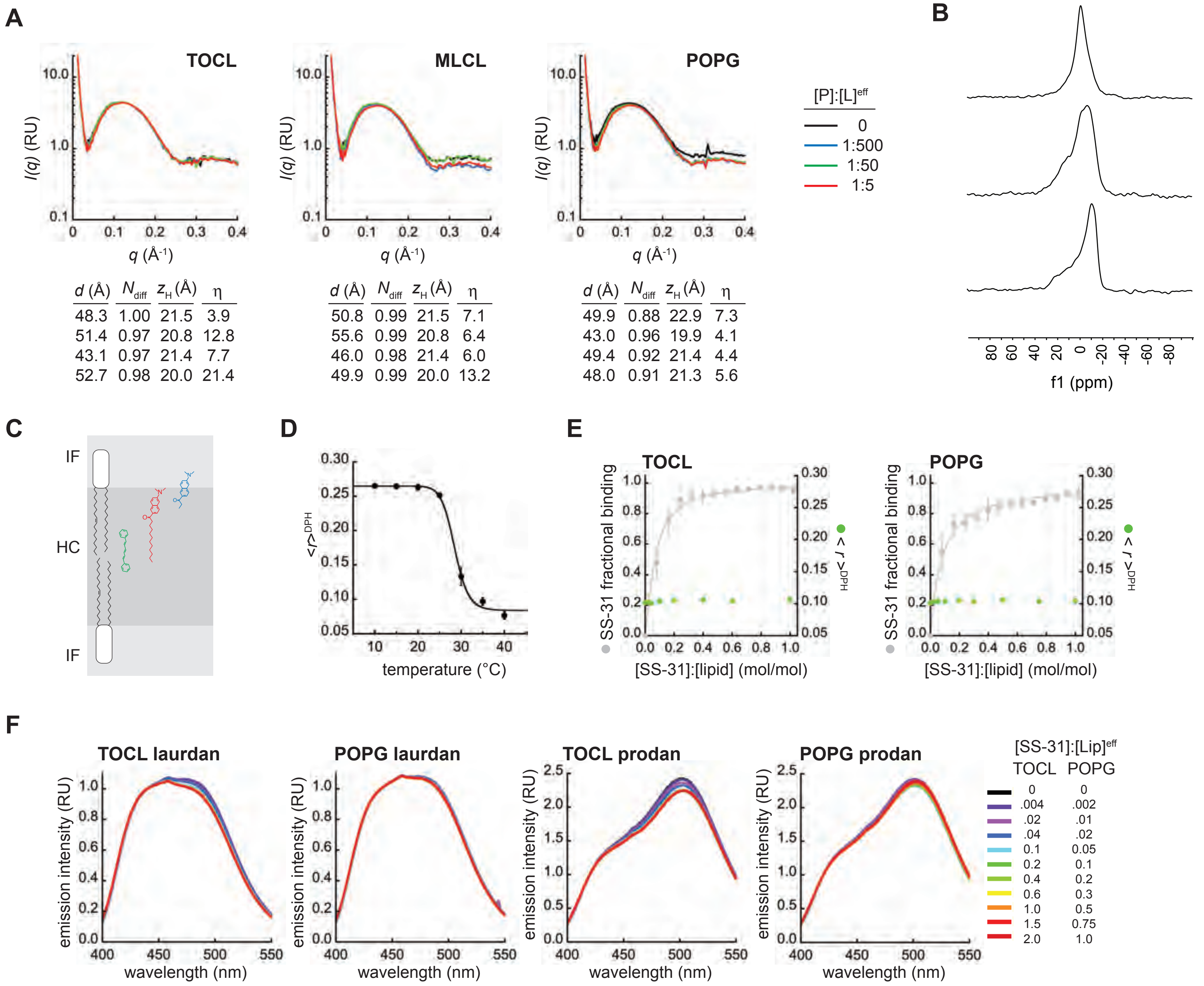


TOCL

| d (Å) | $N_{diff}$ | $z_H$ (Å) | η |
|---|---|---|---|
| 48.3 | 1.00 | 21.5 | 3.9 |
| 51.4 | 0.97 | 20.8 | 12.8 |
| 43.1 | 0.97 | 21.4 | 7.7 |
| 52.7 | 0.98 | 20.0 | 21.4 |

MLCL

| d (Å) | $N_{diff}$ | $z_H$ (Å) | η |
|---|---|---|---|
| 50.8 | 0.99 | 21.5 | 7.1 |
| 55.6 | 0.99 | 20.8 | 6.4 |
| 46.0 | 0.98 | 21.4 | 6.0 |
| 49.9 | 0.99 | 20.0 | 13.2 |

POPG

| d (Å) | $N_{diff}$ | $z_H$ (Å) | η |
|---|---|---|---|
| 49.9 | 0.88 | 22.9 | 7.3 |
| 43.0 | 0.96 | 19.9 | 4.1 |
| 49.4 | 0.92 | 21.4 | 4.4 |
| 48.0 | 0.91 | 21.3 | 5.6 |

**Supplementary Figure S10. Characterization of the effects of SS-31 on model membranes.**
**A. Synchrotron SAXS.** Background-corrected scattering density profiles are shown for LUVs containing anionic lipid (20% TOCL, MLCL, or POPG) or POPC only with the SS-31:lipid molar ratios indicated. Values reported in the table below are taken from fits of the data to Eq. S13 using the Global Analysis Program (GAP).
**B. $^{31}$P ssNMR.** $^{31}$P NMR spectra of lipids (80 mol% POPC, 20 mol% TOCL) prepared as: (i) MLVs (*lower trace*), (ii) MLVs with freeze/thaw cycles (*middle trace*), or (iii) 100 nm LUVs made by extrusion (*upper trace*).
**C. Membrane probes.** Approximate positions of fluorescent reporters DPH (green), laurdan (red) and prodan (blue) relative to the interfacial region (IF) and hydrocarbon core (HC) of the lipid bilayer.
**D-E. Anisotropy measurements of DPH-containing membranes.** (*D*) LUVs containing 80 mol% DMPC and 20 mol% TMCL with DPH were incubated in temperatures at 5°C increments and measured for steady-state anisotropy of DPH ($<r>^{DPH}$, each point n=3 ± SD). Fit to a sigmoidal function (black line) yields a maximal $<r>^{DPH}$ of 0.265 (corresponding to the Lβ phase), a minimal $<r>^{DPH}$ of 0.084 (corresponding to the Lα phase), and an inflection point at 28.5°C. (*E*) LUVs containing 80 mol% POPC and 20 mol% TOCL or POPG as indicated were incubated with increasing [SS-31] and measured for $<r>^{DPH}$ (n=3 ± SD). Data and fits in gray show SS-31 fractional saturation (calculated from Fig. 2A, 25 µM lipid).
**F. Emission scans of laurdan- and prodan-containing membranes.** LUVs composed of 20% TOCL or 20% POPG, containing laurdan or prodan as indicated, were incubated in the presence of varying $[SS\text{-}31]:[Lip]^{eff}$ as shown. Scans are signal averages (n=3) of background-subtracted spectra, normalized relative to $\lambda_{em}$=440 nm (laurdan) or $\lambda_{em}$ = 420 nm (prodan) to facilitate visualization.

A

| 20 mol% TOCL | w/ SS31 | No SS31 |
|---|---|---|
| TOCL P1 | 1.92 | 1.97 |
| | 1.89 - 1.94 | 1.97 - 1.98 |
| TOCL P3 | 1.92 | 1.96 |
| | 1.89 - 1.94 | 1.95 - 1.97 |
| POPC P | 2.00 | 2.00 |
| | 1.99 - 2.01 | 1.99 - 2.01 |

| 20 mol% MLCL | w/ SS31 | No SS31 |
|---|---|---|
| MLCL P1 | 1.90 | 1.93 |
| | 1.88 - 1.91 | 1.92 - 1.94 |
| MLCL P3 | 1.97 | 2.02 |
| | 1. 95 - 1.99 | 2.01 - 2.03 |
| POPC P | 1.94 | 1.93 |
| | 1.93 - 1.95 | 1.93 - 1.94 |

| 20 mol% POPG | w/ SS31 | No SS31 |
|---|---|---|
| POPG P | 1.88 | 1.92 |
| | 1.86 - 1.89 | 1.92 - 1.93 |
| POPC P | 1.95 | 1.95 |
| | 1.94 - 1.95 | 1.95 - 1.96 |

B

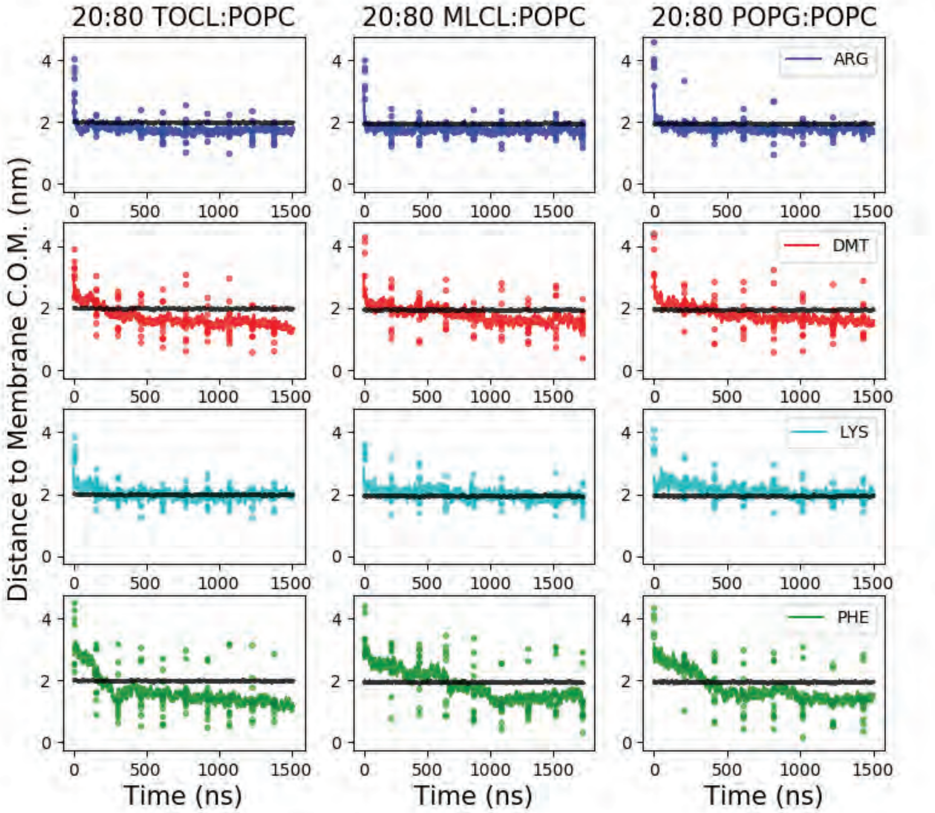


C

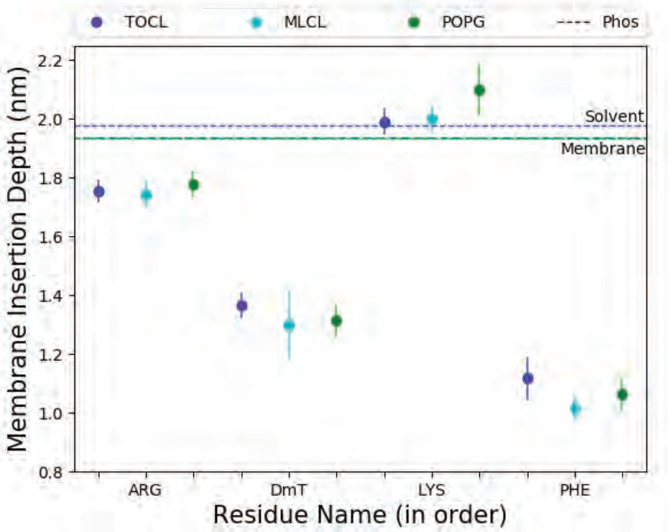


| SS-31 Landmark Atoms | 20:80 TOCL:POPC | | 20:80 MLCL:POPC | | 20:80 POPG:POPC | |
|---|---|---|---|---|---|---|
| | $Z^{pos}$ (nm) | 95% CI | $Z^{pos}$ (nm) | 95% CI | $Z^{pos}$ (nm) | 95% CI |
| Arg, $C_\zeta$ | 1.75 | 1.71 - 1.79 | 1.74 | 1.70 - 1.79 | 1.78 | 1.74 - 1.82 |
| DmT, $O_\eta$ | 1.37 | 1.32 - 1.41 | 1.30 | 1.18 - 1.42 | 1.31 | 1.26 - 1.37 |
| Lys, $N_\zeta$ | 1.99 | 1.95 - 2.03 | 2.00 | 1.95 - 2.05 | 2.10 | 2.01 - 2.19 |
| Phe, $C_\zeta$ | 1.12 | 1.04 - 1.19 | 1.02 | 0.98 - 1.06 | 1.06 | 1.00 - 1.12 |

**Supplementary Figure S11. MD simulations: lipid phosphate positions and SS-31 side chain insertion depths.**

**A. Lipid-specific phosphate z-positions** normalized to the bilayer center of mass (C.O.M.). Values shown for each lipid and phosphate type are means in nm (*upper box*) ± 95% CI (*lower box*).

**B. Time course profiles** are shown for individual SS-31 side chain positions relative to bilayers composed of the indicated lipids. Positions along the z-axis were determined as the distance between each side chain and the calculated C.O.M. of the membrane. The black line shows the average position of all phosphates in the upper leaflet. Colored dots show the z coordinates ($z^{pos}$) of individual side chains, and colored lines represent the average z coordinates.

**C. Comparison of membrane insertion depths** from MD simulations for the "peptide burial state". *Left,* average membrane insertion depths ($z^{pos}$, n=10 ± 95% CI) for landmark atoms of each side chain of SS-31 in bilayers of different lipid compositions indicated in comparison with average $z^{pos}$ levels of headgroup phosphates. *Right*, table showing values of $z^{pos}$ of SS-31 side chains. Each residue assumed a distinct membrane insertion depth with no significant difference observed among the different lipid systems.

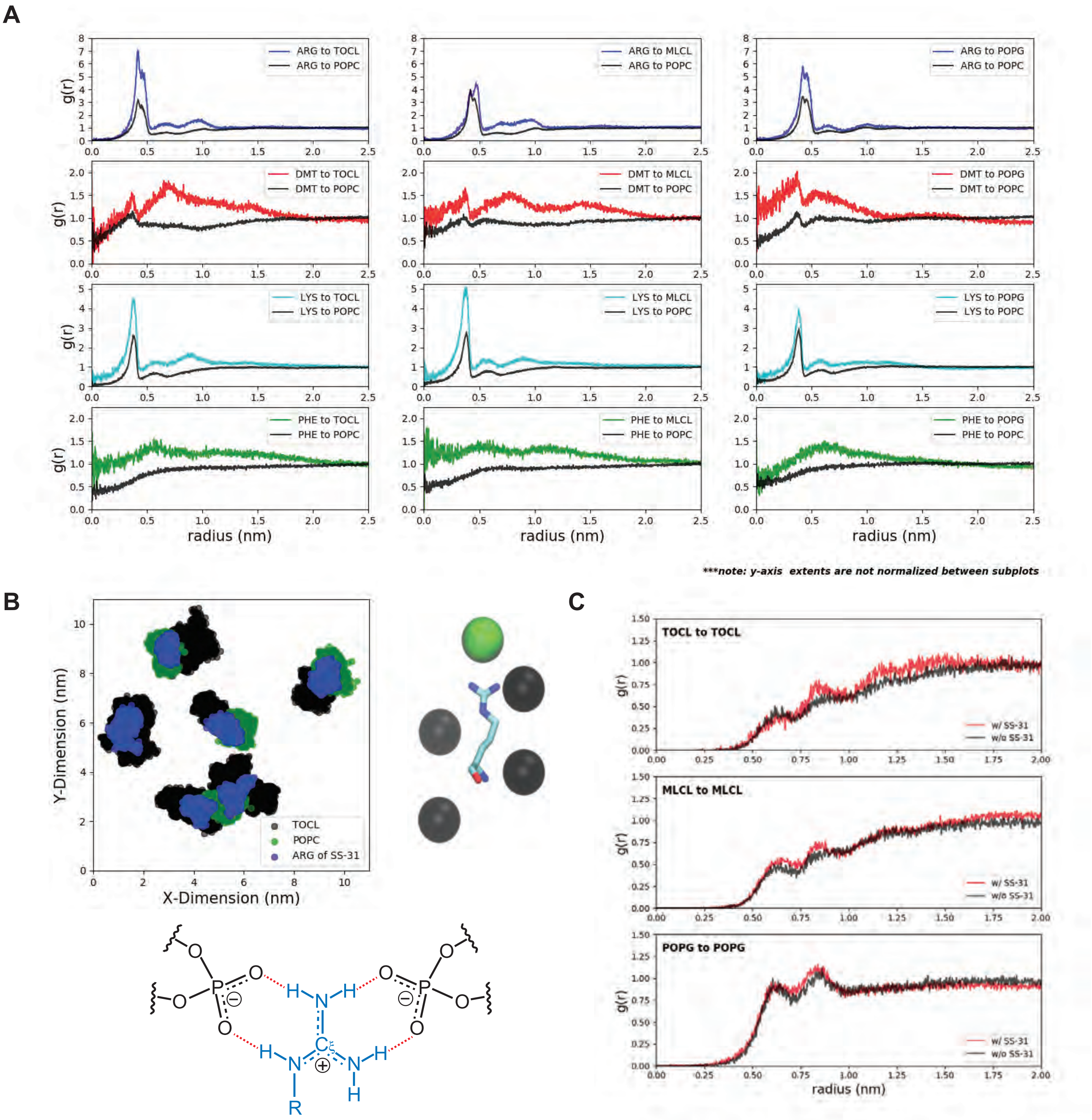


**Supplementary Figure S12. Lipid radial distribution profiles from MD simulations**

**A. Lipid distributions around SS-31 residues.** Lateral (x-y plane) radial distribution profiles from side chain landmark atoms (the D-Arg Cζ atom; the 2´,6´-Dmt Oη atom; the Lys Nζ atom; or the Phe Cζ atom) to lipid headgroup phosphate atoms (TOCL and MLCL $P_1$, POPG and POPC, P). Note that plots have different y-axis scales to aid in distinguishing between curves. The landmark atoms on the aromatic residues (DmT, Phe) can overlap with the phosphates by being positioned above or below the headgroups in the Z-direction, leading to elevated concentrations below the van der Waals radius.

**B. Coordination between SS-31 side chains and lipid phosphates.** *Upper left,* top-down view of the lipid bilayer (in x-y space) showing the diffusion range of Arg (central Cζ atom, blue) over a 100 ns range within a typical simulation. POPC phosphates (green) and TOCL phosphates (black) that were proximal to the Arg over the time course are shown. *Upper right*, snapshot of typical orientation between Arg (ball and stick representation) and headgroup phosphates. *Lower*, schematic of bidentate Arg-phosphate complex stabilized by multiple hydrogen bonding interactions.

**C. Lipid-to-lipid concentration with and without SS-31 from MD simulations.** Radial distribution profiles of phosphate to phosphate in each lipid headgroup (TOCL and MLCL $P_1$, POPG, P) in the upper leaflet of systems with SS-31 (red) and bilayers of systems without SS-31 (black).

**A**

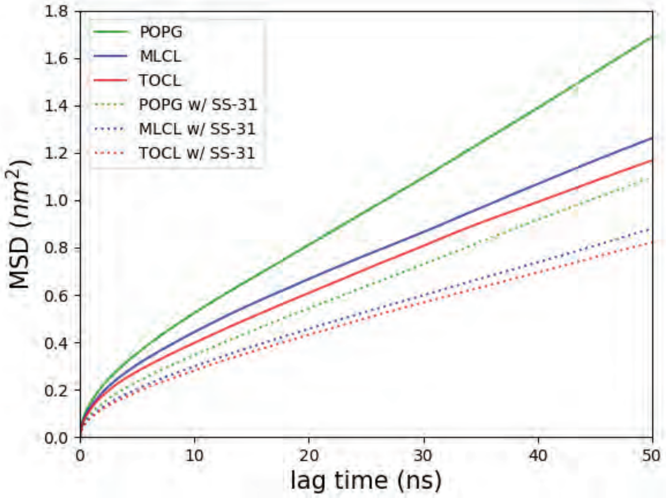


**B**

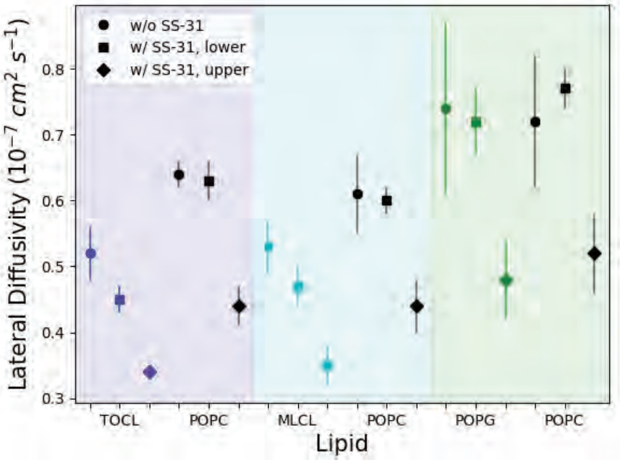


| Lipid System | 20:80 TOCL:POPC | | 20:80 MLCL:POPC | | 20:80 POPG:POPC | |
|---|---|---|---|---|---|---|
| | $D_{xy}$ ($10^{-7}$ $cm^2$ $s^{-1}$) | 95% Conf. Interval | $D_{xy}$ ($10^{-7}$ $cm^2$ $s^{-1}$) | 95% Conf. Interval | $D_{xy}$ ($10^{-7}$ $cm^2$ $s^{-1}$) | 95% Conf. Interval |
| Anionic Lipid w/ SS-31, upper | 0.34 | 0.33 - 0.34 | 0.35 | 0.32 - 0.38 | 0.48 | 0.42 - 0.54 |
| Anionic Lipid w/ SS-31, lower | 0.45 | 0.43 - 0.47 | 0.47 | 0.44 - 0.51 | 0.72 | 0.67 - 0.77 |
| Anionic Lipid | 0.52 | 0.48 - 0.55 | 0.53 | 0.49 - 0.57 | 0.74 | 0.61 - 0.87 |
| POPC w/ SS-31, upper | 0.44 | 0.42 - 0.47 | 0.44 | 0.41 - 0.48 | 0.52 | 0.46 - 0.58 |
| POPC w/ SS-31, lower | 0.63 | 0.60 - 0.66 | 0.60 | 0.58 - 0.62 | 0.77 | 0.74 - 0.80 |
| POPC | 0.64 | 0.62 - 0.66 | 0.61 | 0.55 - 0.66 | 0.72 | 0.62 - 0.82 |

**Supplementary Figure S13. Mean square displacement of lipids with and without SS-31 from MD simulations.**

**A. Representative mean square displacement analysis** in the lateral (xy) plane of phosphates from headgroups from anionic lipids.

**B. Lateral (xy) lipid diffusion coefficients for bilayers with and without SS-31.** Diffusion constants were calculated from mean square displacement analyses of each lipid's headgroup phosphate(s). The figure (*left*) and table (*right)* includes diffusion constants ($D_{xy}$) for lipid systems without any SS-31 at all, as well as separate diffusion constants for the upper (peptide-accessible) and lower (opposing) leaflets.

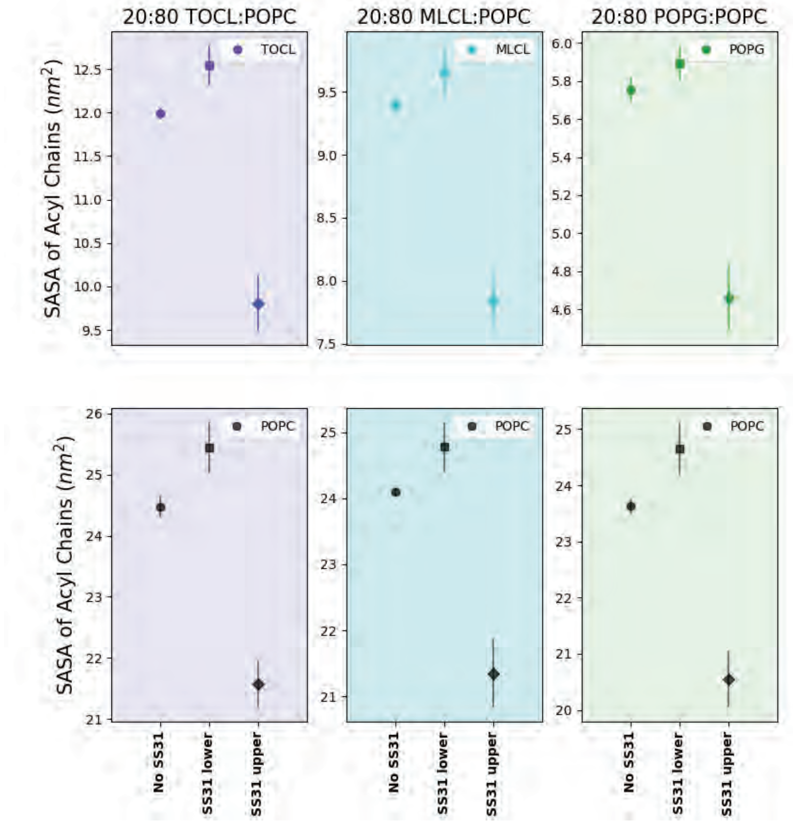


<table>
<tr><th rowspan="2">Lipid System</th><th colspan="2">20:80 TOCL:POPC</th><th colspan="2">20:80 MLCL:POPC</th><th colspan="2">20:80 POPG:POPC</th></tr>
<tr><th>Acyl Chain SASA ($nm^2$)</th><th>95% Conf. Interval</th><th>Acyl Chain SASA ($nm^2$)</th><th>95% Conf. Interval</th><th>Acyl Chain SASA ($nm^2$)</th><th>95% Conf. Interval</th></tr>
<tr><td>Anionic Lipid w/ SS-31, upper</td><td>9.8</td><td>9.5 - 10.1</td><td>7.8</td><td>7.6 - 8.1</td><td>4.7</td><td>4.5 - 4.8</td></tr>
<tr><td>Anionic Lipid w/ SS-31, lower</td><td>12.5</td><td>12.3 - 12.8</td><td>9.7</td><td>9.4 - 9.9</td><td>5.9</td><td>5.8 - 6.0</td></tr>
<tr><td>Anionic Lipid</td><td>12.0</td><td>11.9 - 12.1</td><td>9.4</td><td>9.3 - 9.5</td><td>5.8</td><td>5.7 - 5.9</td></tr>
<tr><td>POPC w/ SS-31, upper</td><td>21.6</td><td>21.2 - 22.0</td><td>21.3</td><td>20.8 - 21.9</td><td>20.5</td><td>20.0 - 21.0</td></tr>
<tr><td>POPC w/ SS-31, lower</td><td>25.4</td><td>25.0 - 25.9</td><td>24.8</td><td>24.4 - 25.1</td><td>24.7</td><td>24.2 - 25.1</td></tr>
<tr><td>POPC</td><td>24.5</td><td>24.3 - 24.6</td><td>24.1</td><td>24.0 - 24.2</td><td>23.6</td><td>23.5 - 23.8</td></tr>
</table>

**Supplementary Figure S14. Acyl chain solvent accessible surface area from MD simulations**
The average SASA of each lipid acyl chain region was calculated for all tested lipids with and without SS-31. Confidence intervals (95%) were calculated from the standard error of the mean from three equal regions of each trajectory. The figure (*left*) and table (*right*) include SASA measurements for lipid systems in the absence of SS-31 as well as separate SASA measurements for the upper (peptide-exposed) and lower (lacking peptide) leaflets of systems with SS-31. The presence of SS-31 in the upper leaflet decreased SASA of the acyl chain region for all tested lipids compared to both a) the lower leaflet of the same SS-31 containing system, and b) bilayer systems without SS-31. The presence of SS-31 in the upper leaflet also increased the SASA of the lower (opposing) leaflet compared with systems lacking SS-31 for specific lipids, including TOCL and POPC in the 20:80 TOCL:POPC system, POPC in the 20:80 MLCL:POPC system, and POPC in the 20:80 POPG:POPC system.

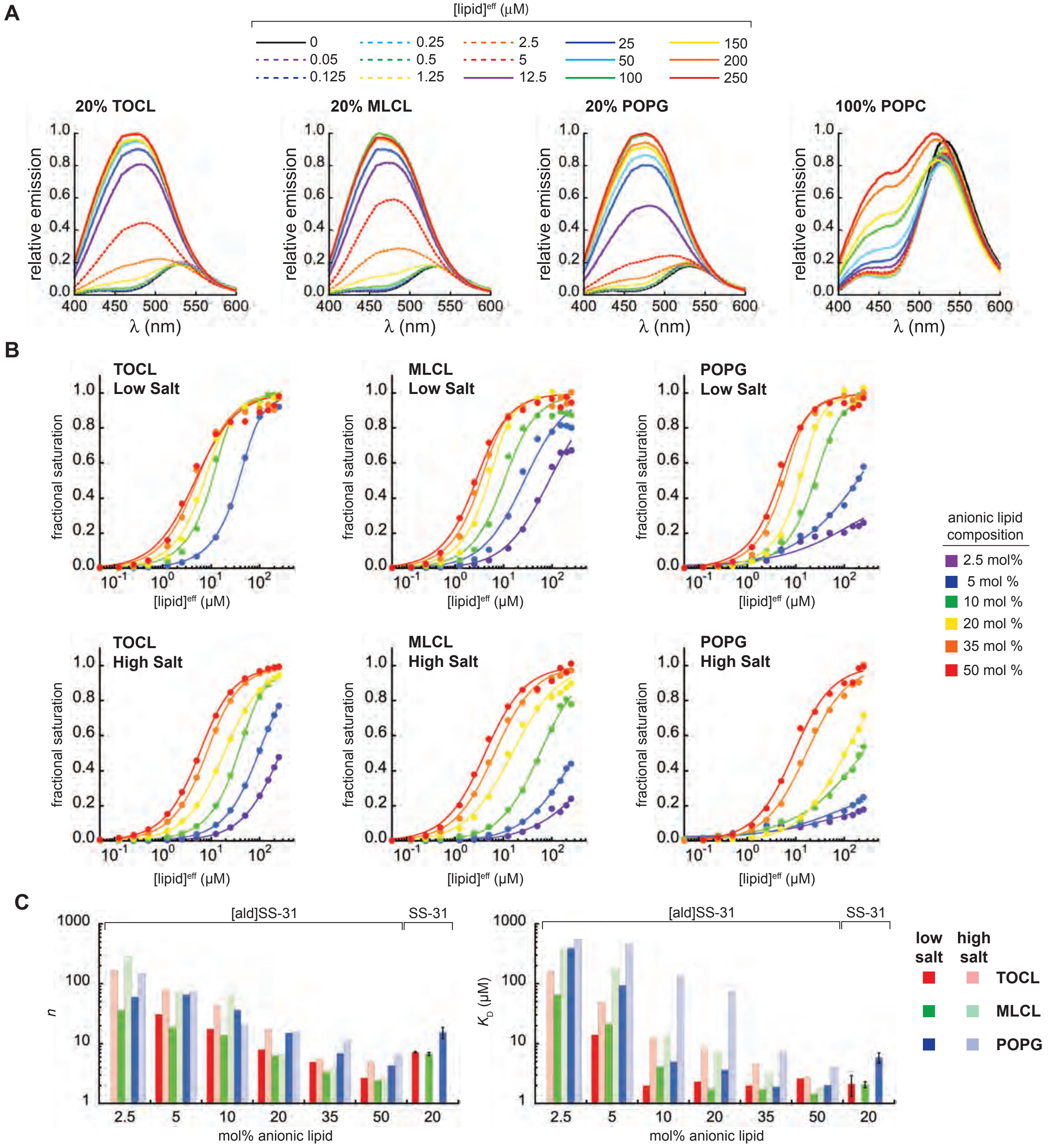


**Supplementary Figure S15. [ald]SS-31 binding isotherms with model membranes**

**A. [ald]SS-31 fluorescence scans.** Emission spectra ($\lambda_{ex}$ = 360 nm; $\lambda_{em}$ = 400-600 nm) of 1 µM [ald]SS-31 in the presence of increasing concentrations of LUVs containing 20 mol% anionic lipid (TOCL, MLCL, or POPG) in a POPC background, or 100% POPC as indicated.

**B. [ald]SS-31 binding isotherm composite.** Binding curves for 1 µM [ald]SS-31 with liposomes containing anionic lipids (TOCL, MLCL, or POPG) at the concentrations indicated with a background of POPC. Measurements were made in the presence of low salt (upper panels) or high salt (lower panels).

**C. Equilibrium binding parameters.** Values under "[ald]SS-31" show calculated parameters *n* and $K_D$ from lipid titration isotherms shown in panel B, fit according to Eq. S8 for each sample (LUVs containing the indicated mol% of TOCL, MLCL, or POPG under high and low salt conditions). For comparison, values under "SS-31" show *n* and $K_D$ values based on binding curves from endogenous SS-31 fluorescence (Fig. 2B).

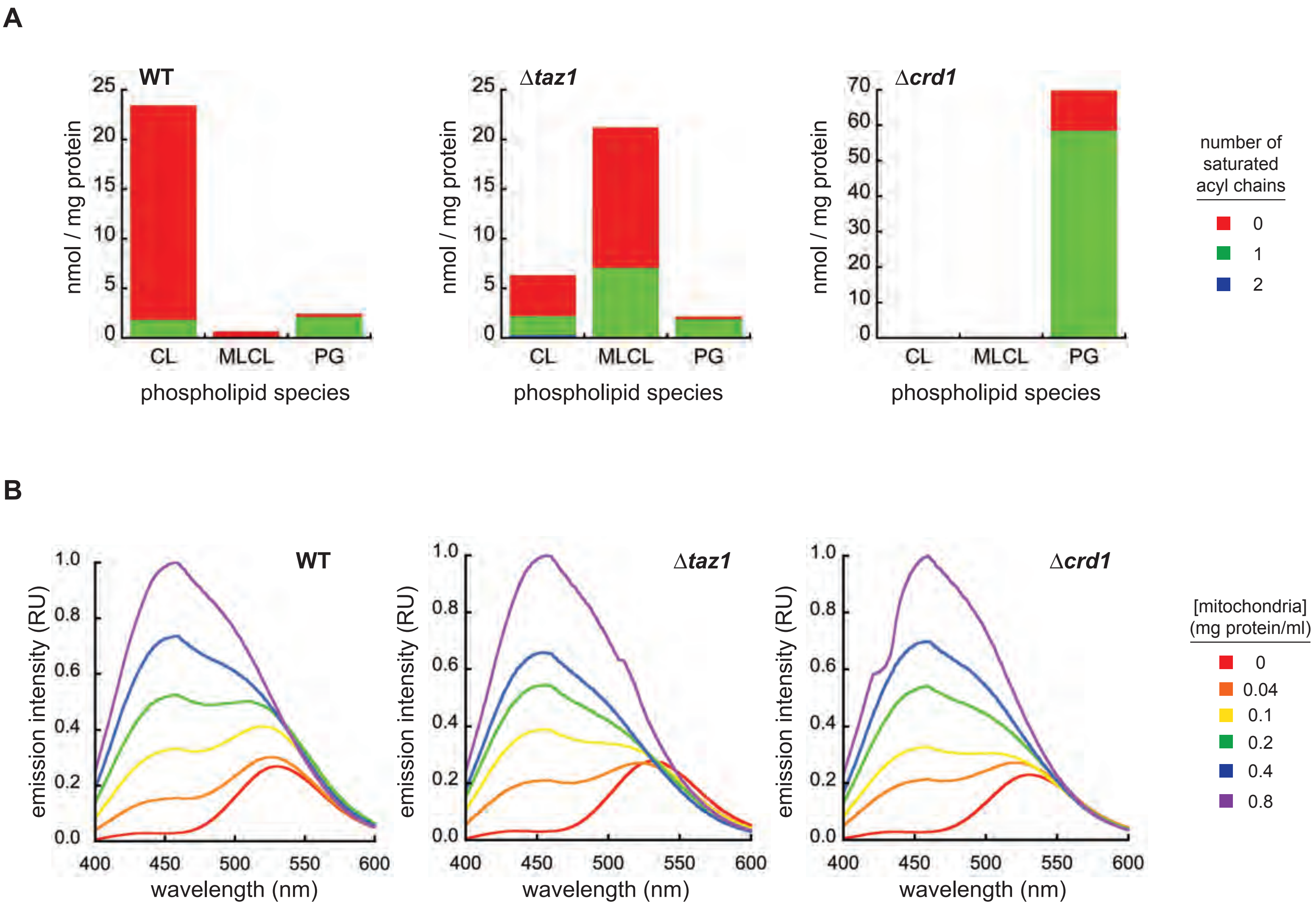


**Supplementary Figure S16. Interaction of SS-31 with isolated mitochondria**

**A. Acyl chain distribution** of anionic phospholipids in mitochondria from different yeast strains. Based on lipidomics analysis of mitochondria from WT, Δ*taz1*, and Δ*crd1* yeast strains, anionic phospholipids CL, MLCL and PG were classified based on chain saturation as indicated (quantified as nmol lipid per mg mitochondrial protein). Shown are the means of n=3 individual samples per strain.

**B. Representative background-subtracted and normalized emission spectra** containing 1 μM [ald]SS-31 with the indicated concentration of isolated mitochondria.

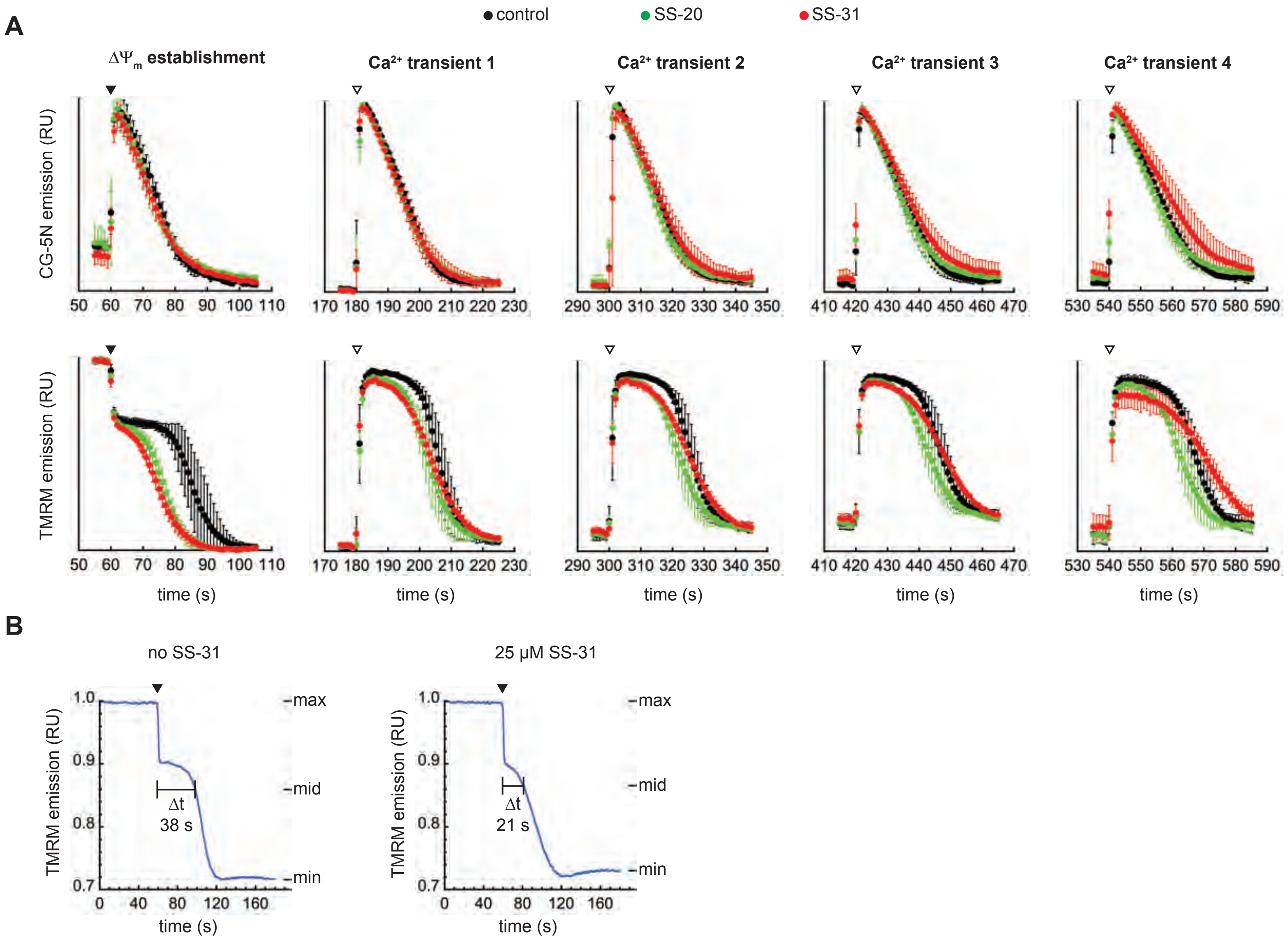


**Supplementary Figure 18. Kinetic responses of $\Delta\Psi_m$ and external [$Ca^{2+}$]**

**A. Expanded views of $\Delta\Psi_m$ establishment and $Ca^{2+}$ transients** are shown for CG-5N measurements (*upper panels*) and TMRM measurements (*lower panels*). Data represent individual traces (n=3 ± SD) from Fig. 7E from samples in the presence of ETH-129 and in the absence of SS peptide (control), or in the presence of 25 µM SS-20 or SS-31 as indicated.

**B. Quantitative analysis of $\Delta\psi_m$ establishment** for the dose response curve shown in Fig. 7G. TMRM traces of mitochondria in the presence of 25 µM $CaCl_2$ and ETH-129 were measured over time courses in which respiratory substrate (1 mM NADH) was added at t=60 s (black arrowhead) and the $\Delta\psi_m$ was established over an additional 120 s. From the kinetic traces of TMRM emission, three points were measured: max (average of points prior to NADH addition), min (average of last 20 s of traces) and mid (average of max and min, representing the midpoint of the TMRM emission range). As a quantitative measure of the $Ca^{2+}$-dependent temporal delay in $\Delta\psi_m$ establishment, we calculated $\Delta t$ as the time required to reach the mid TMRM value following respiratory substrate addition.

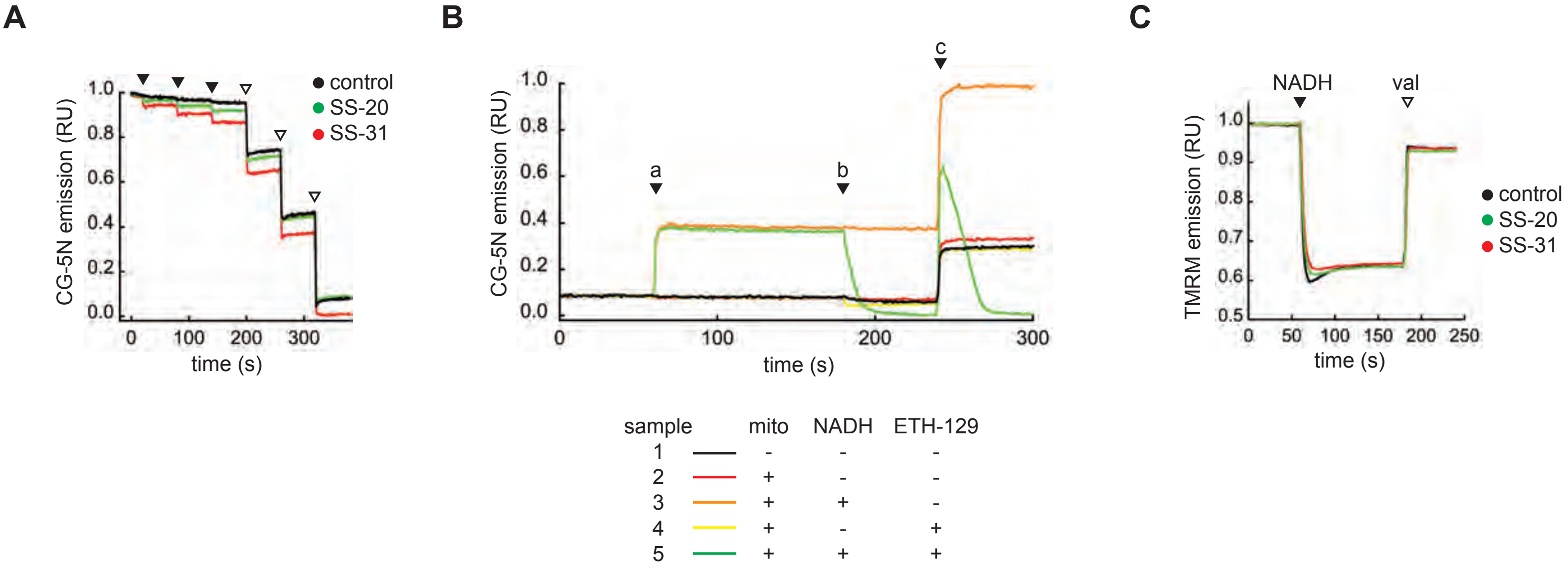


**Supplementary Figure 17. Fluorescence-based assays of calcium dynamics and $\Delta\psi_m$**

**A. Effect of SS peptides on CG-5N fluorescence under conditions used to measure model membranes.** Samples containing minimal LUV buffer (20 mM HEPES, pH 7.5) in the presence of 1 μM CG-5N and supplemented with 2.5 μM $CaCl_2$ were tested for the effects of SS peptide on probe emission. Time course measurements show three additions of peptide (2.5 nmol each addition) or vehicle only (black arrowheads) followed by three additions of EDTA, pH 7.5 (1 nmol each addition, open arrowheads).

**B. Calcium concentration time courses.** Time course measurements of CG-5N emission with samples containing mitochondria measurement buffer (20 mM HEPES, 300 mM sucrose, 2 mM potassium phosphate, 0.05% BSA, pH 7.5) containing 1 μM CG-5N with or without mitochondria (50 μg $ml^{-1}$) as indicated. At the indicated time points (black arrowheads), the following additions were made: a) 1 mM NADH or buffer only; b) 5 μM ETH-129 or DMSO only; c) 25 μM $CaCl_2$. Emission data were normalized relative to values just before addition at point a.

**C. Membrane potential time courses.** Time course measurements of TMRM emission with samples identical to those in panel B, but containing 50 nM TMRM, with mitochondria in the absence of peptide (control) or preincubated with 25 μM SS-31 or SS-20 as indicated. NADH (1 mM) and valinomycin (1 μM) were added at time points indicated by black and open arrowheads, respectively. Emission data were normalized relative to values just prior to NADH addition.

A

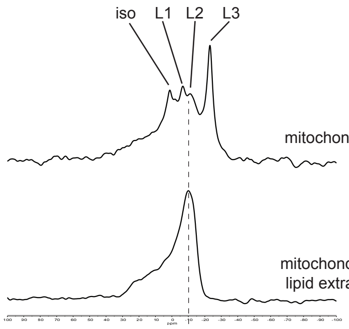


B

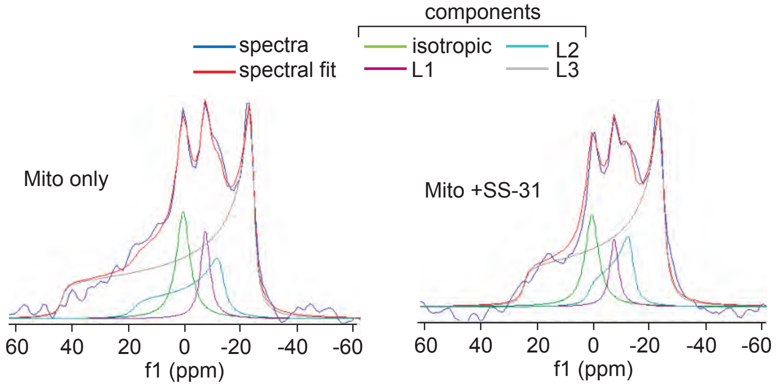


C

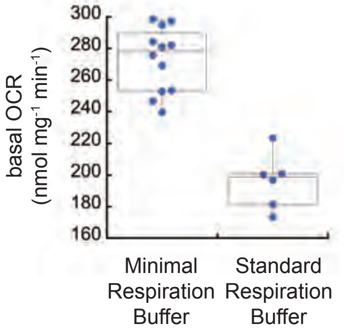


**Supplementary Figure S19. NMR and respirometry measurements with calcium stress**

**A,B. $^{31}$P ssNMR measurements.** (*A*) Static wide-line $^{31}$P NMR spectra of isolated mitochondria (*upper panel*) and MLVs prepared from extracted mitochondrial lipids (*lower panel*). (*B*) Spectra of isolated mitochondria in the absence (*left*) or presence (*right*) of SS-31. Based on spectral deconvolution (dmFit), identified peaks from mitochondria samples include an isotropic component at 0 ppm ("Iso") and three high-field signals labeled "L1", "L2" and "L3" centered at -7, -12, and -23 ppm, respectively.

**C. Condition-dependent respirometry measurements.** Basal oxygen consumption rates are shown for measurement under Minimal Respiration Buffer (data from Fig. 7I) and Standard Respiration Buffer as indicated.
